# Peptide sequencing via reverse translation of peptides into DNA

**DOI:** 10.1101/2024.05.31.596913

**Authors:** Liwei Zheng, Yujia Sun, Michael Eisenstein, Hyongsok Tom Soh

## Abstract

Scalable methods that can accurately sequence peptides at single-amino acid resolution could significantly advance proteomic studies. We present a protein sequencing method based on the “reverse translation” of peptide sequence information into DNA barcodes that document the identity, position, and the originating peptide of each amino acid. We employ a modified Edman degradation process that converts peptides into DNA-barcoded amino acids, which are subsequently detected by proximity extension assay, yielding multi-barcoded DNA outputs that can be PCR amplified and sequenced. Using our method, we sequenced multiple consecutive amino acids within a model peptide. This method also enables the differentiation of single amino acid substitutions, and the identification of post-translational modifications and their positions within multiple peptides simultaneously. With further development, we anticipate that this method will enable highly parallel *de novo* protein sequencing with single-molecule sensitivity.

## Introduction

Single-cell transcriptomics has matured into a powerful method for directly quantifying the heterogeneous functional impact of genomic variations, environmental stimuli, and other physiological factors in different cell types and populations.(*1–3*) Although messenger RNA (mRNA) transcripts are a highly informative indicator of gene expression activity, the consequences of changes in expression can only be fully understood by assessing a cell’s protein content. This is because the final outcome of mRNA translation is heavily influenced by the effects of translational regulation and kinetics, post-translational modification (PTM), and other biochemical processes.(*4, 5*) Unfortunately, the toolbox for single-cell proteomics is still in its nascency. Advances in mass spectrometry (MS) instrumentation and techniques, such as multiplexed isobaric labeling, have enabled the quantification of ∼3,000 proteins from individual cells.(*6–9*) However, MS still lacks the dynamic range and sensitivity to document the full breadth of the cellular proteome.(*10–12*) This has spurred considerable interest in the development of technologies that enable single-molecule analysis and quantitation of proteins.

The past several years have witnessed considerable progress in this domain, including several early-stage commercial platforms.(*13*) One category of approaches emulates nanopore-based DNA sequencing, wherein nucleotide sequences are determined based on real-time changes in ionic current that occur as stretches of nucleic acids are threaded through protein nanopores. Researchers have made substantial headway in developing similarly premised platforms that can mediate the controlled transit of unfolded proteins through nanopores.(*14–18*) In addition, two recent publications have shown that such approaches can identify all 20 natural amino acids as well as certain PTMs in their monomeric forms or when attached to a constant carrier peptide.(*19, 20*) However, it remains unclear whether nanopore methods can consistently resolve individual amino acids in the context of a peptide,(*16, 18*) and true single-molecule protein sequencing with a nanopore-based system has yet to be demonstrated. Other methods employ an approach known as “fluorosequencing”, in which surface-conjugated proteins or peptides are selectively tagged with a range of amino acid-specific fluorescent labels.(*21*) In each round of sequencing, the proteins are imaged and then subjected to Edman degradation to iteratively trim away N-terminal amino acids. By matching labeled amino acids to specific positions on the peptide, one can identify many individual proteins in a highly parallel fashion. This approach is effective for high-level profiling of protein content, but fluorosequencing is inherently limited by the range of reagents available for selective amino acid labeling.(*22, 23*) Another recently developed approach exploits derivatives of the ClpS family proteins, which selectively recognize specific combinations of amino acids at the N-terminus of peptides.(*13*) Similar to fluorosequencing, the immobilized proteins are subjected to enzymatic N-terminal digestion to expose new amino acids for analysis. However, this approach is also limited in the range of amino acids that can be discriminated, particularly since each ClpS is also sensitive to the flanking residues in the peptide.(*24, 25*) As such, although significant progress is being made in terms of single-molecule protein profiling and quantitation, there is still an unmet need for methods that can accurately identify a wide range of proteins—including PTM variants—at a meaningful throughput.

In this work, we present a novel method that could potentially deliver such capabilities. Our process uses DNA-tagged, amino acid-specific antibodies to enable the “reverse translation” of peptide sequences into barcoded DNA libraries that encode the amino acid sequence. This is done in iterative cycles, where a modified Edman degradation process is used to iteratively remove amino acids from the N-terminus of an immobilized peptide.(*26*) During this process, each amino acid is tagged, before cleavage, with a DNA sequence that allows it to be traced back to the source peptide. The DNA-barcoded amino acids are subsequently detected with primer-tagged amino acid-specific antibodies, enabling PCR amplification of the barcode sequences. These can then be analyzed via high-throughput DNA sequencing methods, such that the resulting sequencing data reveal the identity, originating peptide, and position within that peptide for each amino acid. We initially demonstrate this method with a single seven-residue model peptide, and then explore the multiplexing potential of this approach by sequencing multiple peptide species simultaneously. We also demonstrate the feasibility of PTM mapping using antibodies that selectively bind and identify specific PTMs such as phosphotyrosine.

## Results and discussion

### Design of peptide sequencing via “reverse translation”

We designed a three-stage process to reverse translate peptide sequence information to DNA. The first stage entails the conversion of peptides into DNA-barcoded amino acids (**Fig. 1**, stage 1). To begin, the C-terminus of the peptides being sequenced are conjugated to beads modified with DNA-based peptide barcodes, which will subsequently be associated with a particular peptide. The immobilized DNA-barcoded peptides are then subjected to a modified version of the Edman degradation reaction. This reaction is a well-established tool for protein sequencing, where the N-terminal amino acid is conjugated to phenylisothiocyanate (PITC) and cleaved from the peptide under acidic conditions. The resulting degradation fragment is converted to a more stable form and identified, after which the process is repeated to identify the next amino acid. Reaction efficiency is generally not limited by amino acid identity or peptide charge states, and thus, Edman degradation can accommodate a wide range of peptide sequences.(*21*) In our particular implementation, an azide-modified PITC is used for N-terminus conjugation, which allows the click coupling of a dibenzocyclooctyne (DBCO)-modified biotinylated primer that hybridizes with the peptide barcode DNA. The primer is then immediately extended by DNA polymerase, after which the Edman reaction is performed to cleave the N-terminal amino acid. This yields a degradation product that is covalently tagged with the complementary sequence of the peptide barcode.

**Fig. 1.**
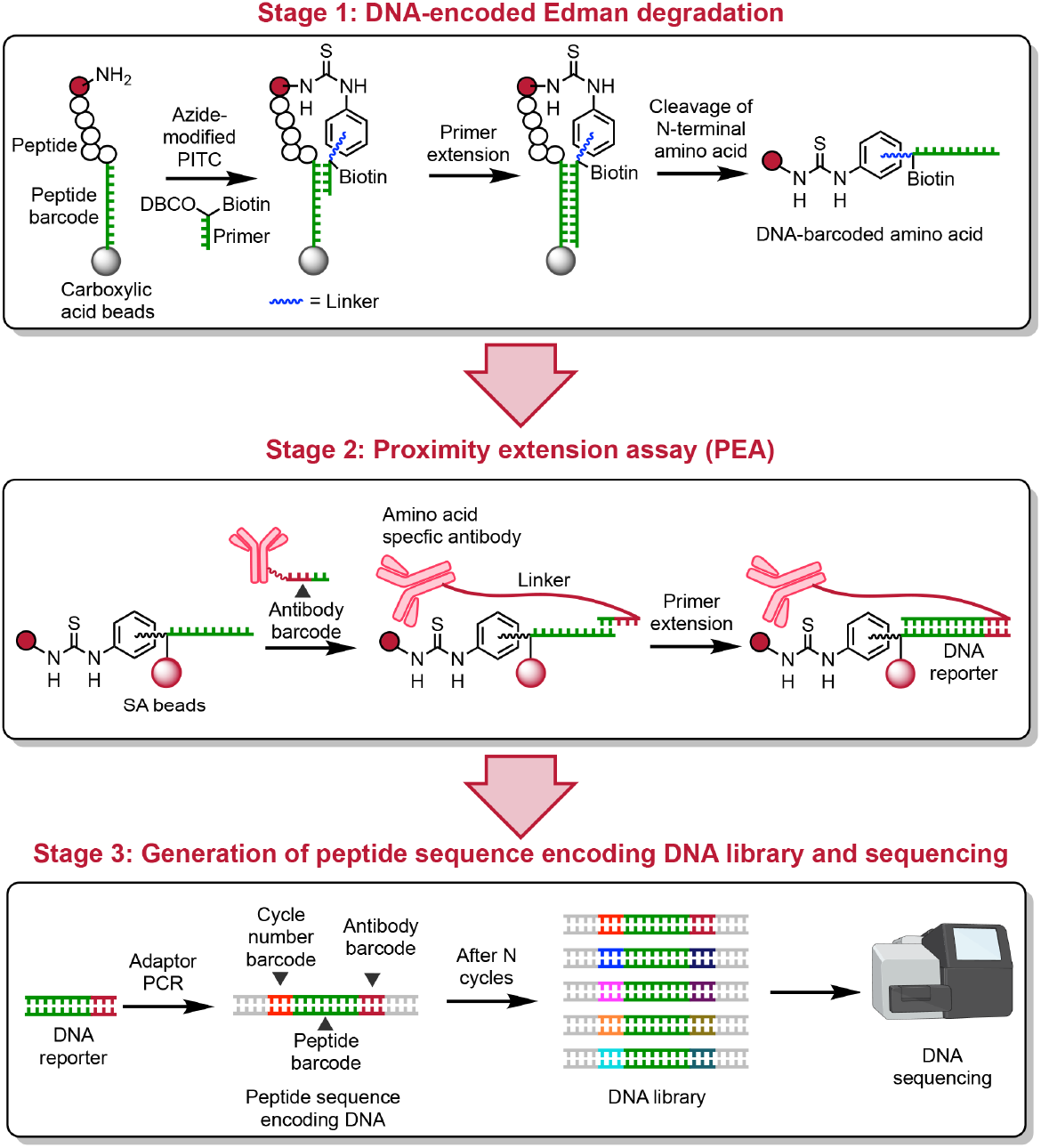
Overview of peptide sequencing via “reverse translation”. In stage 1, peptides are conjugated with peptide barcode DNA sequences immobilized on magnetic beads. A modified Edman degradation reaction is carried out using a PITC molecule modified with an azide group to enable click coupling to a DBCO-modified, biotinylated primer that hybridizes directly to the peptide barcode sequence. This primer is extended to record the peptide barcode before N-terminal amino acid cleavage. In stage 2, the DNA-barcoded amino acids released after cleavage are pulled down by streptavidin (SA) beads via their biotin moiety. These immobilized amino acids are recognized by DNA-barcoded amino acid-specific antibodies, and the DNA-barcoded amino acids are subsequently converted into DNA reporters via proximity extension assay (PEA). In stage 3, the DNA reporters from each cycle are barcoded with cycle number barcodes during adaptor PCR, yielding a DNA sequence that encodes the identity, position, and originating peptide of each amino acid. The final DNA library produced by this process can then be sequenced in a single run, enabling the reconstruction of peptide sequences from the sample.

In the second stage, the DNA-barcoded degradation products are converted to DNA reporters via proximity extension assay (PEA; **Fig. 1**, stage 2).(*27, 28*) The DNA-barcoded degradation products are enriched by streptavidin (SA) pulldown and subsequently recognized by DNA-barcoded amino acid-specific antibodies. The antibody-based identification of cleaved amino acids is more reliable compared to methods that attempt to detect N-terminal amino acids within the peptide context, since antibody binding will not be affected by neighboring amino acids. Binding events are recorded by PEA, where the resulting DNA reporters contain both the initial peptide-specific barcodes and the antibody-specific barcodes.

In the final stage, the DNA reporters generated by PEA from each cycle are tagged with cycle number barcodes during adaptor PCR before sequencing (**Fig. 1**, stage 3). This yields peptide sequence encoding DNA comprising a set of barcodes that specifically encode the identity, position, and originating peptide of each amino acid. This three-stage process is performed iteratively on the immobilized peptide, and the resulting DNA library is then sequenced, enabling the reconstruction of the amino acid sequence for each peptide present in the sample.

### DNA-compatible Edman degradation reaction

Preserving DNA integrity during Edman degradation is critical to our “reverse translation” strategy. However, conventional Edman degradation employs harsh conditions that damage DNA (**Fig. 2A**).(*29, 30*) For instance, the trifluoroacetic acid (TFA) typically used during the cleavage reaction can cause proton-catalyzed deglycosylation. We, therefore, developed alternative reaction conditions that are compatible with DNA. BF_3_ etherate (BF_3_·Et_2_O)-induced Edman degradation in acetonitrile (40 mM BF_3_·Et_2_O, 5 min) has been reported to reduce proton-catalyzed racemization while completely cleaving PITC-modified N-terminal amino acids.(*31, 32*) Moreover, BF_3_ methanol complex has been used as a non-depurinating detritylation reagent in DNA synthesis.(*33*) Encouraged by these reports, we hypothesized that DNA stability would be enhanced in a cleavage reaction mediated by BF_3_·Et_2_O in acetonitrile.

**Fig. 2.**
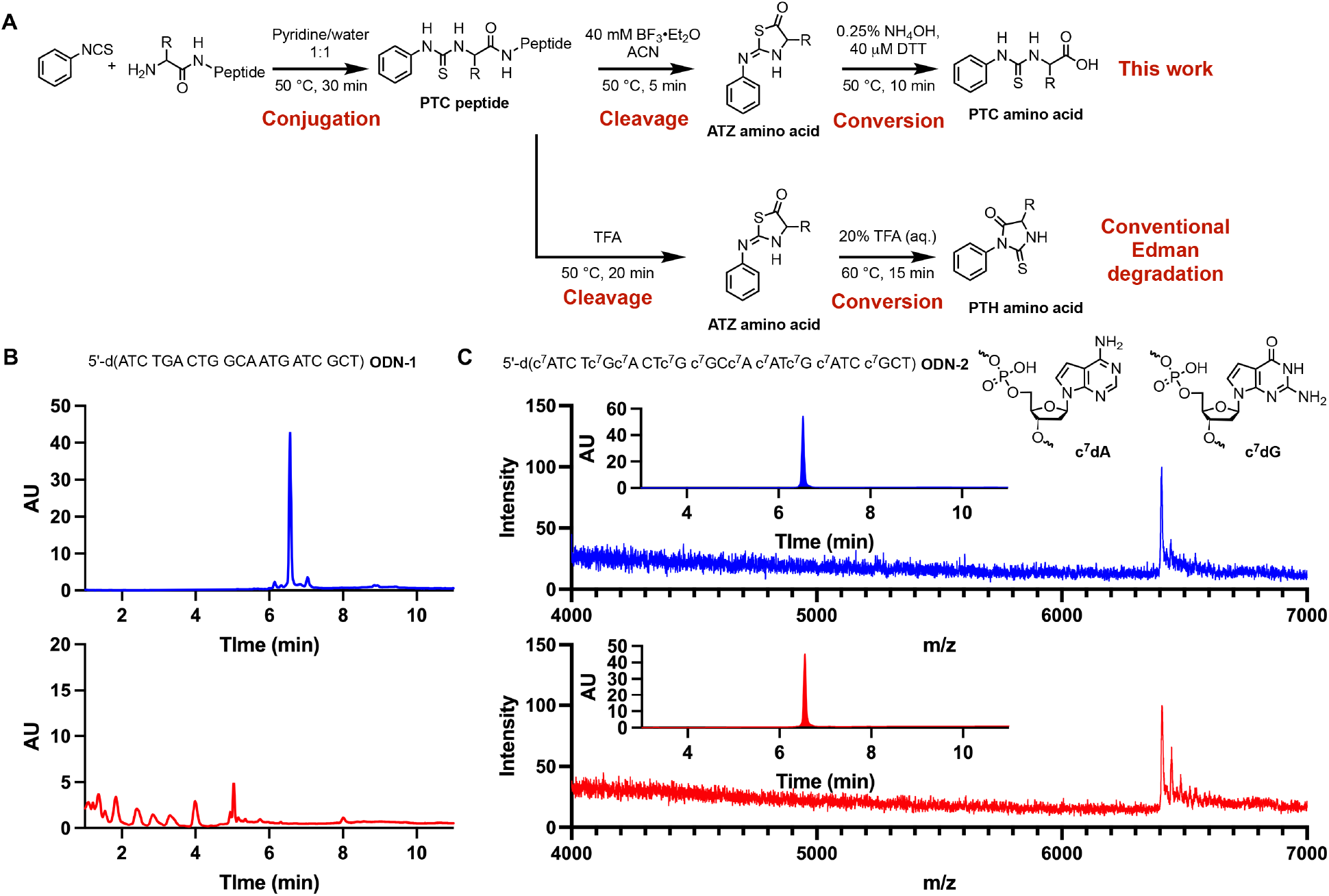
Developing a DNA-compatible Edman degradation reaction. (**A**) Reaction scheme of conventional Edman degradation (bottom) and the BF_3_·Et_2_O-mediated version presented in this work (top). The TFA used in conventional Edman degradation is incompatible with DNA. We instead use BF_3_·Et_2_O, which generates ATZ amino acids that are then converted to more synthetically accessible PTC amino acids under reducing, basic conditions. (**B**) HPLC chromatograms of oligonucleotide **ODN-1** before (top) and after (bottom) treatment with 40 mM BF_3_·Et_2_O in anhydrous acetonitrile for 30 min. (**C**) MS spectra of oligonucleotide **ODN-2** before (top) and after (bottom) treatment with 40 mM BF_3_·Et_2_O in anhydrous acetonitrile for 4 h. Inset: HPLC chromatograms of **ODN-2** before and after the treatment.

However, no intact DNA was observed after 30 min when we treated an oligodeoxynucleotide (ODN) containing all four deoxynucleotides (**ODN-1**) with 40 mM BF_3_·Et_2_O in anhydrous acetonitrile (**Fig. 2B**). To better understand the lability of DNA during BF_3_·Et_2_O treatment, we analyzed the degradation products formed by a poly-dT sequence bearing a single dA (5′-d(T_14_AT_4_), **ODN-S1**) by MS (**Fig. S1, A** and **B**). The major product (m/z = 4,577.3) is consistent with depurination of the single dA and subsequent strand cleavage;(*34*) no cleavage events at dT sites were detected. A similar cleavage pattern was also observed for an oligonucleotide containing a single dG (5′-d(T_14_GT_4_), **ODN-S2**; **Fig. S1, C** and **D**). Based on these results, we concluded that the degradation of native DNA during BF_3_·Et_2_O treatment was largely due to the instability of purine deoxynucleotides. We therefore explored the use of chemically modified 7-deazapurine deoxynucleotides (c7dA and c7dG), which have been reported to be resistant to depurination.(*35, 36*) We synthesized **ODN-2**, a modified version of **ODN-1**, in which dA and dG were substituted with c7dA and c7dG, respectively. This modified oligonucleotide was unaffected by BF_3_·Et_2_O treatment for 4 h (**Fig. 2C**). Given that N-terminal amino acid cleavage in the presence of 40 mM BF_3_·Et_2_O occurs within 5 min, the stability of 7-deazapurine-modified DNA should be more than sufficient for the Edman degradation process. Importantly, 7-deazapurine deoxynucleotide triphosphates are accepted by several commonly used DNA polymerases (e.g., Klenow fragment (exo-), Sequenase version 2.0, and *Bst* 3.0), and primer extension reactions using c7dATP and c7dGTP proceeded with efficiency comparable to that of dATP and dGTP (**Fig. S2**).

As with the standard TFA-mediated cleavage reaction during Edman degradation, anilinothiazolinone (ATZ) amino acids are the major product of the BF_3_·Et_2_O-mediated degradation reaction.(*31*) However, ATZ amino acids are unstable and thus not suitable for antibody-based detection. To overcome this problem, we converted the ATZ amino acids into stable PTC amino acids under alkaline reducing conditions (**Fig. 2A**).(*37, 38*) Dithiothreitol (DTT) was added to inhibit oxidative degradation of PTC amino acids.(*39*) Importantly, PTC amino acids can readily be prepared by reacting PITC or its derivatives with amino acids, greatly simplifying the generation of antibodies against these modified amino acids.

### DNA-encoded Edman degradation

We next developed a chemical process for iteratively barcoding the PTC amino acids generated by our modified Edman degradation reaction (**Fig. 3A**). We performed the degradation reaction on model peptides bearing an azidolysine immobilized onto magnetic beads, which simplifies solvent exchange during the multi-step chemical process while also increasing reagent accessibility to DNA-peptide conjugates when reactions are conducted in neat organic solvents.(*40–42*) To immobilize the azide-modified peptides, we prepared carboxylic acid beads conjugated with DBCO-modified barcode DNA. Because the conjugation of DNA onto the carboxylic acid beads and the conjugation of the DBCO functional group onto the peptide barcode DNA both utilize amino modification of DNA, we performed these reactions in a stepwise fashion. We first immobilized the 5′-amino-modified anchor DNA (**ODN-3**) onto the carboxylic acid beads by carbodiimide coupling.(*43*) We then enzymatically ligated DBCO-modified 5′-phosphorylated DNA containing the primer binding site (**ODN-4**) onto that anchor DNA (**Fig. 3A**, step 1) to form the complete peptide barcode DNA. Subsequently, we conjugated a seven amino acid (aa) peptide (FGGGGGX, X = azidolysine) to the DNA via a strain-promoted alkyne-azide cycloaddition (SPAAC) reaction (**Fig. 3A**, step 2), and reacted azide-modified PITC (**1**) with the N-terminus of the peptide (**Fig. 3A**, step 3). To evaluate the yield of each step, we synthesized a Cy3-labeled anchor DNA bearing an internal disulfide linkage (**ODN-S10, Table S1)**, which enables the release of intermediates from the magnetic beads under reducing conditions. These released intermediates were then analyzed via gel electrophoresis. Step 1 to 3 all proceeded with near quantitative yields (**Fig. 3B**).

**Fig. 3.**
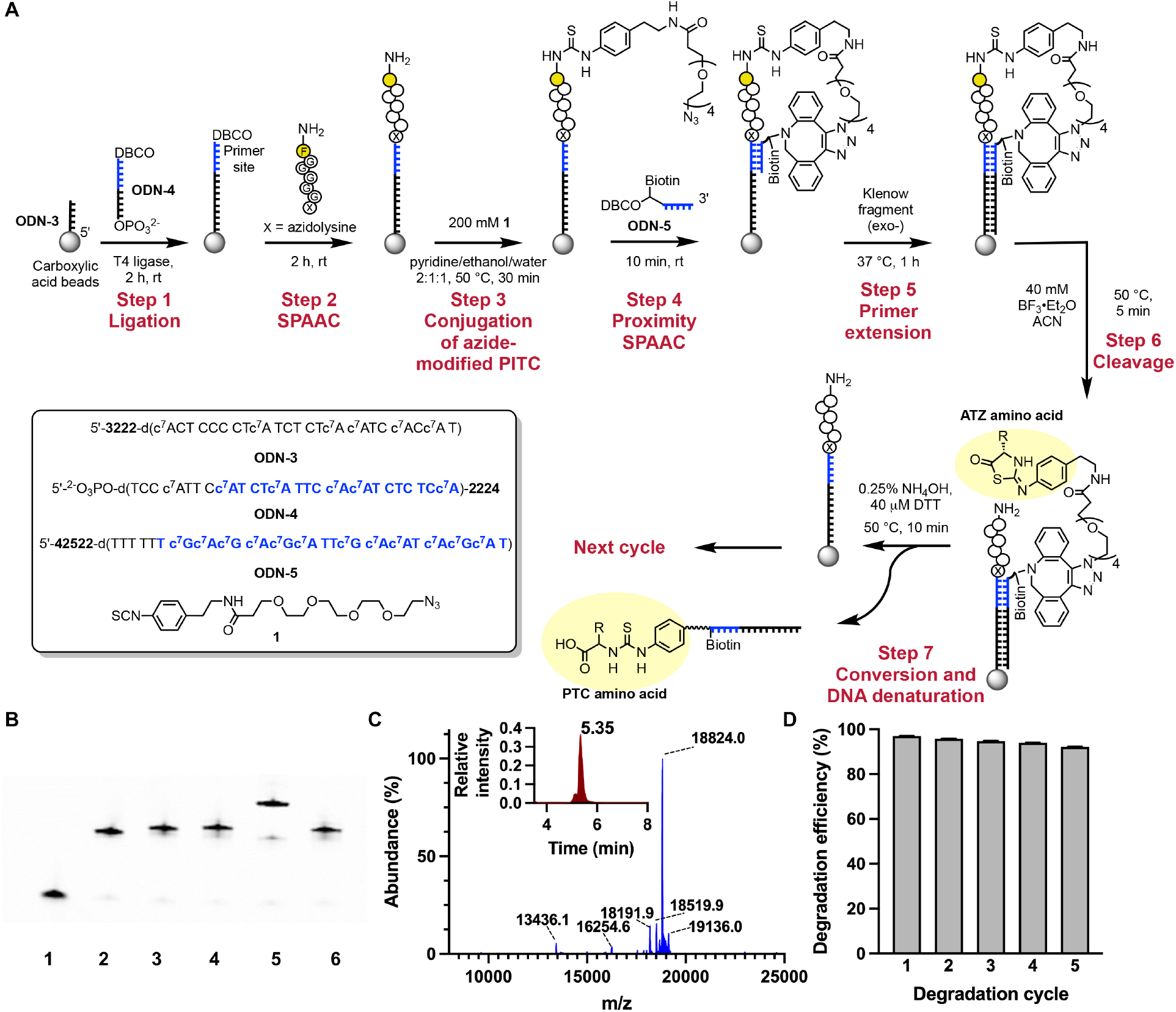
DNA-encoded Edman degradation on magnetic beads. (**A**) Reaction scheme for barcoding PTC amino acids via DNA-encoded Edman degradation on magnetic beads. Anchor DNA (**ODN-3**) is immobilized on carboxylic acid beads via carbodiimide coupling and then ligated to DBCO-modified 5′-phosphorylated DNA (**ODN-4**) (step 1). The azidolysine-modified peptide FGGGGGX is attached to **ODN-4** via click reaction (step 2), after which the peptide’s N-terminus is reacted with azide-modified PITC (**1**) (step 3). The DBCO-modified primer (**ODN-5**) is then coupled via proximity SPAAC reaction (step 4). Next, primer extension generates a complementary sequence to the peptide barcode DNA (step 5). Cleavage by BF_3_·Et_2_O-mediated Edman degradation yields a DNA-barcoded ATZ amino acid, which remains hybridized on the bead due to the insolubility of DNA in acetonitrile (step 6). Finally, hydrolysis under reducing basic conditions converts the ATZ amino acid to a PTC amino acid and denatures the DNA duplex, releasing the DNA-barcoded PTC amino acid from the solid support (step 7). The DNA used in this experiment contains non-nucleotide modifications: hexaethylene glycol (**2**), hexylamine (**3**), DBCO (**4**), biotin-dT (**5**). The structures of the modifications are shown in **Table S2**. (**B**) Denaturing PAGE analysis of DNA-encoded Edman degradation. We used a disulfide-modified anchor DNA (**ODN-S10**) that enables the release of intermediates from magnetic beads by reduction. We analyzed aliquots taken after anchor DNA immobilization (lane 1), DNA ligation (lane 2), conjugation of peptide (lane 3), conjugation of azide-modified PITC (lane 4), and conjugation of DBCO-modified primer (**ODN-5**, lane 5). In lane 6, we used a DBCO-modified poly-dT sequence (**ODN-S12**) to carry out the click reaction as a negative control, and no click product was observed after 10 min. (**C**) MS spectrum of DNA-barcoded PTC-F generated by the first cycle of DNA-encoded Edman degradation on peptide FGGGGGX (calculated molecular weight 18,824.2, observed molecular weight 18,824.0). Peaks at 13,436.1, 16,254.6, 18,191.9, and 18,519.9 correspond to truncated products resulting from incomplete primer extension. The peak at 19,136.0 corresponds to the product from A-tailing. The inset shows HPLC chromatograms of DNA-barcoded PTC-F. (**D**) Efficiency of DNA-encoded Edman degradation over five cycles as measured by flow cytometry. Stepwise yields are 96.9 ± 0.1%, 95.8 ± 0.1%, 94.75 ± 0.08%, 93.94 ± 0.05%, 92.1 ± 0.1%.

For the barcode transfer step, we first conjugated a DBCO-modified primer (**ODN-5**) to the azide group on the PTC peptide via proximity SPAAC (**Fig. 3A**, step 4). Although SPAAC is generally limited by its relatively slow reaction rate (k = 0.2–0.5 M-1s-1),(*44*) we opted for this approach due to the susceptibility of the PTC group to oxidation by reactive oxygen species that are generated during copper (I)-catalyzed azide-alkyne cycloaddition (CuAAC).(*45, 46*) Fortunately, hybridization between the template and primer accelerated the SPAAC reaction by increasing the effective local concentration of reactants,(*47*) and we found that the primer conjugation reaction reached completion with peptide FGGGGGX within 10 min (**Fig. 3B**). In contrast, when the reaction was carried out using a non-complementary poly-dT sequence (**ODN-S11, Table S1**), no detectable conjugation was observed after 10 min, highlighting the importance of DNA hybridization in accelerating the reaction. This rate acceleration of the proximity SPAAC reaction was general. The click reaction reached completion within 1 h for various peptides ranging from 10–30 aa in length and of varying rigidities (**Fig. S3**).

The primer was subsequently extended using Klenow fragment (exo-) (**Fig. 3A**, step 5), after which the beads were subjected to Edman degradation with BF_3_·Et_2_O in acetonitrile (**Fig. 3A**, step 6). During this step, we took advantage of the insolubility of DNA in acetonitrile,(*48*) which allows the DNA-barcoded ATZ amino acids to remain hybridized with the template DNA following the cleavage reaction. The subsequent basic conversion reaction converts ATZ amino acids into stable PTC amino acids while also denaturing the duplex DNA, releasing the barcoded amino acids (**Fig. 3A**, step 7).(*37*) The DBCO-modified primer (**ODN-5**) contains a dT_5_ region that enables the introduction of a Cy3-labeled complementary strand (**ODN-S12, Table S1**) via toehold-mediated strand displacement. This allowed us to quantify the yield of the cleavage and conversion reactions by flow cytometry. We confirmed the structure of the cleaved barcoded PTC amino acid via LC-MS (**Fig. 3C)** and measured a cleavage yield of 96.9 ± 0.1% (**Fig. 3D**). We also validated that peptides bearing N-terminal glycine, tryptophan, tyrosine, aspartic acid, arginine or phosphotyrosine were compatible with this process. Correctly barcoded PTC amino acids were observed in all cases, with degradation yields comparable to that of N-terminal phenylalanine (**Fig. S4** and **Fig. S5**).

Finally, we confirmed that we could perform this process over multiple rounds by carrying out five iterative cycles of cleavage with FGGGGGX. The degradation products from each round were isolated, and their structures were confirmed by LC-MS (**Fig. S6**). The degradation yield for each cycle was determined by flow cytometry as described above (**Fig. 3D**). We observed a slight decrease in available peptides after each cycle, as indicated by a reduction of fluorescence signal before degradation (**Fig. S6**). It is unlikely that this is due to the instability of the DBCO-azide linkage since no decomposition of this linkage was observed by MS (**Fig. 3C**). We hypothesize that the decrease in available peptides is caused by processes that irreversibly block the N-terminus, such as oxidative desulfurization of the PTC moiety.(*45, 49*)

### Conversion of barcoded PTC amino acids to DNA reporters

The DNA-barcoded PTC amino acids are subsequently converted into DNA reporters by an antibody binding-mediated PEA. For PTC amino acids with no existing antibody, we generated novel monoclonal antibodies by immunizing mice with bovine serum albumin (BSA) conjugated to various PTC amino acids (**Fig. S7**). This process yielded antibodies that achieved good specificity and affinity for PTC-modified phenylalanine (PTC-F, **Fig. 4A**), tryptophan (PTC-W, **Fig. 4B**), aspartic acid (PTC-D, **Fig. 4C**), and arginine (PTC-R, **Fig. 4D**), as measured by biolayer interferometry (BLI, **Fig. S8**). An anti-PTC tyrosine (PTC-Y, **Fig. 4E**) antibody was also obtained with only modest specificity against PTC-F, but high specificity against other non-target PTC amino acids. For some PTC amino acids, their side chains remain intact and unobstructed, such that existing antibodies targeting these side chains—including antibodies against PTMs—may maintain their affinity. We experimentally confirmed that commercially available antibodies against phosphotyrosine (pY), asymmetric dimethylarginine (ADMA), phosphoserine (pS), and acetylated lysine (AcK) all retained their high affinity for corresponding PTC amino acids (**Fig. 4F** and **Fig. S9**).

**Fig. 4.**
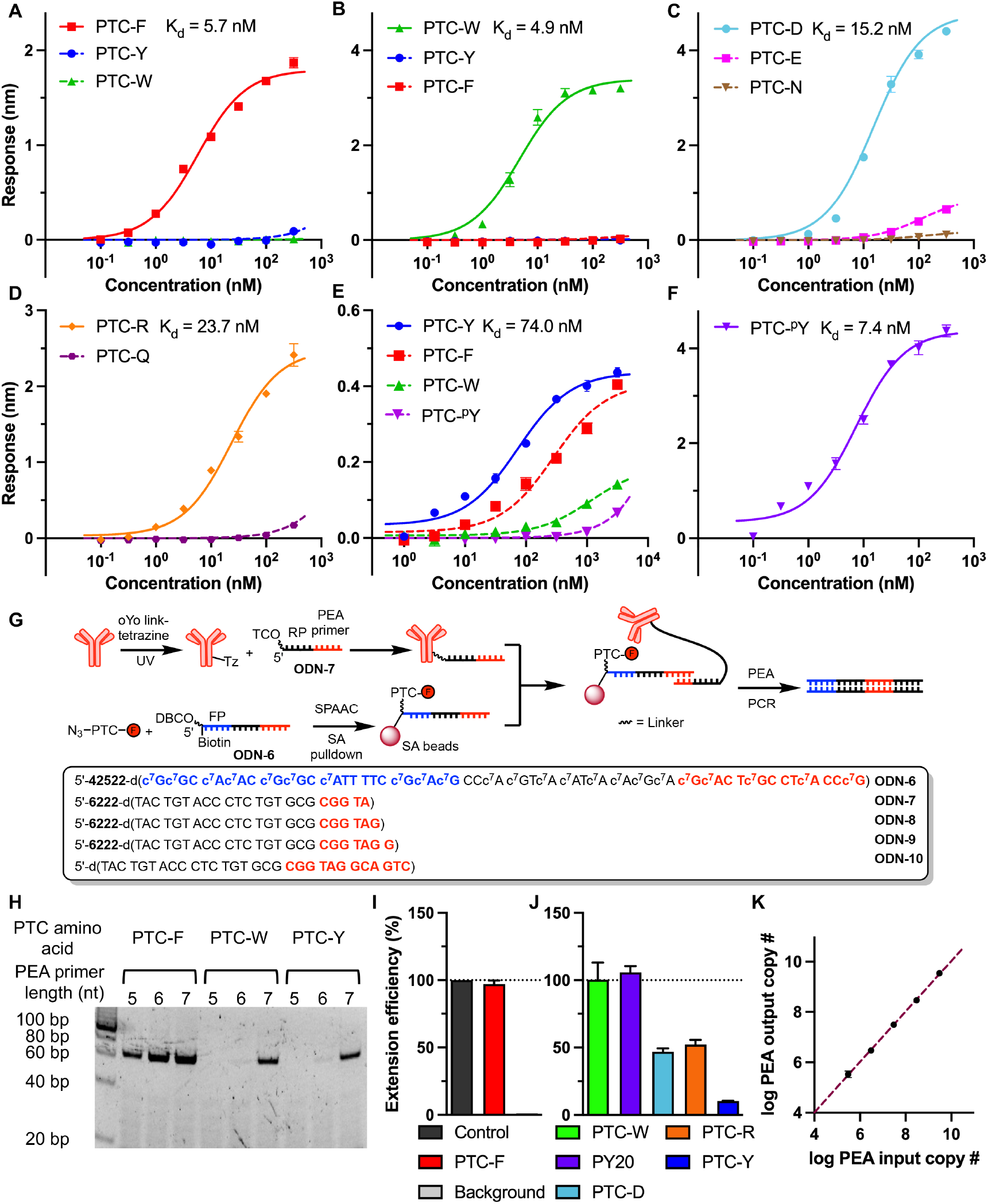
PEA-mediated conversion of DNA-barcoded PTC amino acids into DNA reporters. (**A** to **F**) Binding curves of (A) anti-PTC-F, (B) anti-PTC-W, (C) anti-PTC-Y, (D) anti-PTC-D, (E) anti-PTC-R, and (F) PY20 antibody against target and non-target DNA-tagged PTC-amino acids (PTC-E = PTC glutamic acid, PTC-N = PTC asparagine, PTC-Q = PTC glutamine), as measured by biolayer interferometry (BLI). Calculated K_d_ values for the primary antibody target are shown in each plot. (**G**) Scheme for converting DNA-barcoded PTC amino acids to DNA reporters. First, anti-PTC-F antibody is modified with photoreactive antibody-binding domains (i.e., oYo link) to introduce a tetrazine (Tz) functional group onto the Fc region. Next, a TCO-modified DNA strand featuring a PEA primer sequence of 5–7 nt (**ODN-7, −8**, or **-9**) is conjugated with the Tz group to form a primer-antibody conjugate. PTC-F is tagged with biotinylated template DNA (**ODN-6**) and immobilized onto SA beads. The beads are incubated with primer-modified anti-PTC-F antibody, and then Klenow fragment (exo-) is added to perform PEA. The resulting product is PCR amplified for further analysis. **ODN-7, −8**, or **-9** contain TCO modification (**6**). The structure is shown in **Table S2**. (**H**) Native PAGE of products generated by PEA with primers of varying lengths. 5 nt and 6 nt PEA primers (**ODN-7** and **ODN-8**) can specifically distinguish antibody-antigen interactions, whereas the longer 7 nt primer (**ODN-9**) produces a non-specific signal. (**I**) Quantification of DNA output of PEA by qPCR. PEA was carried out with 30 nM **ODN-8**-tagged anti-PTC-F antibody and **ODN-6-PTC-F** immobilized on SA beads. For the positive control, the primer extension was performed using a 12 nt primer (**ODN-10**, 30 nM) and **ODN-6** immobilized on SA beads without PTC-F conjugation. A non-specific PEA was performed as a negative control with similarly **ODN-6**-modified SA beads and 30 nM **ODN-8**-tagged anti-PTC-F antibody. (**J**) PEA efficiency of other anti-PTC amino acid antibodies. All antibodies were modified with **ODN-8**, and PEA was carried out with 100 nM primer-antibody conjugate. (**K**) Detection of a dilution series of **ODN-6-PTC-F** immobilized on SA beads by PEA in 20 µL reactions. PEA input copy number is calculated based on the amount of **ODN-6-PTC-F** added to the SA pulldown. PEA output copy number is calculated based on the qPCR standard curve. The data is fitted using linear regression with equation *y* = 1.003*x* - 0.0008015 and R^2^ = 0.999.

We functionalized the Fc domain of anti-PTC amino acid antibodies with tetrazine (Tz) using photoreactive antibody-binding domains (**Fig. S10, A** and **B**). The functionalized antibodies were coupled via click chemistry to trans-cyclooctene (TCO)-modified DNA sequences consisting of the PCR reverse primer (RP) and the PEA primer (e.g., **ODN-7, Fig. 4G** and **Fig. S10C**). Independently synthesized PTC amino acids were conjugated with a model barcode DNA (**ODN-6**) containing the PCR forward primer (FP) and the PEA primer binding site via SPAAC and pulled down on SA beads (**Fig. 4G**). The immobilized DNA-barcoded PTC amino acids were recognized by their corresponding primer-functionalized antibodies, and this binding event stabilized the primer-template complex and enabled primer extension.

In traditional PEA, a quantitative DNA output is not required for the quantification of the molecule of interest.(*50*) However, PEA efficiency is critical for our method to achieve high sequencing coverage and single-molecule sensitivity. Specificity is also essential for achieving high-fidelity reads. There are two competing reactions during proximity extension: specific primer extension that only occurs with help from antibody-antigen binding, and non-specific primer extension initiated in the absence of antibody-antigen recognition. The efficiency of specific primer extension is affected by both the affinity of the antibodies and the stability of the primer-template complex, while the extent of non-specific primer extension is only affected by the latter. Thus, tuning primer-template complex stability could increase the yield of specific primer extension and suppress the non-specific one. To test this, we assessed the specificity and efficiency of proximity extension when barcoded PTC-F, -W, or -Y were incubated with anti-PTC-F antibodies modified with PEA primer sequences ranging in length from 5–7 nt (**ODN-7**, -**8**, or -**9**, respectively). Considerable non-specific primer extension was observed with **ODN-9**, whereas the shorter primers favored specific primer extension (**Fig. 4H**). Next, we evaluated the efficiency of proximity extension by qPCR. At a concentration of 100 nM, anti-PTC-F antibody modified with **ODN-7** only converted 24% of PTC-F molecules to a DNA output (**Fig. S11D**). On the other hand, **ODN-8**-modified anti-PTC-F yielded complete primer extension at the same concentration (**Fig. S11F**). In addition to primer length, the stability of primer-template complexes is also affected by the primer concentration. Because non-specific primer extension solely originates from primer-template hybridization, it can be greatly inhibited by reducing the primer concentration. With 30 nM **ODN-8**-conjugated anti-PTC-F, the extent of non-specific primer extension was just 1% of that of specific primer extension, and the reduced concentration had minimal impact on the yield of specific primer extension (**Fig. 4I**). In contrast, increasing the antibody concentration to 300 nM only increased the extent of non-specific primer extension (**Fig. S11F**). PEA efficiencies were determined for other antibodies conjugated with **ODN-8**. Specific primer extension reactions were observed for all antibodies, with a positive correlation between the yield of the proximity extension reaction and the affinity of the antibody (**Fig. 4J**). Lastly, for a certain pair of primer-modified antibody and PTC amino acid, the efficiency of proximity extension is determined by the antibody concentration (see Supplementary Text). Therefore, reducing the amount of input PTC amino acids should not affect PEA performance. We tested this hypothesis with PTC-F/anti-PTC-F and observed that the efficiency of proximity extension remained quantitative for input quantities as low as 10 amol **ODN-6-PTC-F** in a 20 µL reaction (**Fig. 4K, Fig. S11, H** and **I**).

### Sequencing of model peptides

Having optimized the various steps of our protocol, we carried out proof-of-concept sequencing on a model peptide (**Fig. 5A**). We prepared carboxylic acid beads as described above, ligating the bead-immobilized anchor DNA (**ODN-11**) with a peptide-barcoding strand featuring a 2 nt peptide barcode (**ODN-12**). We then immobilized the 7 aa peptide RGFDWGX, which consists of four amino acids that can be detected by previously mentioned anti-PTC amino acid antibodies (anti-PTC-R, -F, -D, and -W). The glycine residue serves as an exemplar of an amino acid that cannot be detected currently, which allows us to assess the true background in the absence of a specific antibody recognition event during the sequencing process. We performed five cycles of sequencing on ∼100 pmol peptide. To introduce cycle number-specific barcodes during the adaptor PCR step, the PTC amino acids from each cycle were collected and processed separately. The DNA-barcoded PTC amino acids were enriched by SA bead pulldown, yielding a final loading density of ∼10 pmol biotinylated DNA-barcoded PTC amino acids per mg of beads. Then PEA was carried out with a mixture of DNA-modified anti-PTC-R, -F, -D, and -W antibodies, where each DNA strand comprised the RP sequence for PCR amplification, a 3 nt antibody barcode, and a 6 nt PEA primer sequence (**ODN-13, ODN-S23** to **S25, Table S1**). After PEA, sequencing adaptors were incorporated via PCR, during which 2 nt cycle number barcodes were introduced with forward adaptor primers (**ODN-14, ODN-S26** to **S29, Table S1**). The PCR products were then combined for indexing and sequencing. After sequencing, barcodes encoding cycle number and antibody identity were extracted and counted, and the raw counts of reads bearing the same antibody barcode were normalized by setting the highest read count to 100 (**Fig. 5B**). Overall, the correct amino acid assignment for each position consistently produced the highest normalized read count, producing signals that were 20–100 times above the background. While sequencing the undetectable glycine in the second cycle, we observed a normalized read count comparable to that of background in other cycles. This demonstrates that undetectable amino acids at a given position can be clearly identified rather than being incorrectly assigned. The raw read counts were generally consistent with degradation yield and proximity extension efficiencies established previously (**Fig. S12A**).

**Fig. 5.**
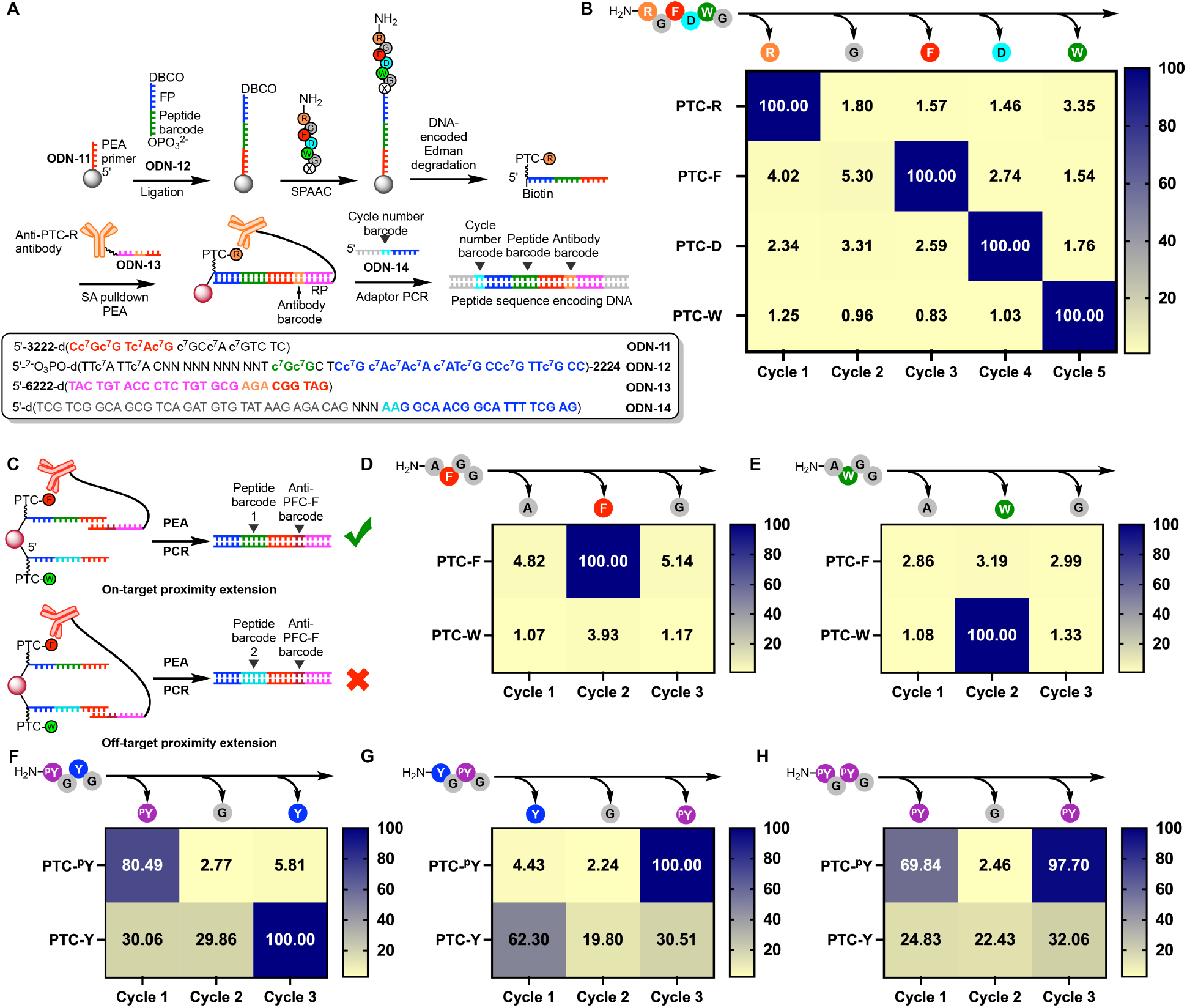
Demonstrations of peptide sequencing. (**A**) Overview of the complete peptide “reverse translation” process. A DNA strand bearing the FP and 2 nt peptide barcode (**ODN-12**) is ligated onto anchor DNA (**ODN-11**)-modified carboxylic acid beads, forming the complete peptide barcode DNA. The model peptide RGFDWGX is then linked to the DNA strand via SPAAC and subjected to our modified Edman degradation process, which yields a DNA-barcoded PTC amino acid. This is recognized by an antibody barcoded with a DNA consisting of the RP sequence, a 3 nt antibody barcode, and a 6 nt PEA primer (e.g., **ODN-13** for PTC-R). The resulting PEA product is amplified by adaptor PCR, during which a 2 nt cycle number barcode is introduced through barcoded forward adaptor primers (e.g., **ODN-14** for cycle 1). The final DNA reporter sequence contains barcodes for peptide, antibody (i.e., amino acid identity), and cycle number (i.e., amino acid position). (**B**) Heatmap of normalized barcode read counts from the first five residues of RGFDWGX. The raw counts of reads bearing the same antibody barcode were normalized by setting the highest read count to 100. The correct residue at each position is shown at the top. (**C**) Scheme of on-target and off-target proximity extension, the latter of which leads to barcode crosstalk. (**D** and **E**) Heatmaps of normalized read counts for peptide sequences (D) AFG and (E) AWG after sequencing an equimolar mixture of the two peptides. (**F** to **H**) Heatmaps of normalized read counts for peptide sequences (D) ^p^YGY, (G) YG^p^Y, and (H) ^p^YG^p^Y after analysis with PY20 and anti-PTC-Y antibodies. Normalization for (D to E) was applied as in (B).

Our procedure is also capable of sensitively discriminating subtle sequence differences in mixtures of peptides. One major challenge in processing multiple peptide species is the possibility of barcode crosstalk during PEA (**Fig. 5C**), where antibody binding to one PTC amino acid may initiate primer extension on another nearby template. We could address this by reducing the loading density of DNA-barcoded PTC amino acid on the beads—at lower loading densities, the effective template concentration for on-target proximity extension is unaffected. In contrast, the effective template concentration for off-target proximity extension is reduced, yielding a substantially smaller off-target DNA reporter readout. We performed preliminary screening of loading density and determined that the extent of off-target proximity extension was reduced to 3% of that of on-target proximity extension at 0.5 pmol/mg loading (**Fig. S13, A** and **B**).

Having demonstrated the feasibility of minimizing barcode crosstalk between sequencing readouts for different peptides, we set out to distinguish two peptides that differ by a single amino acid (AFGGGX and AWGGGX) by sequencing the first three amino acids in parallel. To sequence these two peptides simultaneously, we slightly modified the initial stages of our method; each of the two peptides was first conjugated with a unique peptide barcode, after which these DNA-barcoded peptides were mixed in an equimolar ratio and immobilized onto anchor strand-modified magnetic beads by ligation. Subsequent degradation and sample processing steps were performed similarly to the single-peptide sequencing process described above. The DNA-barcoded PTC amino acids were detected with an equimolar mixture of primer-conjugated anti-PTC-F and -W antibodies (30 nM each). The resulting primer extension products were then barcoded with cycle number barcodes and sequenced. After separating our analyses based on the peptide-identifying barcodes, we confirmed that the normalized read counts accurately reflected each peptide sequence, such that the two could be clearly distinguished (**Fig. 5, D** and **E**).

Finally, we sequenced three peptides with different phosphorylation patterns in parallel: ^p^YGYGGX, YGpYGGX, and pYGpYGGX. Existing methods like enzyme-linked immunosorbent assay (ELISA) and MS often struggle to identify the correct position of a PTM given multiple candidate sites. Because our method allows sequential analysis of individual amino acids, phosphorylation sites could be accurately assigned when sequenced with anti-pY (PY20, 30 nM, **Fig. 5, F** to **H**). We also sequenced the mixture with anti-PTC-Y antibody (100 nM) separately, which is less efficient during PEA due to its low affinity (**Fig. 4J**) but demonstrates specificity for PTC-Y over PTC-pY (**Fig. 4E**). Although the readouts here were also consistently accurate, the signal-to-noise ratio was much lower than that of PY20, in concordance with the proximity extension efficiency of anti-PTC-Y antibody. Collectively, these results highlight our platform’s capacity to accurately discriminate both native and post-translationally modified versions of the same amino acid within a peptide sequence.

## Conclusions

Our “reverse translation” strategy for peptide sequencing offers numerous advantages relative to other methods described to date. The Edman degradation-based method employed here excises one amino acid at a time, enabling each to be analyzed independently outside the larger context of the peptide. This enables peptide sequencing at single-amino acid resolution and eliminates the potential for interference from adjacent amino acids or PTMs. The released amino acids are subsequently detected by antibodies, which can, in principle, be directly generated against the full spectrum of proteinogenic amino acids. In this work, we have demonstrated such capabilities with four antibodies that exhibit good affinity and specificity for their natural amino acid targets. Furthermore, we demonstrated that we can leverage PTM-specific antibodies to accurately identify the presence and position of such modifications. Our method converts peptide sequence information into DNA strands. These DNA strands can be amplified by PCR, enabling detection with single-molecule sensitivity.(*51, 52*) Thus, the sensitivity of our method ultimately depends on the efficiency of PEA. Using the best-performing antibodies, we have demonstrated that PEA can quantitatively convert 10 amol of PTC amino acids to DNA reporters in a 20 µL reaction containing approximately 1,000 beads (**Fig. 4K**). Since the efficiency of PEA is determined by the antibody concentration, in theory, the sensitivity of our assay can be further enhanced by dividing the bulk assay into droplet-based single-bead assays, which has been shown to enable the detection of targets with single-molecule sensitivity.(*53–55*) Additionally, by incorporating unique molecular identifiers, our method should be applicable to the sequencing of complex peptide mixtures as well as the digital quantification of peptides.(*56*) Finally, this approach employs conventional reagents (e.g., antibodies and oligonucleotides) and does not require specialized instrumentation. The DNA that encodes peptide sequences is analyzed using high-throughput sequencing platforms, the current generation of which can generate tens of billions of reads per run, making this strategy cost-effective and highly scalable.

However, several challenges remain in realizing the sequencing of native peptides with single-molecule sensitivity. First, our method requires the conjugation of DNA to the C-terminus of peptides, which we achieved in this work by modifying the C-termini of our peptides with azidolysine residues. While several strategies for C-terminus-specific labeling have been reported,(*57–60*) they are limited in their substrate scope, and a generalizable C-terminus-specific modification reaction is currently unavailable. Alternatively, cysteine can serve as a handle for conjugation, but this approach restricts the scope of peptides that can be sequenced.(*61*) Additionally, further improvements in the efficiency of DNA-encoded Edman degradation and degradation fragment recovery are needed to accommodate longer sequencing reads. Although we have not observed loss of synchrony due to incomplete cleavage during the sequencing of short model peptides, this potential issue could be addressed by implementing cycle number barcodes during Edman degradation. Finally, the scope, sensitivity, and accuracy of our sequencing method are determined by the performance of our PTC amino acid-specific antibodies. Some applications may not require the identification of all amino acids, and indeed, there are already commercially available instruments that focus on peptide identification based on limited sequence profiling. However, the *de novo* sequencing of proteins with single-molecule sensitivity will require a complete set of amino acid-specific antibodies. We have demonstrated a complete pipeline for antibody generation via immunization. However, generating antibodies with high affinity and sufficient specificity to discriminate structurally similar amino acids via this process may be challenging. Further improvement of antibody affinity can be achieved via directed evolution,(*62*) and specificity could also be reprogrammed from existing cross-reactive binders.(*63*) Alternatively, new binders could be generated using emerging computational design methods.(*64, 65*)

In summary, our work lays the groundwork for sequencing peptides by converting their sequence to DNA. With the expansion of the antibody arsenal and further refinement and streamlining of our chemical process, we believe that our method will be capable of high-throughput *de novo* protein sequencing of both unmodified and PTM-bearing proteins with single-amino acid resolution and single-molecule sensitivity, opening up exciting new opportunities for single-molecule proteomic analysis.

## Acknowledgments

We thank Dr. Adrian Hugenmatter at Stanford Innovative Medicines Accelerator for the helpful discussion. We thank Joshua Lowitz at Antibody Solutions for his assistance with custom antibody generation. We thank Theresa McLaughlin at Stanford University Mass Spectrometry for her assistance with developing LC-MS methods for oligonucleotides. This work was supported by the Vincent Coates Foundation Mass Spectrometry Laboratory, Stanford University Mass Spectrometry (RRID:SCR_017801) utilizing the Bruker Microflex MALDI TOF mass spectrometer (RRID:SCR_018696) and Thermo Exploris 240 LC/MS system (RRID:SCR_022216) that was purchased with funding from Stanford C-ShaRP (RRID:SCR_022986).

## Author contributions

Conceptualization: L.Z., H.T.S.

Methodology: L.Z.

Investigation: L.Z., Y.S.

Visualization: L.Z., Y.S.

Funding acquisition: H.T.S.

Project administration: H.T.S.

Writing – original draft: L.Z., H.T.S.

Writing – review & editing: all authors

## Data and materials availability

All data are available in the main text or the supplementary materials. Next-generation sequencing data will be made available via figshare upon publication.

## Supplementary Materials

Materials and Methods

Supplementary Text

Figs. S1 to S13

Tables S1 to S4

References (*66*)

MS Spectra of modified oligonucleotides

NMR spectra of organic compounds

