## Supplementary Materials V2 for "Peptide sequencing via reverse translation of peptides into DNA"

**The PDF file includes:**

Materials and Methods
Supplementary Text
Figs. S1 to S13
Tables S1 to S4
Reference (66)
MS Spectra of modified oligonucleotides
NMR spectra of organic compounds

### Materials and Methods

#### Materials

Unmodified oligonucleotides were purchased from IDT unless otherwise stated (**Table S1**). Peptides were purchased from Genscript (**Table S3**). Phosphoramidites, oligonucleotide synthesis reagents, DBCO-sulfo-NHS ester, and azidobutyrate NHS ester were purchased from Glen Research. Native deoxynucleotide triphosphates, T4 DNA ligase, Klenow fragment (3'→5' exo-), and *Bst* 3.0 DNA polymerase were purchased from New England Biolabs. 7-Deaza-2'-deoxyadenosine-5'-triphosphate ($c^7$dATP) and 7-deaza-2'-deoxyguanosine-5'-triphosphate ($c^7$dGTP) were purchased from Jena Bioscience. N-Hydroxysuccinimide (NHS), 1-ethyl-3-(3-dimethylaminopropyl)carbodiimide (EDC), L-amino acids, 4-(2-aminoethyl)aniline, Tween-20, anhydrous acetonitrile (ACN), anhydrous dimethyl sulfoxide (DMSO), anhydrous dichloromethane (DCM), anhydrous triethylamine (TEA), anhydrous N,N-diisopropylethylamine (DIPEA), tris(2-carboxyethyl)phosphine (TCEP), DL-dithiothreitol (DTT), and $BF_3$ etherate ($BF_3 \cdot Et_2O$) were purchased from Sigma Aldrich. Azido-PEG4-NHS ester and TCO-NHS ester (axial) were purchased from Click Chemistry Tools. Tetramethylammonium trifluoromethanethiolate ($(Me_4N)SCF_3$) was purchased from Oakwood Chemical. Phosphotyrosine antibody (PY20) was purchased from Abcam. The oYo-Link tetrazine and tetrazine mIgG1 kits were purchased from AlphaThera. All reagents for DNA sequencing were obtained from Illumina. Dynabeads MyOne carboxylic acid, Dynabeads MyOne streptavidin C1, ultrapure bovine serum albumin (BSA), Sequenase version 2.0 DNA polymerase, phosphoserine antibody (3C171), asymmetric dimethyl arginine antibody (21C7), and acetylated lysine antibody (RM101), HPLC grade organic solvents (ACN, DCM, methanol (MeOH), pyridine, ethanol, and hexanes), 1 M Tris hydrochloride (pH 7.5), 1× PBS (pH 7.4), 1 N NaOH, 1 M $MgCl_2$, 3 M NaOAc, 5 M NaCl, ammonium hydroxide (28-30% solution in water), methyl-PEG12-amine, TFP Ester-PEG4-DBCO, deuterated solvents for NMR, and other reagents were purchased form Thermo Fisher Scientific.

#### Methods

#### Solid-phase synthesis of oligonucleotides

DNA was synthesized on an Applied Biosystems Expedite 8909 nucleic acid synthesis system. Native oligonucleotides were synthesized as recommended by the manufacturer using standard β-cyanoethyl phosphoramidites. Oligonucleotides containing deazapurine nucleotides were synthesized using 0.5 M camphorsulfonyl oxaziridine in anhydrous acetonitrile as oxidant, with 3 min oxidation time. Coupling time for all modified phosphoramidites was 10 min. The final dimethoxytrityl groups were retained after synthesis to facilitate purification by Glen-Pak reversed-phase cartridge (Glen Research). Synthesized oligonucleotides were deprotected with ammonium hydroxide/methylamine (AMA) for 10 min at 65 °C in most cases; however, oligonucleotides synthesized using universal support were deprotected with AMA for 2 h at 65 °C, and oligonucleotides containing DBCO modifications were deprotected with concentrated ammonia for 2 h at 65 °C. Deprotected oligonucleotides were purified with Glen-Pak reversed-phase cartridges following the provided protocol, and the resulting crude products were dried by Thermo Fisher SpeedVac vacuum concentrator. Oligonucleotides < 30 nt were then purified via 20% denaturing polyacrylamide gel electrophoresis (PAGE), while those > 30 nt were purified with 12% denaturing PAGE. Oligonucleotides containing DBCO were not PAGE purified due to instability of the DBCO group. After electrophoresis, oligonucleotides were detected by UV shadowing or directly visualized if the oligonucleotides were fluorescently labeled. The desired bands were cut and eluted from the gel using the “crush and soak" technique. Finally, the eluent was desalted using C18-Sep-Pak cartridges (Waters) and dried by vacuum concentrator.

**Immobilization of oligonucleotides on magnetic carboxylic acid beads**

Dynabeads MyOne carboxylic acid (5 mg, 500 µL) was washed once with 500 µL of 10 mM NaOH and five times with 500 µL of nuclease-free water. The washed beads were resuspended in a 150 µL reaction mixture containing 200 mM NaCl, 200 µM 5'-amino-modified oligonucleotide, 1 mM imidazole, 50% v/v DMSO, and 250 mM EDC. Beads were mixed well with reagents, vortexed, sonicated and incubated overnight on a rotator at room temperature. After coupling the oligonucleotides, the beads were washed three times with cold 100 mM 2-(N-morpholino)ethanesulfonic acid (MES) buffer (pH 4.7). The unconjugated carboxylic acid groups on the beads were activated with 250 mM EDC and 100 mM NHS in MES buffer at room temperature for 30 min. Activated beads were washed three times with cold PBS buffer and passivated with 20 mM methyl-PEG12-amine in PBS buffer for 2 h at room temperature. The passivated beads were washed three times with 500 µL of Tris-Tween (TT) buffer (100 mM Tris-HCl, 0.01% v/v Tween-20, pH 7.5), and stored in 500 µL of PBS-Tween (PBST) buffer (1× PBS, pH 7.4, 0.01% v/v Tween-20).

**HPLC analysis of oligonucleotides**

Oligonucleotides (10 µM, 10 µL) were analyzed on an Agilent 1260 Infinity II HPLC equipped with a Waters XBridge oligonucleotide BEH C18 column (130Å, 2.5 µm, 4.6 mm × 50 mm). The mobile phase consisted of 100 mM triethylammonium acetate (A) and acetonitrile (B). The gradient began with 5% B from t = 0 to 2 min, ramped to 30% B linearly over 13 min, ramped to 97% B over 5 min, and was then held at 97% B from t = 20 to 24 min. Mobile phase B was reduced to 5% linearly over 2 min and kept at 5% for 8 min. The flow rate was 1 mL/min, and the column was maintained at ambient temperature. The oligonucleotides were monitored by the absorbance at 260 nm and 284 nm.

**Investigation of DNA stability during $BF_3 \cdot Et_2O$ treatment**

Amino-modified oligonucleotide **ODN-S3** was immobilized onto 500 µL (5 mg) of Dynabeads MyOne carboxylic acid beads. The beads were divided into 100 µL aliquots, and each aliquot was incubated with oligonucleotide **ODN-1** or **ODN-2** (10 µM, 100 µL) in PBS-Tween (PBST) buffer (1× PBS, pH 7.4, 0.01% v/v Tween-20) for 10 min at room temperature. After hybridization, the beads were washed three times with 100 µL of PBST buffer to remove excess DNA, and then washed five times with 100 µL of PBST plus acetonitrile (50% v/v), and five times with anhydrous acetonitrile. The beads were dried under vacuum in a desiccator for 30 min. The dried beads were incubated in 100 µL of 40 mM $BF_3 \cdot Et_2O$ in anhydrous acetonitrile at 50 °C for up to 4 h. After incubation, the beads were washed five times with 100 µL of anhydrous acetonitrile, and once with 50 µL of 20 mM NaOH to neutralize excess $BF_3 \cdot Et_2O$. The washed beads were then incubated with 100 µL of 20 mM NaOH at room temperature for 10 min to release the oligonucleotides. The oligonucleotides were then purified by ethanol precipitation (0.3 M NaOAc in 70% ethanol), resuspended in nuclease-free water, and analyzed by HPLC (Agilent 1260 infinity II HPLC) and MALDI-TOF (Bruker Microflex MALDI-TOF). A similar procedure was performed with beads modified with amino-modified oligonucleotide **ODN-S4** and oligonucleotide **ODN-S1** and **ODN-S2**.

**Primer extension with $c^7$dATP and $c^7$dGTP**

A typical 20 μL primer extension reaction contained 1.5 μM template (**ODN-S5**) and 1 μM Cy3-labeled primer (**ODN-S6** or **-S7**). We tested different polymerases using the following conditions (conditions are shown for **ODN-S6**; for **ODN-S7** reactions, $c^7$dATP and $c^7$dGTP were replaced with dATP and dGTP):

*Sequenase version 2.0 DNA polymerase*: 0.5 U polymerase, 300 μM $c^7$dATP, 300 μM dTTP, 300 μM $c^7$dGTP, 300 μM dCTP, 40 mM Tris-HCl (pH 7.5), 10 mM $MgCl_2$, and 5 mM DTT, incubated for 15 or 30 min at 37 °C.

*Klenow Fragment (3'→5' exo-)*: 1 U polymerase, 300 μM $c^7$dATP, 300 μM dTTP, 300 μM $c^7$dGTP, 300 μM dCTP, 10 mM Tris-HCl (pH 7.9), 50 mM NaCl, 10 mM $MgCl_2$, and 1 mM DTT, incubated for 15 or 30 min at 37 °C.

*Bst 3.0 DNA Polymerase*: 1 U polymerase, 300 μM $c^7$dATP, 300 μM dTTP, 300 μM $c^7$dGTP, 300 μM dCTP, 20 mM Tris-HCl (pH 8.8), 10 mM $(NH_4)_2SO_4$, 150 mM KCl, 2 mM $MgSO_4$, and 0.1% Tween-20, incubated for 15 or 30 min at 55 °C.

After primer extension, the DNA was cleaned by ethanol precipitation (0.3 M NaOAc, 70% ethanol), and analyzed by denaturing PAGE. The gel was imaged on an Amersham Typhoon RGB imager.

**Functionalization of amino-modified oligonucleotides with NHS ester**

Amino-modified oligonucleotide (100 nmol) was dissolved in 250 μL of 100 mM sodium bicarbonate buffer (pH 8.2). NHS ester (150 mM, 25 μL) in anhydrous DMSO was added to the oligonucleotide, and the reaction was incubated overnight on a rotator at room temperature. The conjugated oligonucleotide was then separated from excess salt and NHS ester by ethanol precipitation (0.3 M NaOAc, 70% ethanol).

**DNA-encoded Edman degradation**

We began by preparing oligonucleotide-modified magnetic beads. The amino-modified oligonucleotide **ODN-3** was immobilized onto 100 μL (1 mg) of Dynabeads MyOne carboxylic acid beads. We then ligated 10 μM 5'-phosphorylated, DBCO-modified DNA **ODN-4** in a 100 μL reaction containing 10 μM splint DNA **ODN-S9**, T4 ligase (2,000 U), 50 mM Tris-HCl (pH 7.5), 10 mM $MgCl_2$, 1 mM ATP, and 0.01% Tween-20 at room temperature for 2 h. After ligation, splint DNA was removed by incubation with 100 μL of 20 mM NaOH for 10 min, after which the beads were washed three times with PBST buffer.

Next, the beads were incubated with 10 mM peptide FGGGGGX (X = azidolysine) dissolved in 100 μL of PBST at room temperature for 2 h. After the completion of the SPAAC reaction, the beads were washed with PBST buffer three times to remove unconjugated peptide, and then washed with 100 μL of PITC conjugation solution (pyridine/ethanol/water, 2:1:1 v/v). Azide-modified PITC (**1**) (200 mM, 100 μL) in conjugation solution was added to the beads, and the peptides were allowed to react for 30 min at 50 °C. The beads were then washed three times with 100 μL of conjugation solution and three times with 100 μL of PBST-Mg buffer (PBST plus 2 mM $MgCl_2$, pH 7.4). Next, proximity SPAAC was carried out by incubating the washed beads with 20 μM primer **ODN-5** in 100 μL of PBST-Mg buffer for 10 min at room temperature.

For the primer extension step, the beads were washed three times with 100 μL of primer extension buffer (50 mM NaCl, 10 mM Tris-HCl, 10 mM $MgCl_2$, 0.01% v/v tween-20, pH 7.5). We then added 60 μL primer extension mixture to the beads, which included Klenow fragment exo-) (8 U), 300 μM $c^7$dATP, 300 μM dTTP, 300 μM $c^7$dGTP, 300 μM dCTP, 50 mM NaCl, 10 mM Tris-HCl (pH 7.5), 10 mM $MgCl_2$, and 0.01% v/v tween-20, with incubation for 1 h at 37 °C. The beads were then washed three times with 100 μL of PBST buffer to remove excess reagents.

For the cleavage step, the beads were washed five times with 100 μL of 50% acetonitrile (v/v) in PBST and five times with anhydrous acetonitrile. The beads were dried under vacuum in a desiccator for 30 min, and then incubated in 100 μL of 40 mM $BF_3 \cdot Et_2O$ in anhydrous acetonitrile at 50 °C for 5 min. The beads were then washed five times with 100 μL of anhydrous acetonitrile, and once with 50 μL of basic conversion solution (0.25% v/v $NH_4OH$, 200 μM DTT) to neutralize excess $BF_3 \cdot Et_2O$. The washed beads were then incubated with 100 μL of basic conversion solution at 50 °C for 10 min to release the DNA-barcoded PTC amino acid. The DNA-barcoded PTC amino acid was cleaned by ethanol precipitation (0.3 M NaOAc in 70% ethanol). The DNA-barcoded PTC amino acid was resuspended in HPLC running buffer (10 mM $NH_4OAc$ in 80% acetonitrile) and analyzed by HPLC-MS (see below). The beads were washed three times with PBST buffer and stored in PBST buffer.

**Isolation and analysis of intermediates of DNA-encoded Edman degradation**

Amino-modified anchor DNA containing a disulfide linker and a Cy3 modification (**ODN-S10**), was immobilized onto 500 μL (5 mg) of Dynabeads MyOne carboxylic acid beads as described above. A 50 μL aliquot of these beads were treated with 50 μL of 10 mM TCEP in PBST buffer at room temperature for 30 min to cleave the oligonucleotide off the bead surface. The released oligonucleotide was purified by ethanol precipitation (0.3 M NaOAc in 70% ethanol). The rest of the beads were subjected to the procedure described above, with 50 μL aliquots taken after DNA ligation, conjugation of peptide, conjugation of azide-modified PITC, and conjugation of DBCO-modified primer. All aliquots were reduced with 50 μL of 10 mM TCEP, and the released intermediates were cleaned by ethanol precipitation (0.3 M NaOAc in 70% ethanol). All intermediates were analyzed by denaturing PAGE, and the gel was imaged by an Amersham Typhoon RGB imager.

**Time-course experiment of proximity SPAAC for peptides of varying length and rigidity**

Dynabeads MyOne carboxylic acid beads (5 mg, 500 μL) were modified with oligonucleotides as described in the DNA-encoded Edman degradation procedure. Beads were divided into 100 μL aliquots and conjugated to peptides ranging from 10–30 aa with varying rigidity (see **Table S3**) as described previously, after which the peptides were modified with azide-modified PITC and subjected to proximity SPAAC with **ODN-S13**. As a no-proximity negative control, we performed a reaction using beads modified with the shortest peptide GGGGSGGGGSX and **ODN-S14**. The reactions were incubated at room temperature for 2 h, with 1 μL aliquots collected at 10 min, 30 min, 1 h, and 2 h. These aliquots were washed with 100 μL of 20 mM NaOH for 10 min to remove unreacted DBCO-modified DNA, and then resuspended in PBST and analyzed on a NovoCyte Quanteon flow cytometer.

**Quantification of degradation yield**

After completion of the primer conjugation via proximity SPAAC during the Edman degradation procedure above, we collected a 0.2 μL aliquot from the reaction mixture and incubated it with 20 μL of 10 μM Cy3-labeled complementary strand of the primer (**ODN-S12**) for 10 min at room temperature. The beads were washed three times with PBST buffer to remove excess Cy3-labeled strand, and then analyzed by a NovoCyte Quanteon flow cytometer. The mean fluorescence signal in PE channel was recorded as [mean PE-A before degradation]. Similarly, after the release of the DNA-barcoded PTC amino acid, we removed a 0.2 μL aliquot from the reaction mixture and incubated with **ODN-S12**. The mean fluorescence signal in the PE channel was designated as [mean PE-A after degradation]. The percent yield of the cleavage reaction was determined as: $100\% - \frac{[\text{mean PE−A after degradation}]}{[\text{mean PE−A before degradation}]} \times 100\%$.

**LC-MS analysis of DNA-barcoded PTC amino acids**

The DNA-barcoded PTC amino acid (10 μM, 5 μL) was analyzed by LC-ESI/MS on a Waters Acquity UPLC and a Thermo Exploris 240 BioPharma Orbitrap MS. The UPLC was equipped with a Waters Acquity UPLC BEH Amide Column (130 Å, 1.7 μm, 2.1 mm × 100 mm). The mobile phase consisted of 10 mM ammonium acetate in 80% acetonitrile (A) and 10 mM ammonium acetate in 25% acetonitrile (B). The gradient began with 10% B from t = 0 to 1 min, ramped to 50% B linearly over 2 min, ramped to 90% B over 7 min, and was then held at 90% B from t = 10 to 14 min. The flow rate was 0.3 mL/min, and the column was maintained at 60 °C. Full-scan MS spectra were acquired over the mass range 570–4,000 m/z with 120,000 resolution. The molecular weight of the DNA-barcoded PTC amino acids was obtained after deconvolution using Protein Metrics Byos software.

**Preparation of PTC amino acid-conjugated BSA**

BSA (1 mg) was dissolved in 270 μL of 100 mM sodium bicarbonate buffer (pH 8.2). TFP Ester-PEG4-DBCO (70 mM, 30 μL) in anhydrous DMSO was added to the BSA solution, and the reaction was incubated for 2 h on a rotator at room temperature. After conjugation, DBCO-modified BSA was purified by buffer exchange to 1× PBS using a Zeba spin desalting column (7K MWCO), yielding a final concentration of ~3 mg/mL. The degree of labeling was determined via SDS-PAGE and UV-Vis (Thermo Fisher Scientific Nanodrop 2000). PTC-amino acid azide (100 mM, 15 μL) in DMSO was added to 300 μL (~1 mg) of DBCO-modified BSA in 1× PBS and incubated for 1 h on a rotator at room temperature. Another 15 μL aliquot of 100 mM PTC-amino acid azide in DMSO was then added, and the reaction was incubated for an additional hour. After conjugation, the PTC amino acid-conjugated BSA was purified by buffer exchange to 1× PBS using a Zeba spin desalting column (7K MWCO), yielding a final concentration of ~3 mg/mL. The degree of labeling was again determined as described above. Completion of the click reaction was marked by the disappearance of DBCO absorbance at 310 nm.

**Immunization, hybridoma generation, and antibody purification**

Immunization and hybridoma generation was carried out by Antibody Solutions. Each BSA-conjugated PTC amino acid was administered to a cohort of three female CD1 mice as twice-weekly subcutaneous injections to a single hind footpad for four weeks. All injections consisted of 10 μL (10 μg) of BSA-conjugated PTC amino acid in 1× PBS buffer mixed with an equal volume of Sigma Adjuvant System. On day 21, serum from each mouse was evaluated by ELISA for binding to its corresponding target. In short, ELISA plates were coated with each respective target, blocked, and then incubated with serial dilutions of each serum sample. Plates were washed and then incubated with HRP-conjugated Mouse IgG-specific secondary antibody, followed by another wash step, development with TMB substrate, and finally absorbance measurement at 450 nm. The specificity of each serum sample was evaluated in parallel against duplicate wells coated with the respective target antigen. Before the serum samples were added to the ELISA plates, each sample was pre-incubated with an excess amount of other BSA-conjugated PTC amino acids to block antibodies against the BSA carrier or conserved moieties (e.g., the $PEG_4$ linker in azide-modified PITC and DBCO). On day 28, selected mice for each PTC amino acid received a final boost injection. Three days after the final injection, popliteal and inguinal lymph nodes were harvested from selected mice within each cohort and teased to isolate lymphocytes. These were then mixed with P3X63Ag8.653 myeloma cells lacking hypoxanthine-guanine phosphoribosyltransferase (HGPRT). Cell fusions were performed via the traditional PEG method by Kohler and Milstein. The cells were then cultured for 7 days in hypoxanthine-aminopterin-thymidine medium to select for hybridoma cells capable of growth in the presence of aminopterin while eliminating unfused lymphocytes and myeloma cells. On day 7, the cells were transitioned to hypoxanthine-thymidine (HT) medium and cultured for four more days. For each PTC amino acid target, the resulting population of hybridomas was split into six aliquots and cryopreserved to generate a hybridoma library, and the hybridoma-conditioned HT medium was harvested to evaluate the binding performance of the secreted antibodies by ELISA as described above.

To isolate monoclonal hybridomas, one vial of each hybridoma library was thawed, and single cells were sorted by flow cytometry into ten 96-well culture plates. After two weeks in culture, supernatants from all ten plates were evaluated by ELISA for binding to the respective target antigen. Positive clones were identified and further characterized by ELISA against a panel of other PTC-modified amino acid targets to confirm specificity. Clones selective for each target were expanded from 96-well plates into T-25 culture flasks with HT medium. Once cells reached confluence, each clonal line was cryopreserved in two aliquots as was the conditioned hybridoma supernatant from each line. For selected monoclonal hybridomas, the secreted antibody was purified by protein A chromatography.

**Measurement of antibody affinity**

The biotinylated DBCO modified oligonucleotide (50 nmol, **ODN-S16**) was dissolved in 100 µL of 1× PBS buffer. 10 µL of 100 mM PTC amino acid in DMSO was added to the oligonucleotide, and the reaction was incubated for 2 h on a rotator at room temperature. After conjugation, the conjugated oligonucleotide was purified by ethanol precipitation (0.3 M NaOAc in 70% ethanol). Biolayer interferometry (BLI) experiments were carried out on an Octet RED384 instrument. Briefly, SA biosensors were equilibrated in PBST buffer for 10 min. The biotinylated DNA-tagged PTC amino acid (100 nM) was loaded onto the SA biosensors for 150 s. The reference biosensors were incubated with PBST buffer alone. All biosensors were washed with PBST and then incubated with varying concentrations of anti-PTC amino acid antibodies in PBST buffer for 800 s. Finally, the associated antibodies were allowed to dissociate in PBST buffer for 150 s. The responses after reference subtraction were plotted against concentration and fitted to a Langmuir binding isotherm using GraphPad Prism 10.0 to determine the antibody dissociation constant ($K_d$).

**Preparation of primer-modified anti-PTC amino acid antibodies**

oYo-Link tetrazine was received as dry pellet and resuspended in 100 µL of nuclease-free water. This solution was mixed with 100 µL of 1 mg/mL antibody in 1× PBS, and then irradiated by UV (365 nm) on ice for 2 h. After UV irradiation, excess oYo-Link tetrazine was removed by filtering with an Amicon centrifugal filter (50 kDa MWCO). The tetrazine-modified antibody (~1 mg/mL, 100 μL) was mixed with 10 μL of 300 μM TCO-modified primer **ODN-7**, **ODN-8**, or **ODN-9**, and the reaction was incubated for 2 h at room temperature. After conjugation, excess primer was removed by filtering with an Amicon centrifugal filter (50 kDa MWCO). The concentrations of primer-antibody conjugate were determined by UV-Vis on a NanoDrop spectrophotometer, and the degree of labeling was determined by SDS-PAGE.

**Proximity extension assay (PEA)**

The model PEA template (50 nmol, **ODN-6**) was dissolved in 100 μL of 1× PBS buffer. Next, 10 μL of 100 mM PTC amino acid in DMSO was added to the oligonucleotide and the mixture was incubated for 2 h on a rotator at room temperature. After conjugation, the conjugated oligonucleotide was purified by ethanol precipitation (0.3 M NaOAc in 70% ethanol). Dynabeads MyOne streptavidin C1 beads (1 mg) were washed twice with 1× binding and washing (B&W) buffer (5 mM Tris-HCl, 0.5 mM EDTA, 1 M NaCl, pH 7.5). Then, 100 μL of 100 nM biotinylated DNA-tagged PTC amino acids in 1× B&W buffer was incubated with the washed beads on a rotator for 2 h at room temperature, and then washed three times with PBST buffer. DNA-loaded beads (0.01 mg) was incubated with 10 μL of 200 nM primer-modified anti-PTC amino acid antibody in 50 mM NaCl, 10 mM Tris-HCl, 10 mM $MgCl_2$, and 0.01% v/v Tween-20 (pH 7.5) for 30 min at room temperature. We next added 10 μL of mixture containing Klenow fragment (exo-) (1 U), 600 μM dATP, 600 μM dTTP, 600 μM dGTP, 600 μM dCTP, 50 mM NaCl, 10 mM Tris-HCl (pH 7.5), 10 mM $MgCl_2$, and 0.01% v/v Tween-20, and performed primer extension for 30 min at 37 °C. After extension, beads were washed three times with PBST and resuspended in 20 μL PBST.

**Quantification of DNA output of PEA by qPCR**

A 50 μL PCR reaction was prepared with 1 μL (0.5 μg) of beads after the PEA step and 2× GoTaq qPCR master mix. Final concentrations of forward and reverse primers (**ODN-S17** and **ODN-S18**) were 1 μM. Reactions were run on a Bio-Rad CFX96 real-time PCR detection system using the following program: 3 min at 95 °C, and 40 cycles of 10 s at 95 °C and 1 min at 55 °C. The Cq values were determined using Bio-Rad CFX Manager software. Negative control reactions were performed without beads.

**Sequencing of a single peptide**

We generally followed the Edman procedure described above, with the following adjustments. To begin, 1 mg of Dynabeads MyOne carboxylic acid beads were modified with anchor DNA (**ODN-11**), and DBCO-modified DNA (**ODN-12**) was then conjugated by DNA ligation. Peptide RGFDWGX was subsequently conjugated via SPAAC. The subsequent proximity SPAAC reaction was carried out with DBCO-modified primer (**ODN-13**). After each cycle, the PTC amino acid was precipitated and processed separately. PTC amino acids were resuspended in 1 mL of 1× B&W buffer and a 100 μL aliquot was incubated with 1 mg of Dynabeads MyOne streptavidin C1 beads on a rotator for 2 h at room temperature. Subsequently, PEA was carried out with a mixture of DNA-barcoded (**Table S4**) anti-PTC-R, -F, -D, and -W antibodies (100 nM each) using the procedure described above.

For each individual PEA reaction, a 100 μL adaptor PCR reaction was prepared with 2 μL (1 μg) of the post-PEA beads and 2× colorless GoTaq G2 hot start master mix. The final concentration of forward and reverse adaptor primers (**Table S1**) was 1 μM each, where the forward primer contained the cycle number barcode. Reactions were run on an Eppendorf Mastercycler X50 PCR

Thermocycler using the following program: 2 min at 95 °C, 8 cycles of 15 s at 95 °C, 15 s at 54 °C and 30 s at 72 °C. The adaptor PCR reaction was cleaned up using an Axygen AxyPrep MAG PCR clean-up kit. PCR products from all peptide sequencing cycles were combined and indexed with the Nextera XT index kit v2. The indexed library was sequenced on an Ilumina MiSeq sequencing system. The sequencing data was processed with Galaxy. Briefly, barcodes for cycle number, antibody, and peptide were extracted using the "Trim sequences" tool. The extracted barcodes were joined to give a single barcode using the "FASTQ joiner" tool, and the counts of each barcode were counted with "Collapse" tool.

**Probing barcode crosstalk during PEA**

A 1:1 mixture of the 42 nt template coupled to PTC-F **ODN-6-PTC-F** and the 60 nt template **ODN-S33** (total DNA concentration = 1 μM) was prepared and immobilized onto SA beads at densities ranging from 0.5–100 pmol per mg of beads. PEA reactions were carried out with 0.01 mg of modified SA beads using anti-PTC-F antibody conjugated with the 6 nt PEA primer **ODN-8** as described previously. After PEA, beads were resuspended in 20 μL PBST. A 50 μL PCR reaction was prepared with 1 μL (0.5 μg) of beads and 2× GoTaq Master Mix plus a 1 μM final concentration of forward and reverse primers **ODN-S17** and **ODN-S18**. Reactions were run on an Eppendorf Mastercycler X50 PCR thermocycler using the following program: 2 min at 95 °C, 8 cycles of 15 s at 95 °C, 15 s at 54 °C and 30 s at 72 °C. The products were analyzed by native PAGE. The gel was stained with GelStar nucleic acid gel stain and imaged by a Bio-Rad Gel Doc XR+ gel imaging system.

**Probing barcode crosstalk during proximity SPAAC**

A 1:1 mixture of two DBCO-modified anchor DNA sequences (**ODN-S32** and **ODN-S33**, total DNA concentration = 200 μM) was immobilized onto carboxylic acid beads as described previously. We performed a proximity SPAAC reaction on 1 mg of modified beads with 10 μM azide-modified complementary strands **ODN-S34** and **ODN-S35** in 100 μL PBST-Mg buffer (PBST plus 2 mM $MgCl_2$, pH 7.4) for 10 min at room temperature. Control experiments were carried out using beads only modified with **ODN-S32** reacting to **ODN-S34**, or beads only modified with **ODN-S33** reacting to **ODN-S35**. The beads were then washed with 100 μL PBST buffer three times and treated with 50 μL of 10 mM TCEP, and the released DNA was cleaned by ethanol precipitation (0.3 M NaOAc, 70% ethanol). DNA was analyzed by 10% native PAGE, and the gel was imaged by a Bio-Rad Gel Doc XR+ gel imaging system.

**Parallel sequencing of multiple peptide species**

To sequence multiple peptides simultaneously, we made the following modifications to the procedure for peptide immobilization. First, Dynabeads MyOne carboxylic acid (1 mg) were modified with anchor DNA (**ODN-11**). Two azide-modified peptides, AFGGGX and AWGGGX, were clicked with DBCO-modified peptide-barcoding strand **ODN-12** and **ODN-S19**, respectively. For Fig. 5, E to G, the azide-modified peptides $^{p}$YGYGGX, YG$^{p}$YGGX, and $^{p}$YG$^{p}$YGGX were clicked with DBCO-modified peptide-barcoding strand **ODN-12**, **ODN-S19**, and **ODN-S20** respectively. The peptide-DNA conjugates were mixed and immobilized on beads via DNA ligation. After each cycle of Edman degradation, a pulldown was carried out with a 10 μL aliquot of DNA-barcoded PTC amino acids and 2 mg of Dynabeads MyOne streptavidin C1 beads to minimize barcode crosstalk during PEA. For Fig. 5, C and D, PEA was carried out with a mixture of DNA-barcoded anti-PTC-F and -W antibodies (30 nM each). For Fig. 5, E to G, two separate PEA reactions were performed: one with PY20 (30 nM), and the other with anti-PTC-Y antibody (100 nM). The rest of the procedure was identical to that used to sequence a single peptide species.

**Synthesis of organic compounds**

*General synthetic procedures.* All chemicals purchased from commercial suppliers were used without further purification. The purities of compound were determined by nuclear magnetic resonance (NMR) with an Agilent 400 NMR spectrometer and high-resolution mass spectrometry (HRMS) were obtained by LC-ESI/MS with a Waters Acquity UPLC equipped with an Agilent Zorbax SB-C18 column (2.1 × 50 mm, 2.7 µm) and Thermo Exploris 240 BioPharma orbitrap mass spectrometer.

**Scheme S1. Synthesis of azide-modified PITC (1)**

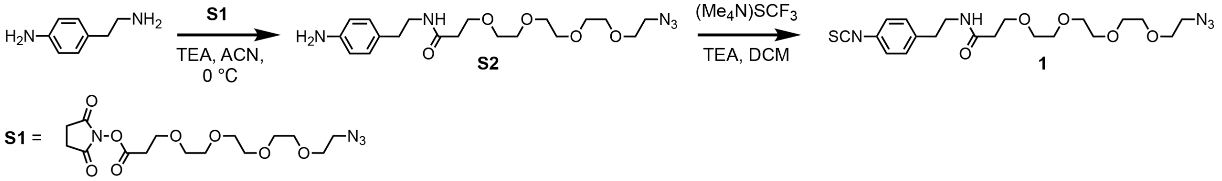


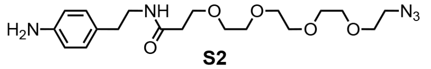

S2

Under nitrogen atmosphere, 4-(2-aminoethyl)aniline (389 mg, 2.86 mmol), acetonitrile (20 mL), and triethylamine (261 mg, 360 µL, 2.6 mmol) were added to a dry round bottom flask. The reaction was stirred on an ice bath, and azido-PEG4-NHS ester (**S1**, 1 g, 2.6 mmol) dissolved in ACN (5 mL) was added dropwise. The reaction was kept in the ice bath which slowly warmed up to room temperature over the course of the reaction. The progress of the reaction was monitored by thin-layer chromatography (TLC), and the reaction was completed in 1 h. The reaction was quenched by water and extracted with DCM (3 × 50 mL). The combined organic phase was washed with brine, dried over $Na_2SO_4$, filtered, and concentrated under reduced pressure. The mixture was purified by flash chromatography on a silica column. Elution with 5% MeOH in DCM gave **S2** as light yellow oil (1.05 g, 98%). $^1$H NMR (400 MHz, $CDCl_3$) δ 7.11–6.84 (m, 2H), 6.73–6.61 (m, 2H), 6.53 (s, 1H), 3.74–3.49 (m, 16H), 3.46–3.24 (m, 4H), 2.67 (t, $J$ = 7.1 Hz, 2H), 2.41 (t, $J$ = 5.8 Hz, 2H). $^{13}$C NMR (101 MHz, $CDCl_3$) δ 171.47, 144.25, 129.57, 129.24, 115.60, 70.64, 70.62, 70.56, 70.46, 70.21, 70.17, 70.00, 67.28, 53.48, 50.62, 40.71, 36.88, 34.73. HRMS $C_{19}H_{32}N_5O_5^+$ $(M + H)^+$: calculated $m/z$ = 410.2398, found $m/z$ = 410.2394.

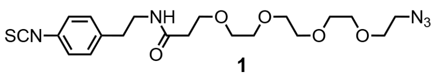

1

Isothiocyanate **1** was synthesized using a previously reported method.[66] Under nitrogen atmosphere, compound **S2** (1.05 mg, 2.57 mmol), DCM (16 mL), and TEA (394 mg, 540 µL, 3.9

mmol) were added to a dry round bottom flask. The reaction was stirred at room temperature, and $(Me_4N)SCF_3$ (501 mg, 2.86 mmol) was added. After 1 h, 10 mL of hexanes was added to the reaction mixture, which was then filtered through a pad of celite. The celite pad was subsequently washed with 10 mL of a 1:1 mixture of DCM/hexanes. The filtrate was concentrated under reduced pressure. The mixture was purified by flash chromatography on a silica column. Elution with 3% MeOH in DCM gave **1** as light orange oil (1.10 g, 95%). $^1$H NMR (400 MHz, $CDCl_3$) δ 7.24–7.10 (m, 4H), 6.73 (s, 1H), 3.78–3.50 (m, 16H), 3.46 (td, *J* = 7.1, 5.9 Hz, 2H), 3.42–3.25 (m, 2H), 2.80 (t, *J* = 7.1 Hz, 2H), 2.44 (dd, *J* = 6.1, 5.2 Hz, 2H). $^{13}$C NMR (101 MHz, $CDCl_3$) δ 171.89, 138.74, 134.92, 129.99, 129.32, 125.77, 70.67, 70.62, 70.54, 70.47, 70.19, 70.13, 70.03, 67.14, 53.47, 50.63, 40.31, 36.70, 35.24. HRMS C20H30N5O5S$^+$ (M + H)$^+$: calculated *m/z* = 452.1962, found *m/z* = 452.1958.

**Scheme S2. Synthesis of PTC amino acid S3-S14**

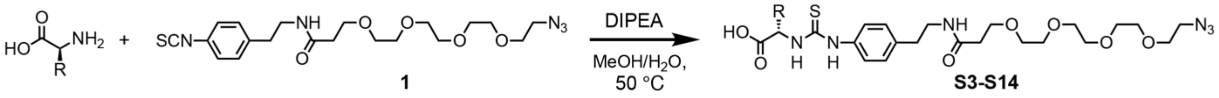


*General procedure for synthesizing PTC amino acids*: Under nitrogen atmosphere, isothiocyanate **1** (90.2 mg, 0.2 mmol) was added to a round bottom flask and dissolved in MeOH/water (1:1, v/v, 4 mL). Amino acid (0.21 mmol) and DIPEA (39.3 mg, 53 µL, 0.3 mmol) were added, and the reaction was stirred at 50 °C for 30 min. The reaction was concentrated under reduced pressure and redissolved in MeOH/DCM (1:1, 1 mL). The mixture was purified with preparative TLC (60 Å silica gel plate, with UV254 indicator, glass backed, 250 µm, 20 × 20cm) using the developing solvent indicated below.

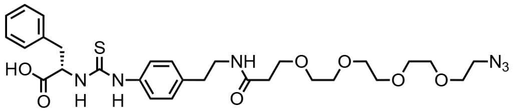

**S3**

*PTC-F:* Preparative TLC was developed with 20% MeOH in DCM. PTC-F (**S3**) was isolated as light yellow foam (97.2 mg, 79%). $^1$H NMR (400 MHz, MeOH-d4) δ 7.38–6.97 (m, 9H), 3.83–3.52 (m, 16H), 3.47–3.33 (m, 6H), 3.14 (dd, *J* = 13.9, 5.8 Hz, 1H), 2.76 (t, *J* = 7.2 Hz, 2H), 2.39 (t, *J* = 6.0 Hz, 2H). $^{13}$C NMR (101 MHz, MeOH-d4) δ 180.00, 172.64, 136.91, 136.87, 136.03, 129.75, 129.36, 129.25, 128.93, 128.22, 128.07, 128.02, 126.43, 124.14, 70.12, 70.04, 69.95, 69.88, 69.68, 66.91, 58.41, 40.53, 40.40, 36.75, 36.20, 36.15, 34.52. HRMS $C_{29}H_{39}N_6O_7S^-$ (M − H)$^-$: calculated *m/z* = 615.2606, found *m/z* = 615.2607.

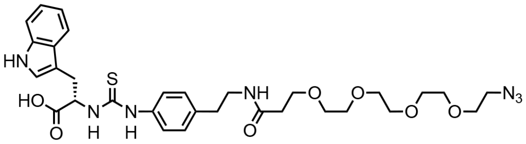

**S4**

*PTC-W:* Preparative TLC was developed with 20% MeOH in DCM. PTC-W (**S4**) was isolated as light yellow foam (106.3 mg, 81%). $^1$H NMR (400 MHz, MeOH-d4) δ 7.58 (d, *J* = 7.9

Hz, 1H), 7.33 (d, $J$ = 8.1 Hz, 1H), 7.19–6.71 (m, 7H), 3.78–3.48 (m, 17H), 3.33 (h, $J$ = 7.6 Hz, 6H), 2.70 (t, $J$ = 7.1 Hz, 2H), 2.35 (t, $J$ = 5.9 Hz, 2H). $^{13}$C NMR (101 MHz, MeOH-d4) δ 179.62, 172.66, 136.89, 136.47, 135.75, 129.32, 127.98, 124.18, 123.22, 121.02, 118.52, 118.39, 110.86, 109.36, 69.98, 69.91, 69.82, 69.79, 69.59, 66.92, 58.61, 40.44, 40.32, 36.00, 34.46, 29.35, 26.71. HRMS $C_{31}H_{40}N_7O_7S^-$ $(M - H)^-$: calculated $m/z$ = 654.2715, found $m/z$ = 654.2717.

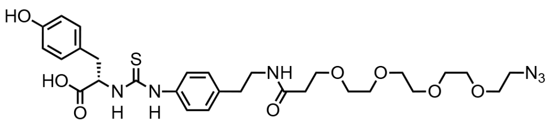


**S5**

*PTC-Y:* Preparative TLC was developed with 20% MeOH in DCM. PTC-Y (**S5**) was isolated as light yellow foam (105.5 mg, 83%). $^{1}$H NMR (400 MHz, MeOH-d4) δ 7.14 (q, $J$ = 8.2 Hz, 4H), 7.00 (d, $J$ = 8.0 Hz, 2H), 6.69 (d, $J$ = 8.1 Hz, 2H), 3.89–3.50 (m, 17H), 3.42–3.34 (m, 5H), 3.05 (dd, $J$ = 13.9, 5.4 Hz, 1H), 2.76 (t, $J$ = 7.1 Hz, 2H), 2.39 (t, $J$ = 5.7 Hz, 2H). $^{13}$C NMR (101 MHz, MeOH-d4) δ 179.67, 172.71, 155.93, 136.99, 135.91, 130.79, 130.27, 129.44, 128.93, 127.54, 124.21, 114.78, 70.04, 69.96, 69.86, 69.64, 66.95, 58.84, 50.27, 40.51, 40.38, 36.05, 35.92, 34.50. HRMS $C_{29}H_{39}N_6O_8S^-$ $(M - H)^-$: calculated $m/z$ = 631.2555, found $m/z$ = 631.2559.

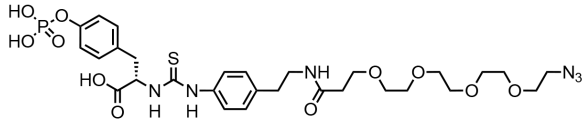


**S6**

*PTC-$^{p}$Y:* Preparative TLC was developed with 30% MeOH in DCM. PTC-$^{p}$Y (**S6**) was isolated as light yellow foam (91.7 mg, 64%). $^{1}$H NMR (400 MHz, MeOH-d4) δ 7.14 (d, $J$ = 19.0 Hz, 8H), 3.70–3.55 (m, 17H), 3.49–3.42 (m, 1H), 3.42–3.34 (m, 4H), 3.20–3.13 (m, 1H), 3.13–3.04 (m, 1H), 2.76 (t, $J$ = 7.1 Hz, 2H), 2.41 (t, $J$ = 5.9 Hz, 2H).$^{13}$C NMR (101 MHz, MeOH-d4) δ 179.02, 175.38, 172.63, 152.15, 152.08, 136.73, 136.06, 131.86, 130.03, 129.42, 123.95, 119.75, 119.70, 70.02, 69.99, 69.96, 69.92, 69.85, 69.76, 69.61, 66.94, 59.59, 50.31, 40.35, 36.05, 34.45. $^{31}$P NMR (162 MHz, DMSO-d6) δ -5.04. HRMS $C_{29}H_{40}N_6O_{11}PS^-$ $(M - H)^-$: calculated $m/z$ 711.2218, found $m/z$ = 711.2220.

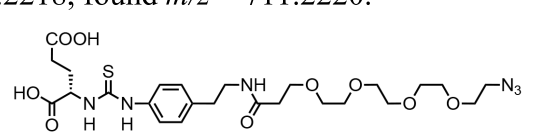


**S7**

*PTC-E:* Preparative TLC was developed with 30% MeOH in DCM. PTC-E (**S7**) was isolated as white solid (97.8 mg, 82%). $^{1}$H NMR (400 MHz, MeOH-d4) δ 7.31 (t, $J$ = 9.5 Hz, 2H), 7.27–7.16 (m, 2H), 3.64 (qd, $J$ = 14.5, 4.5 Hz, 17H), 3.47–3.34 (m, 4H), 3.21 (q, $J$ = 7.4 Hz, 1H), 2.79 (t, $J$ = 7.1 Hz, 2H), 2.54–2.28 (m, 4H), 2.09 (d, $J$ = 14.8 Hz, 1H). $^{13}$C NMR (101 MHz, MeOH-d4) δ 179.74, 172.63, 136.74, 129.35, 129.04, 128.77, 128.44, 124.15, 118.90, 70.02, 69.99, 69.96, 69.92, 69.86, 69.78, 69.61, 66.93, 58.25, 50.31, 40.40, 36.06, 34.53, 30.30, 27.97. HRMS $C_{25}H_{37}N_6O_9S^-$ $(M - H)^-$: calculated $m/z$ = 597.2348, found $m/z$ = 597.2350.

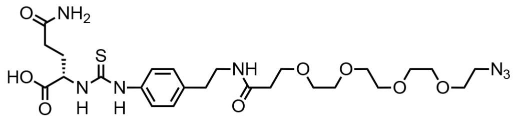


**S8**

*PTC-Q:* Preparative TLC was developed with 25% MeOH in DCM. PTC-Q (**S8**) was isolated as yellow solid (101.2 mg, 85%). $^{1}$H NMR (400 MHz, MeOH-d4) δ 7.34 (d, *J* = 7.9 Hz, 2H), 7.24 (d, *J* = 7.8 Hz, 2H), 3.82–3.53 (m, 17H), 3.41 (t, *J* = 6.4 Hz, 2H), 3.37 (t, *J* = 4.8 Hz, 2H), 3.21 (q, *J* = 7.4 Hz, 1H), 2.79 (t, *J* = 7.1 Hz, 2H), 2.41 (t, *J* = 6.0 Hz, 2H), 2.30 (s, 3H), 2.08 (s, 1H). $^{13}$C NMR (101 MHz, MeOH-d4) δ 180.14, 176.88, 172.69, 136.80, 136.39, 129.35, 124.19, 70.13, 70.06, 69.97, 69.87, 69.67, 66.90, 57.94, 50.34, 40.56, 40.44, 36.24, 36.20, 34.56, 31.26, 28.31. HRMS $C_{25}H_{38}N_7O_8S^-$ (M − H)$^-$: calculated *m/z* = 596.2508, found *m/z* = 596.2512.

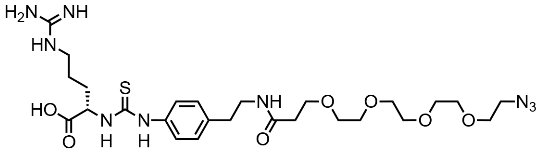


**S9**

*PTC-R:* Preparative TLC was developed with 30% MeOH in DCM. PTC-R (**S9**) was isolated as white foam (103.6 mg, 83%). $^{1}$H NMR (400 MHz, MeOH-d4) δ 7.32 (d, *J* = 8.0 Hz, 2H), 7.21 (d, *J* = 8.0 Hz, 2H), 3.76–3.52 (m, 17H), 3.45–3.33 (m, 4H), 3.20 (d, *J* = 7.5 Hz, 2H), 2.77 (t, *J* = 7.1 Hz, 2H), 2.41 (t, *J* = 6.1 Hz, 2H), 2.13–1.96 (m, 1H), 1.88 (t, *J* = 6.0 Hz, 1H), 1.63 (s, 2H). $^{13}$C NMR (101 MHz, MeOH-d4) δ 179.29, 177.23, 172.56, 157.21, 136.62, 136.46, 129.31, 124.14, 70.15, 70.08, 70.01, 69.89, 69.69, 66.89, 58.70, 50.36, 40.88, 40.48, 36.23, 34.57, 29.53, 24.46. HRMS $C_{26}H_{42}N_9O_7S^-$ (M − H)$^-$: calculated *m/z* = 624.2933, found *m/z* = 624.2933.

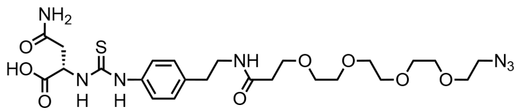


**S10**

*PTC-N:* Preparative TLC was developed with 25% MeOH in DCM. PTC-N (**S10**) was isolated as white solid (96.5 mg, 83%). $^{1}$H NMR (400 MHz, MeOH-d4) δ 7.34 (d, *J* = 7.9 Hz, 2H), 7.22 (d, *J* = 7.9 Hz, 2H), 3.74–3.57 (m, 16H), 3.38 (dt, *J* = 14.1, 5.0 Hz, 4H), 3.19 (q, *J* = 7.4 Hz, 1H), 2.98–2.85 (m, 2H), 2.78 (t, *J* = 7.1 Hz, 2H), 2.41 (t, *J* = 6.0 Hz, 2H). $^{13}$C NMR (101 MHz, MeOH-d4) δ 179.31, 174.70, 172.67, 172.58, 136.55, 129.36, 123.88, 70.13, 70.06, 70.01, 69.97, 69.87, 69.68, 66.91, 55.85, 50.35, 40.59, 40.46, 37.48, 36.25, 36.20, 34.54, 16.70. HRMS $C_{28}H_{46}N_9O_7S^-$ (M − H)$^-$: calculated *m/z* = 652.3246, found *m/z* = 652.3249.

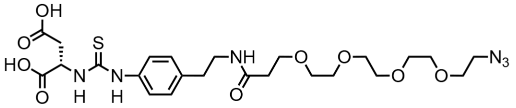


**S11**

*PTC-D:* Preparative TLC was developed with 25% MeOH in DCM. PTC-D (**S11**) was isolated as white foam (97.1 mg, 83%). $^{1}$H NMR (400 MHz, MeOH-d4) δ 7.34 (d, *J* = 6.9 Hz, 2H), 7.23 (d, *J* = 7.9 Hz, 2H), 3.80–3.51 (m, 16H), 3.39 (dt, *J* = 14.5, 5.5 Hz, 4H), 3.20 (q, *J* = 7.2 Hz, 1H), 3.06–2.82 (m, 2H), 2.78 (t, *J* = 7.1 Hz, 2H), 2.42 (t, *J* = 6.0 Hz, 2H). $^{13}$C NMR (101 MHz, MeOH-d4) δ 179.42, 175.17, 172.61, 136.52, 129.31, 123.86, 70.10, 70.03, 69.95, 69.85, 69.66, 66.91, 50.34, 40.57, 40.45, 36.17, 34.53, 29.32. HRMS $C_{24}H_{35}N_6O_9S^-$ (M − H)$^-$: calculated *m/z* = 583.2191, found *m/z* = 583.2192.

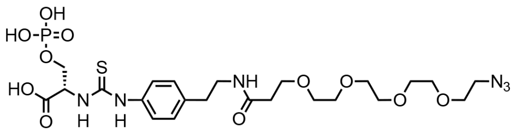

**S12**

*PTC-$^{p}$S:* Preparative TLC was developed with 50% MeOH in DCM. PTC-$^{p}$S (**S12**) was isolated as light yellow solid (48.4 mg, 38%). $^{1}$H NMR (400 MHz, MeOH-d4) δ 7.40 (d, *J* = 7.9 Hz, 2H), 7.22 (d, *J* = 7.9 Hz, 2H), 3.78–3.50 (m, 16H), 3.46–3.33 (m, 4H), 3.15 (s, 3H), 2.78 (t, *J* = 7.0 Hz, 2H), 2.42 (t, *J* = 5.8 Hz, 2H). $^{13}$C NMR (101 MHz, MeOH-d4) δ 172.63, 136.25, 129.14, 70.10, 70.03, 69.95, 69.85, 69.67, 66.90, 50.32, 40.47, 36.14, 34.51, 7.69. $^{31}$P NMR (162 MHz, DMSO-d6) δ -0.17. HRMS $C_{23}H_{36}N_6O_{11}PS^-$ $(M - H)^-$: calculated *m/z* = 635.1905, found *m/z* = 635.1903.

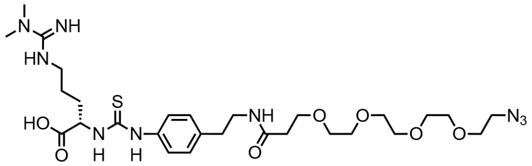

**S13**

*PTC-ADMA:* Preparative TLC was developed with 30% MeOH in DCM. PTC-ADMA (**S13**) was isolated as white solid (102.7 mg, 78%). $^{1}$H NMR (400 MHz, MeOH-d4) δ 7.30 (d, *J* = 8.2 Hz, 2H), 7.24 (d, *J* = 8.2 Hz, 2H), 3.51–3.36 (m, 4H), 3.33–3.23 (m, 3H), 3.02 (d, *J* = 0.8 Hz, 6H), 2.78 (t, *J* = 7.2 Hz, 2H), 2.42 (t, *J* = 6.0 Hz, 2H), 2.04 (dq, *J* = 14.1, 6.4 Hz, 1H), 1.95–1.79 (m, 1H), 1.78–1.53 (m, *J* = 6.9 Hz, 2H). $^{13}$C NMR (101 MHz, MeOH-d4) δ 1 178.94, 176.84, 172.60, 156.21, 136.72, 136.32, 129.43, 124.00, 70.05, 70.00, 69.98, 69.91, 69.81, 69.65, 66.92, 58.53, 50.32, 41.81, 40.46, 37.25, 36.11, 34.56, 29.59, 24.35. HRMS $C_{24}H_{35}N_6O_9S^-$ $(M - H)^-$: calculated *m/z* = 583.2191, found *m/z* = 583.2192.

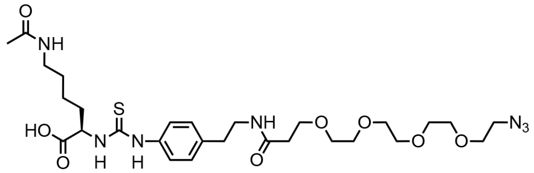

**S14**

*PTC-$^{Ac}$K:* Preparative TLC was developed with 30% MeOH in DCM. PTC- $^{Ac}$K (**S14**) was isolated as white solid (92.2 mg, 72%). $^{1}$H NMR (400 MHz, MeOH-d4) δ 7.31 (d, *J* = 8.4 Hz, 2H), 7.24 (d, *J* = 8.4 Hz, 2H), 3.92–3.52 (m, 16H), 3.41 (t, *J* = 7.1 Hz, 2H), 3.37 (t, *J* = 5.0 Hz, 2H), 3.21 (q, *J* = 7.4 Hz, 1H), 3.14 (t, *J* = 6.9 Hz, 2H), 2.79 (t, *J* = 7.1 Hz, 2H), 2.42 (t, *J* = 6.0 Hz, 2H), 2.14–1.96 (m, 1H), 1.91 (s, 3H), 1.81 (dq, *J* = 13.9, 7.0 Hz, 1H), 1.60–1.45 (m, 2H). $^{13}$C NMR (101 MHz, MeOH-d4) δ 179.30, 176.61, 172.60, 171.71, 136.69, 129.40, 124.02, 70.09, 70.02, 69.94, 69.84, 69.66, 66.90, 58.83, 50.32, 40.44, 39.06, 36.14, 34.53, 31.99, 28.76, 22.24, 21.18. HRMS $C_{24}H_{35}N_6O_9S^-$ $(M - H)^-$: calculated *m/z* = 638.2977, found *m/z* = 638.2977.

### Supplementary Text

Theoretical considerations on the factors that affect PEA efficiency

The efficiency of PEA is a function of the fraction of PTC amino acids that is bound by the antibody $fA_{\text{bound}}$.

The relationship between the concentration of free primer-tagged antibody $[A]$, free DNA-barcoded PTC amino acid $[P]$, and complex $[PA]$ at equilibrium can be written as Eq. 1, where $K_d$ is the dissociation constant of the interaction.

$$K_d = \frac{[A][P]}{[PA]} \quad [1]$$

Under our assay condition, primer-tagged antibody is in large excess comparing to the PTC amino acid, and thus the concentration of free primer-tagged antibody $[A]$ approximately equals to the initial concentration of primer-tagged antibody $[A]_0$. If we set initial concentration of DNA-barcoded PTC amino acid as $[P]_0$, Eq.1 can be written as Eq. 2.

$$K_d = \frac{[A]_0 \times ([P]_0 - [PA])}{[PA]} \quad [2]$$

The fraction of the PTC amino acid that is bound by antibody $fA_{\text{bound}}$ can be calculated using Eq. 3.

$$fA_{\text{bound}} = \frac{[PA]}{[P]_0} = \frac{[A]_0}{[A]_0 + K_d} \quad [3]$$

Therefore, PEA efficiency is a function of $[A]_0$ and $K_d$, and for a given primer-tagged antibody, PEA efficiency is determined by $[A]_0$ and not affected by the concentration of input $[P]_0$.

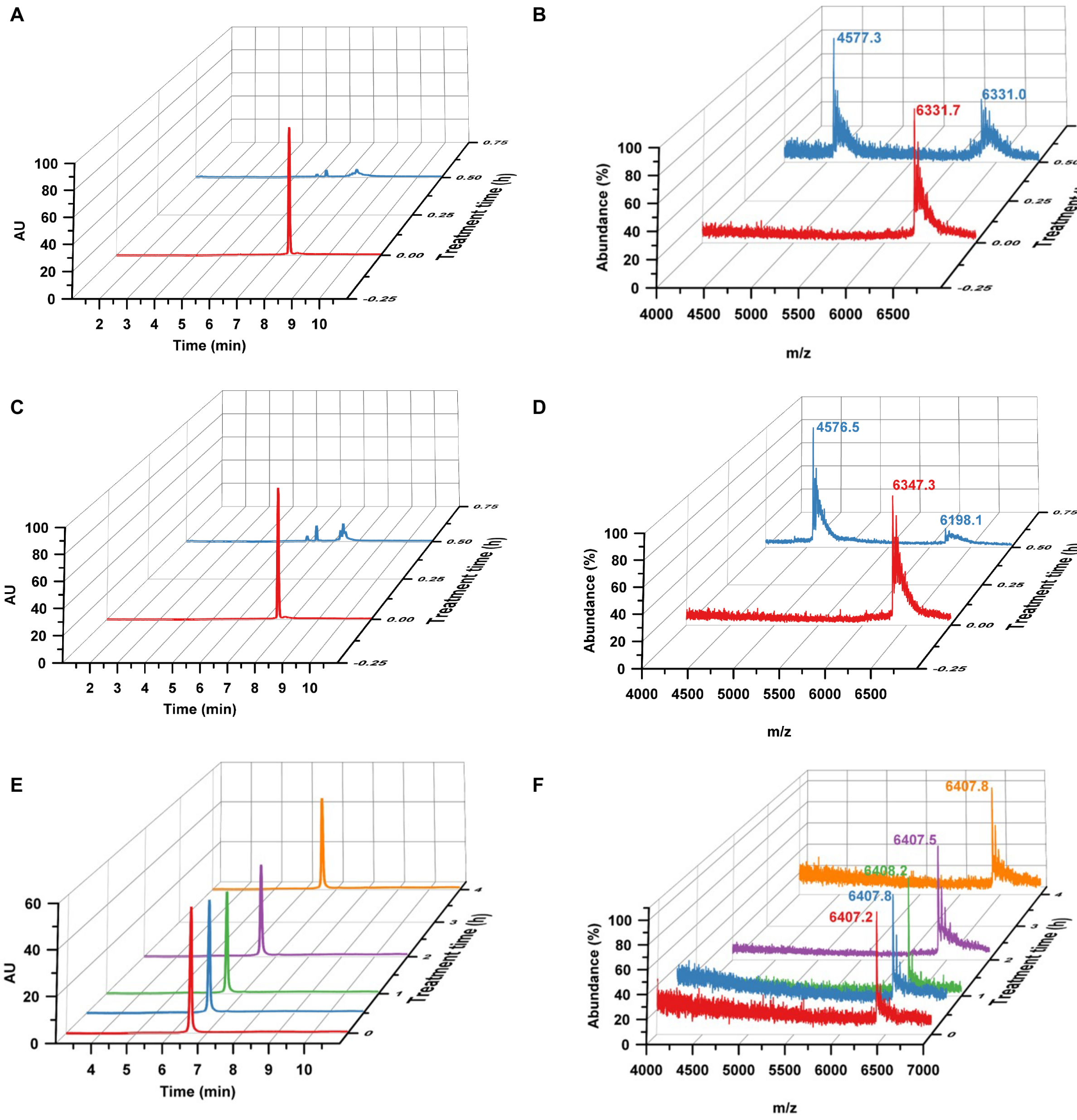


**Fig. S1. Stability of DNA during $BF_3 \cdot Et_2O$ treatment.** (**A**) HPLC chromatograms of **ODN-S1** before (red) and after (blue) treatment with 40 mM $BF_3 \cdot Et_2O$ for 30 min. (**B**) MS spectra of **ODN-S1** before (red) and after (blue) treatment with 40 mM $BF_3 \cdot Et_2O$ for 30 min. The new species with m/z = 4,577.3 is consistent with 5’-d$T_{15}$-$OPO_3^{2-}$ (calculated m/z = 4577.7). (**C**) HPLC chromatograms of **ODN-S2** before (red) and after (blue) treatment with 40 mM $BF_3 \cdot Et_2O$ for 30 min. (**D**) MS spectra of oligonucleotide **ODN-S2** before (red) and after (blue) treatment with 40 mM $BF_3 \cdot Et_2O$ for 30 min. The new species with m/z = 4,576.5 is consistent with 5’-d$T_{15}$-$OPO_3^{2-}$. (**E**) HPLC chromatograms of 7-deazapurine nucleotide-substituted oligonucleotide **ODN-2** before (red) and after up to 4h (orange) of treatment with 40 mM $BF_3 \cdot Et_2O$. (**F**) MS spectra of **ODN-2** before (red) and after up to 4h (orange) of treatment with 40 mM $BF_3 \cdot Et_2O$.

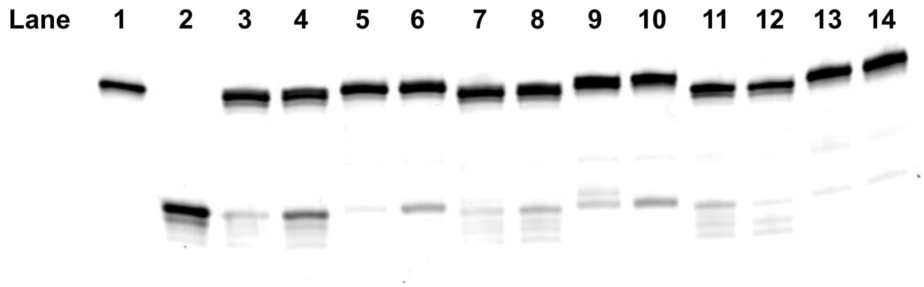


**Fig. S2. Primer extension with deazapurine nucleotide-substituted primers and templates.** Denaturing PAGE gel showing primer extension on deazapurine-substituted template (**ODN-S5)** with controls. Lane 1: chemically synthesized full-length extension product (**ODN-S8**). Lane 2: native DNA primer (**ODN-S7**) only. Lane 3: primer extension by Sequenase version 2.0 DNA polymerase with deazapurine-substituted primer (**ODN-S6**) with $c^7$dATP, dTTP, $c^7$dGTP, and dCTP (300 µM each) after 15 min at 37 °C. Lane 4: primer extension by Sequenase version 2.0 with native primer (**ODN-S7**) using standard dNTPs (300 µM each) after 15 min at 37 °C. Lane 5: primer extension by Sequenase version 2.0 with **ODN-S6** using a mixture of $c^7$dATP, dTTP, $c^7$dGTP, and dCTP (300 µM each) after 30 min at 37 °C. Lane 6: primer extension by Sequenase version 2.0 with **ODN-S7** using standard dNTPs (300 µM each) after 30 min at 37 °C. Lane 7: primer extension by Klenow fragment (exo-) with **ODN-S6** using a mixture of $c^7$dATP, dTTP, $c^7$dGTP, and dCTP (300 µM each) after 15 min at 37 °C. Lane 8: primer extension by Klenow fragment (exo-) with **ODN-S7** using standard dNTPs (300 µM each) after 15 min at 37 °C. Lane 9: primer extension by Klenow fragment (exo-) with **ODN-S6** using a mixture of $c^7$dATP, dTTP, $c^7$dGTP, and dCTP (300 µM each) after 30 min at 37 °C. Lane 10: primer extension by Klenow fragment (exo-) with **ODN-S7** using standard dNTPs (300 µM each) after 30 min at 37 °C. Lane 11: primer extension by *Bst* 3.0 DNA polymerase with **ODN-S6** using a mixture of $c^7$dATP, dTTP, $c^7$dGTP, and dCTP (300 µM each) after 15 min at 55 °C. Lane 12: primer extension by *Bst* 3.0 with **ODN-S7** using standard dNTPs (300 µM each) after 15 min at 55 °C. Lane 13: primer extension by *Bst* 3.0 with **ODN-S6** using a mixture of $c^7$dATP, dTTP, $c^7$dGTP, and dCTP (300 µM each) after 30 min at 55 °C. Lane 14: primer extension by *Bst* 3.0 with **ODN-S7** using standard dNTPs (300 µM each) after 30 min at 55 °C.

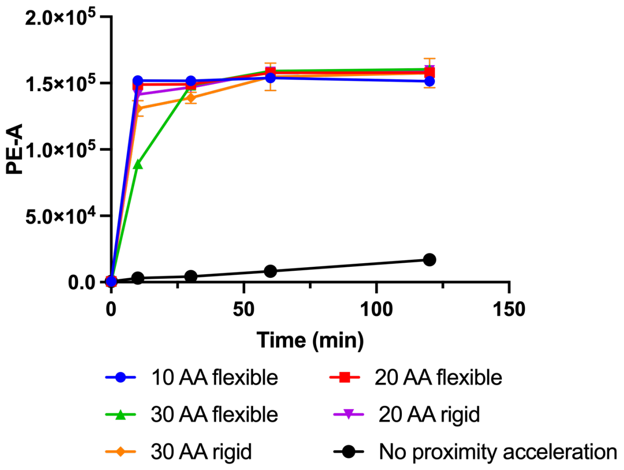


**Fig. S3. Time course of proximity SPAAC using peptides of varying length and rigidity.** Progress of proximity SPAAC was monitored by flow cytometry using Cy3-labeled complementary DNA (**ODN-S13**). The "no proximity acceleration" control was performed using the flexible 10 aa peptide and a non-complementary DNA (**ODN-S14**). Peptide sequences are shown in **Table S3**.

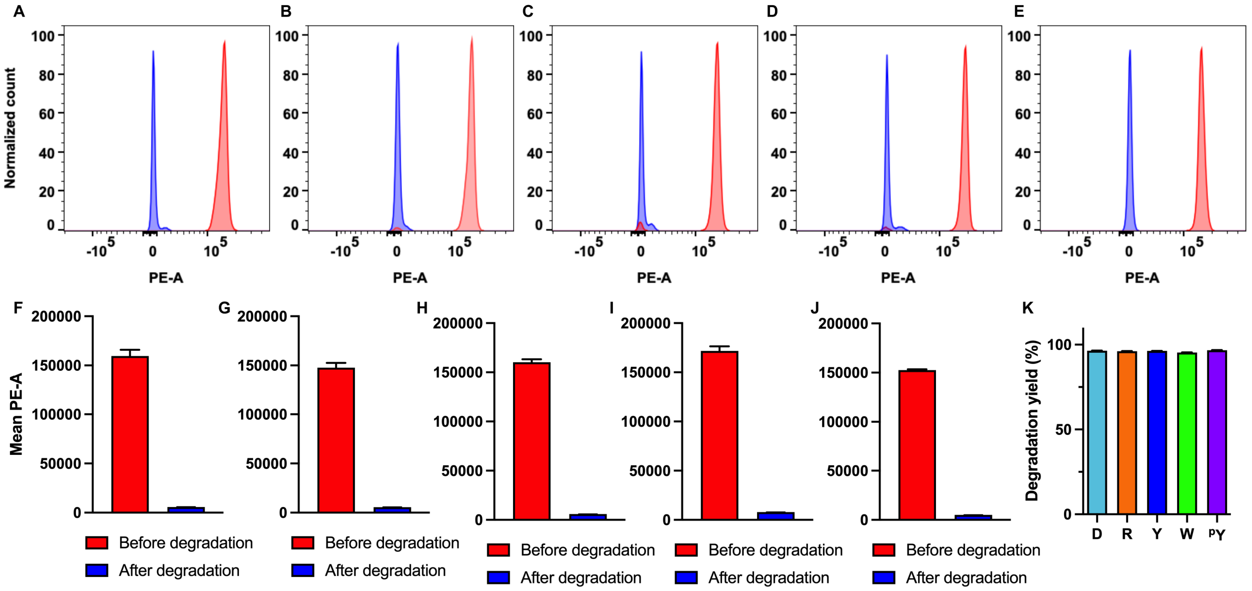


**Fig. S4. Compatibility of DNA-encoded Edman degradation with various N-terminal amino acids.** (**A** to **E**) Representative flow cytometry histograms for first-cycle Edman degradation of five peptides, (A) RGGGGGX, (B) DGGGGGX, (C) YGGGGGX, (D) WGGGGGX, and (E) $^{p}$YGGGGGX. (**F** to **J**) Fluorescence measured by flow cytometry before and after the degradation of the same five peptides, the ratio of which was used to calculate the degradation yield. (**K**) Degradation yield of first-cycle Edman degradation of the same five peptides, RGGGGGX (96.2 ± 0.2%), DGGGGGX (96.4 ± 0.3%), YGGGGGX (96.3 ± 0.1%), WGGGGGX (95.4 ± 0.2%), and $^{p}$YGGGGGX (96.8 ± 0.1%) as measured by flow cytometry. N = 3 for all data in (**F** to **K**).

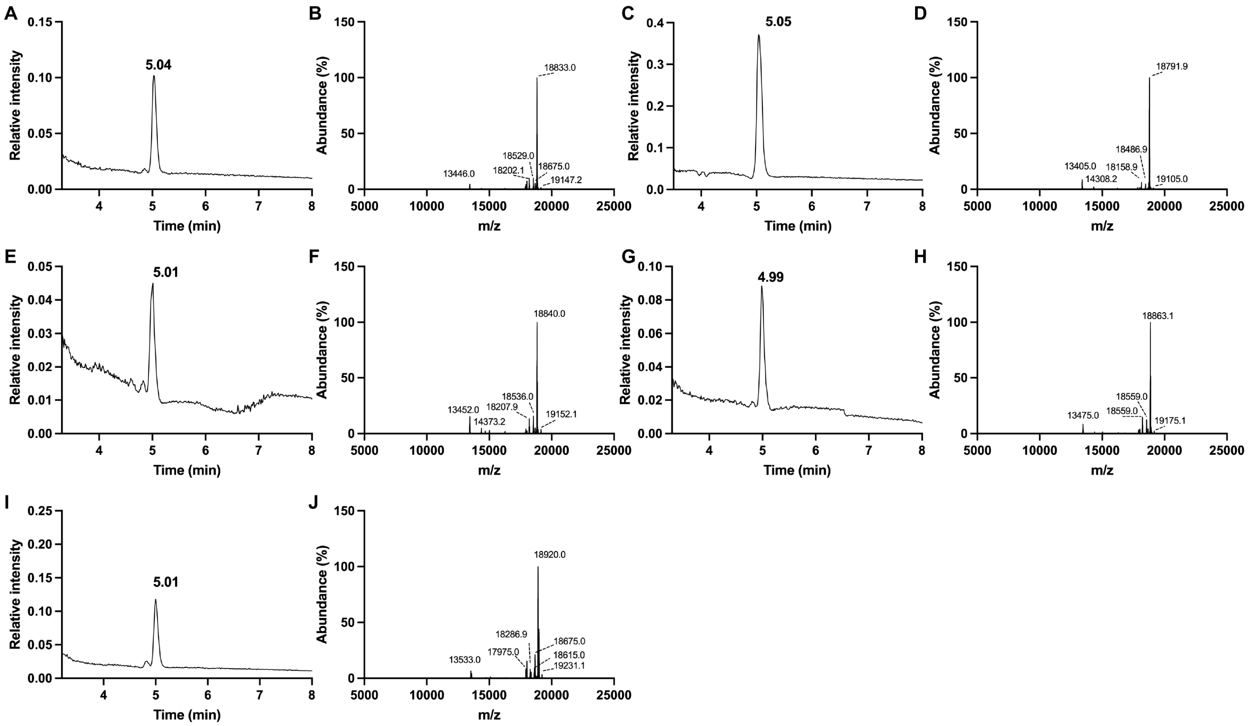


**Fig. S5. LC-MS analysis of PTC amino acids generated by various N-terminal amino acids.** LC-MS analysis of PTC amino acids generated by the first cycle of DNA-encoded Edman degradation on peptide (**A** and **B**) RGGGGGX, (**C** and **D**) DGGGGGX, (**E** and **F**) YGGGGGX, (**G** and **H**) WGGGGGX, and (**I** and **J**) $^{p}$YGGGGGX. (A, C, E, G, and I) show HPLC chromatographs of DNA-tagged PTC amino acids. (B, D, F, H, and J) show MS spectra of DNA barcoded PTC amino acids. (**B**) DNA-tagged PTC-R (calculated MW = 18,833.2, observed MW = 18,833.0), peaks at 134,446.0, 18,202.1, and 18,529.0 correspond to truncated products from incomplete primer extension. The peak at 19,147.2 corresponds to product from A-tailing. The peak at 18,675.0 corresponds to a product of oxidative basic degradation of PTC amino acids. (**D**) DNA-tagged PTC-D (calculated MW = 18,792.1, observed MW = 18,791.9), peaks with lower MW correspond to truncated products from incomplete primer extension. The peak at 19,105.0 corresponds to product from A-tailing. (**F**) DNA-tagged PTC-Y (calculated MW = 18,840.2, observed MW = 18,840.0), peaks with lower MW correspond to truncated products from incomplete primer extension. The peak at 19,152.1 corresponds to product from A-tailing. The peak at 18,675.0 corresponds to a product of oxidative basic degradation of PTC amino acids. (**H)** DNA-tagged PTC-W (calculated MW = 18,863.2, observed MW = 18,863.1), peaks with lower MW correspond to truncated products of incomplete primer extension. The peak at 19,175.1 corresponds to a product of A-tailing. The peak at 18,675.0 corresponds to a product of oxidative basic degradation of PTC amino acids. (**J**) DNA-tagged PTC-$^{p}$Y(calculated MW = 18,920.2, observed MW = 18,920.0), peaks with lower MW correspond to truncated product resulted from incomplete primer extension. The peak at 19,231.1 corresponds to a product of A-tailing. The peak at 18,675.0 corresponds to a product of oxidative basic degradation of PTC amino acids.

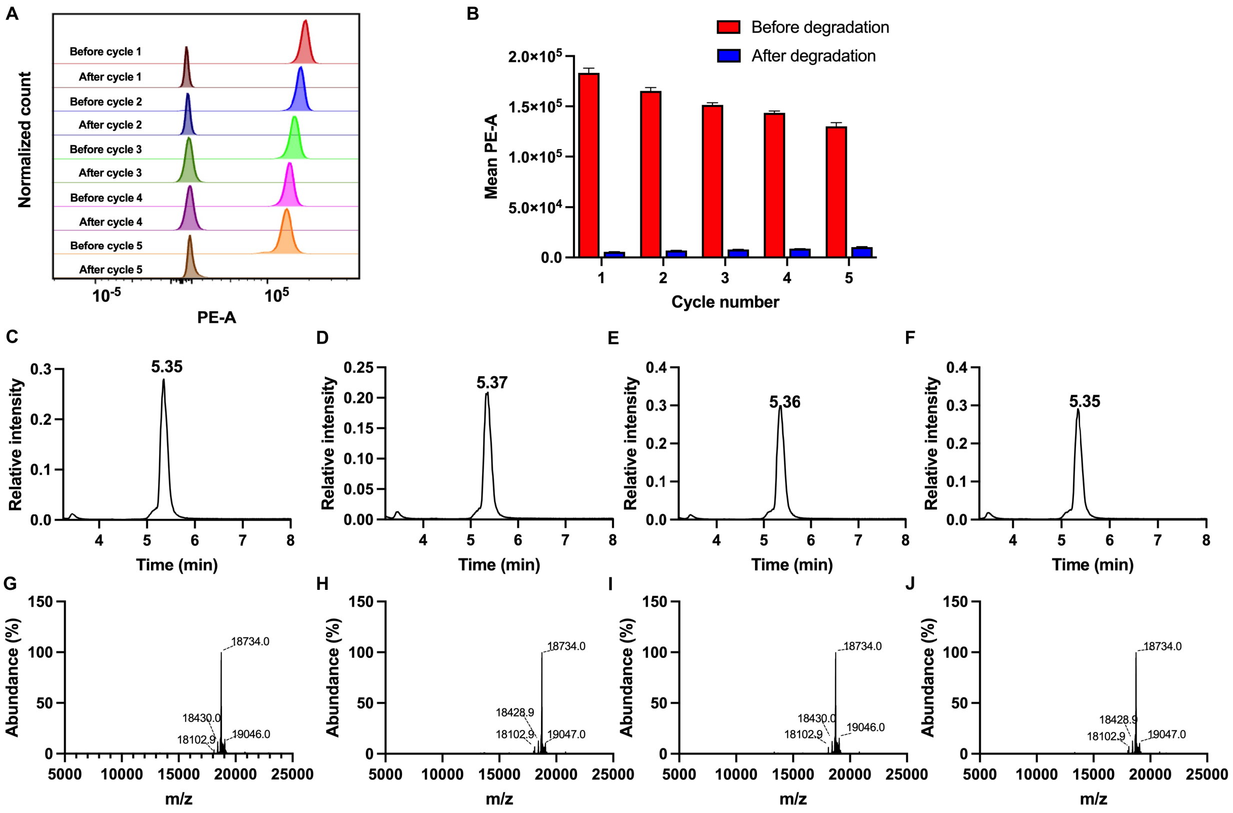


**Fig. S6. Five-cycle DNA-encoded Edman degradation on peptide FGGGGGX.** (**A**) Representative flow cytometry histograms of Edman degradation of peptide FGGGGGX. (**B**) Fluorescence measured by flow cytometry before and after the degradation (N = 3), the ration of which was used to calculate the degradation yield shown in **Fig. 3D**. (**C** to **F**) HPLC chromatograms of products generated by the second (C), third (D), fourth (E), and fifth (F) cycle of DNA-encoded Edman degradation on peptide FGGGGGX. (**G** to **J**) MS spectra of DNA-barcoded PTC-G generated by the second (G), third (H), fourth (I), and fifth (J) cycle of DNA-encoded Edman degradation (calculated MW = 18,734.1, observed MW = 18,734.0). For the second cycle, peaks at 18,102.9 and 18,428.9 correspond to truncated products of incomplete primer extension. The peak at 19,046.0 corresponds to the product of A-tailing. Similar product compositions were observed from the third to the fifth cycle.

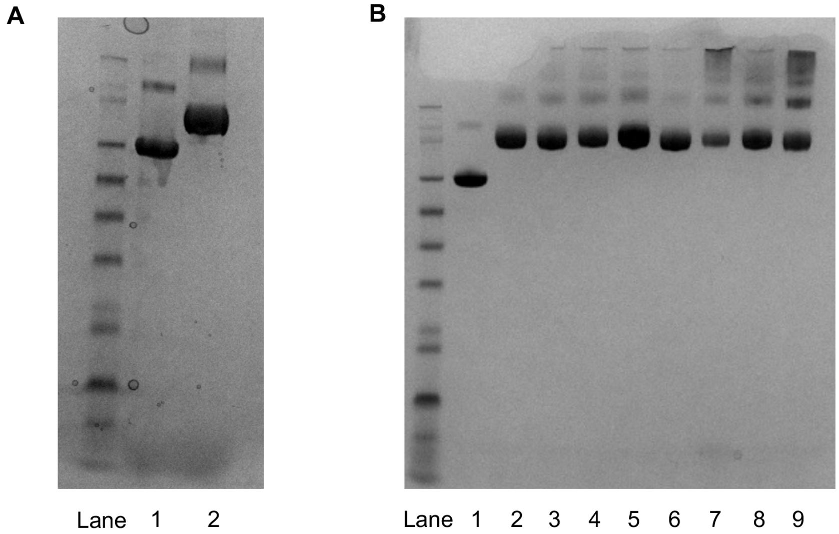


**Fig. S7. Preparation of PTC amino acid-conjugated BSA.** (**A**) Analysis of conjugation of TFP ester-PEG4-DBCO with BSA by reducing SDS-PAGE. Lane 1: BSA before conjugation. Lane 2: BSA after conjugation. (**B**) Analysis of conjugation of azide-modified PTC amino acids with DBCO-modified BSA by reducing SDS-PAGE. Lane 1: BSA alone. Lane 2: BSA conjugated with PTC-Q. Lane 3: BSA conjugated with PTC-D. Lane 4: BSA conjugated with PTC-F. Lane 5: BSA conjugated with PTC-R. Lane 6: BSA conjugated with PTC-N. Lane 7: BSA conjugated with PTC-E. Lane 8: BSA conjugated with PTC-W. Lane 9: BSA conjugated with PTC-Y.

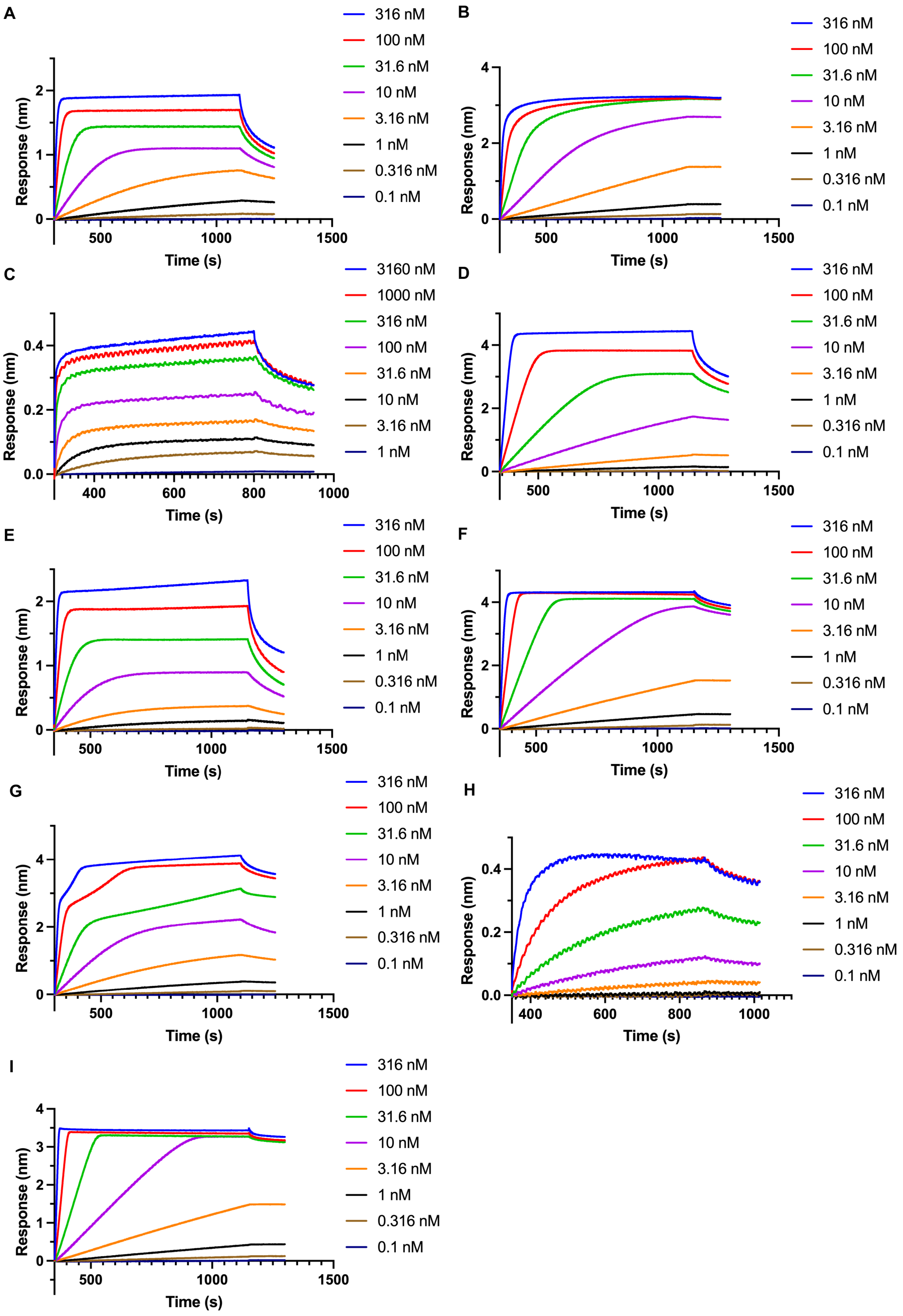


**Fig. S8. Representative biolayer interferometry (BLI) sensorgrams.** (**A**) Anti-PTC-F antibody against immobilized PTC-F. (**B**) Anti-PTC-W antibody against immobilized PTC-W. (**C**) Anti-PTC-Y antibody against immobilized PTC-Y. (**D**) Anti-PTC-D antibody against immobilized PTC-D. (**E**) Anti-PTC-R antibody against immobilized PTC-R. (**F**) Phosphotyrosine antibody PY20 against immobilized PTC-$^{p}$Y. (**G**) Acetylated lysine antibody RM101 against PTC-$^{Ac}$K. (**H**) Asymmetric dimethyl arginine antibody 21C7 against PTC-ADMA. (**I**) Phosphoserine antibody 3C171 against PTC-$^{p}$S.

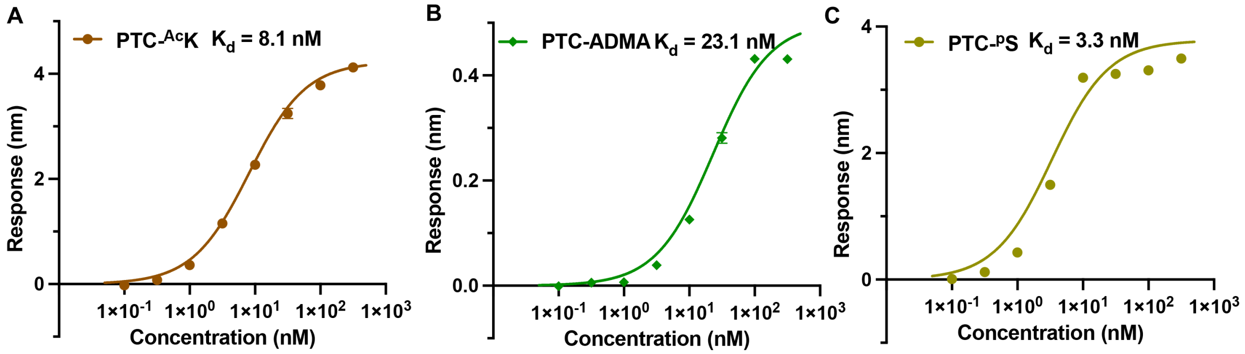


**Fig. S9. Antibody binding to PTC amino acids bearing PTM.** Binding curves of (**A**) acetylated lysine antibody RM101 against PTC-$^{Ac}$K, (**B**) asymmetric dimethyl arginine antibody 21C7 against PTC-ADMA, and (**C**) phosphoserine antibody 3C171 against PTC-$^{p}$S.

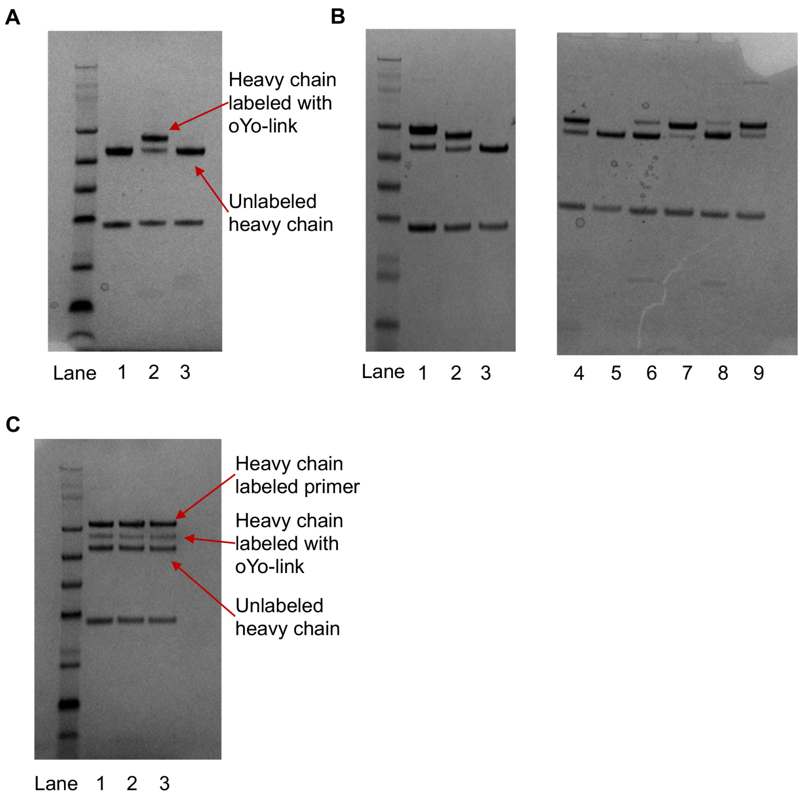


**Fig. S10. Labeling of antibody with primers for PEA.** (**A**) Conjugation of oYo-link tetrazine with anti-PTC-F antibody analyzed by reducing SDS-PAGE. Lane 1: unconjugated antibody. Lane 2: anti-PTC-F antibody conjugated with mouse IgG1 (mIgG1) oYo-link tetrazine kit. Lane 3: anti-PTC-F antibody conjugated with oYo-link tetrazine kit for non-mIgG1 antibodies. Anti-PTC-F antibody was exclusively conjugated by the mIgG1 oYo-link tetrazine kit. (**B**) Screening of specificity of oYo-link kits against anti-PTC amino acid antibodies. Lane 1: anti-PTC-F antibody conjugated with mIgG1 oYo-link tetrazine kit. Lane 2: anti-PTC-D antibody conjugated with mIgG1 oYo-link tetrazine kit. Lane 3: anti-PTC-D antibody conjugated with oYo-link tetrazine kit. Lane 4: anti-PTC-Y antibody conjugated with mIgG1 oYo-link tetrazine kit. Lane 5: anti-PTC-Y antibody conjugated with oYo-link tetrazine kit. Lane 6: anti-PTC-R antibody conjugated with mIgG1 oYo-link tetrazine kit. Lane 7: anti-PTC-R antibody conjugated with oYo-link tetrazine kit. Lane 8: anti-PTC-W antibody conjugated with mIgG1 oYo-link tetrazine kit. Lane 9: anti-PTC-W antibody conjugated with oYo-link tetrazine kit. Anti-PTC-D and -Y antibodies were specifically conjugated by the mIgG1 oYo-link tetrazine kit, while anti-PTC-R and -W antibodies were specifically conjugated with the oYo-link tetrazine kit. Note that the isotype of PY20 is reported as IgG2b, and as such, it is labeled with the oYo-link tetrazine kit. (**C**) SDS-PAGE analysis of conjugation of tetrazine-modified anti-PTC-F antibody with the following TCO-labeled primers: Lane 1, 5 nt primer **ODN-7**; Lane 2, 6 nt primer **ODN-8**; Lane 3: 7 nt primer **ODN-9**. About 50% of heavy chain was primer-labeled after oYo-link conjugation and the TCO-tetrazine click reaction.

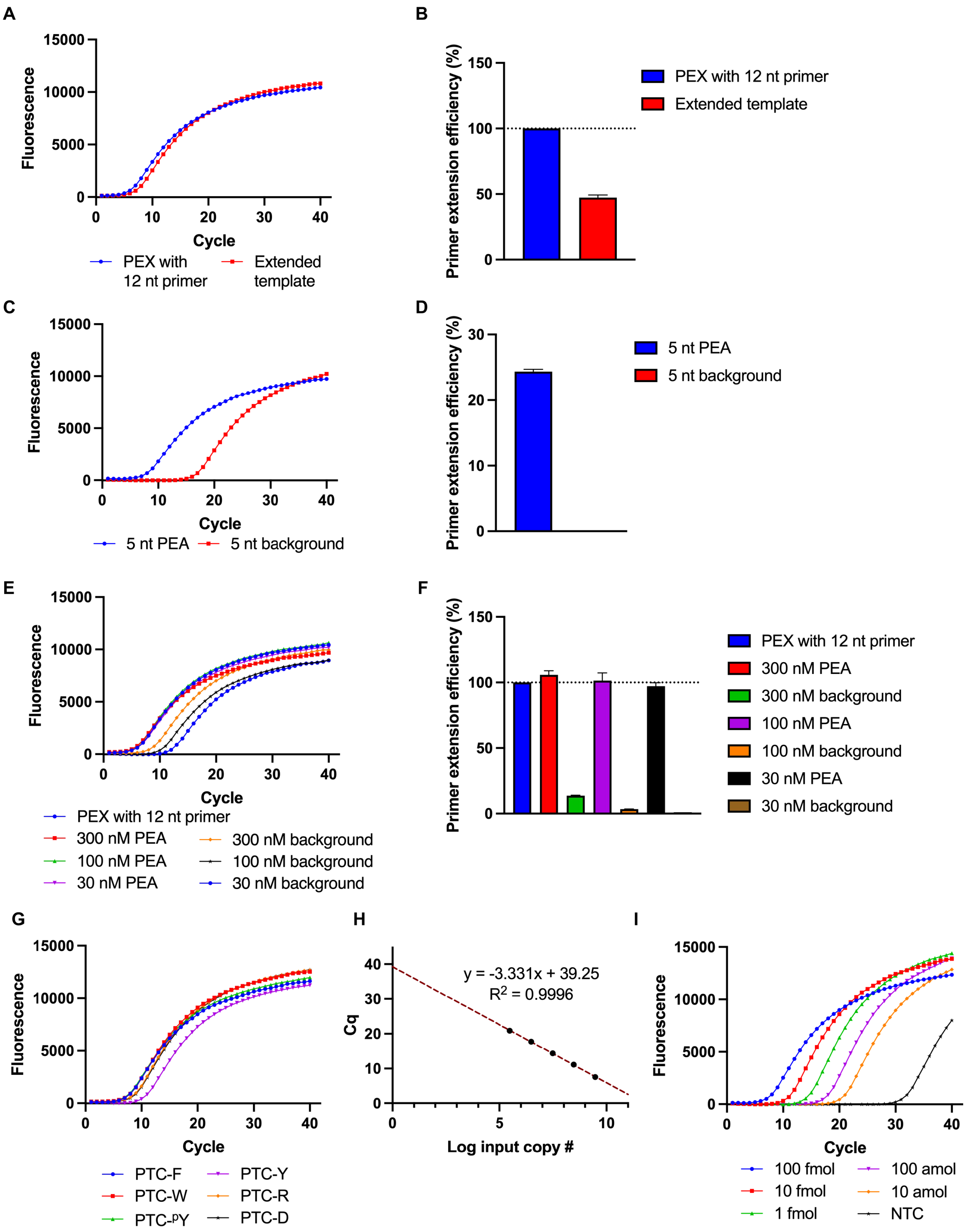


**Fig. S11. Quantification of PEA efficiency via qPCR.** (**A**) Representative qPCR curve of positive controls. PEA was carried out with a 12 nt primer (**ODN-10**) and the SA bead-immobilized template **ODN-6**. qPCR reactions were carried out with both beads after PEA (blue) and beads modified with chemically synthesized single-stranded DNA with sequence of the expected extension product (**ODN-S16**; red). Because PEA yields a duplex product, these two reactions should be separated by ΔCq = 1 if primer extension is complete. We observed a ΔCq =

1.08 ± 0.6, which indicates complete primer extension, and we therefore used **ODN-10** to normalize PEA efficiency under other conditions. (**B**) PEA efficiency for conditions tested in (A) (N = 3). (**C**) Representative qPCR curve quantifying PEA efficiency of anti-PTC-F antibody modified with 5 nt primer **ODN-7** at 100 nM. (**D**) PEA efficiency for conditions tested in (C) (N = 3). (**E**) Representative qPCR curve quantifying PEA efficiency of anti-PTC-F antibody modified with 6 nt primer **ODN-8** at concentrations ranging from 30 to 300 nM. (**F**) PEA efficiency for the conditions tested in (E) (N = 3). (**G**) Representative qPCR curve quantifying PEA efficiency of different antibodies modified with **ODN-8** at 100 nM. (**H**) qPCR standard curve in which the Cq values were plotted against the logarithm of the copy number of the template (N = 3). (**I**) Representative qPCR curve quantifying PEA efficiency of anti-PTC-F antibody modified with **ODN-8** at template input ranging from 100 fmol to 10 amol in 20 µL reactions (concentrations ranging from 500 fM to 5 nM).

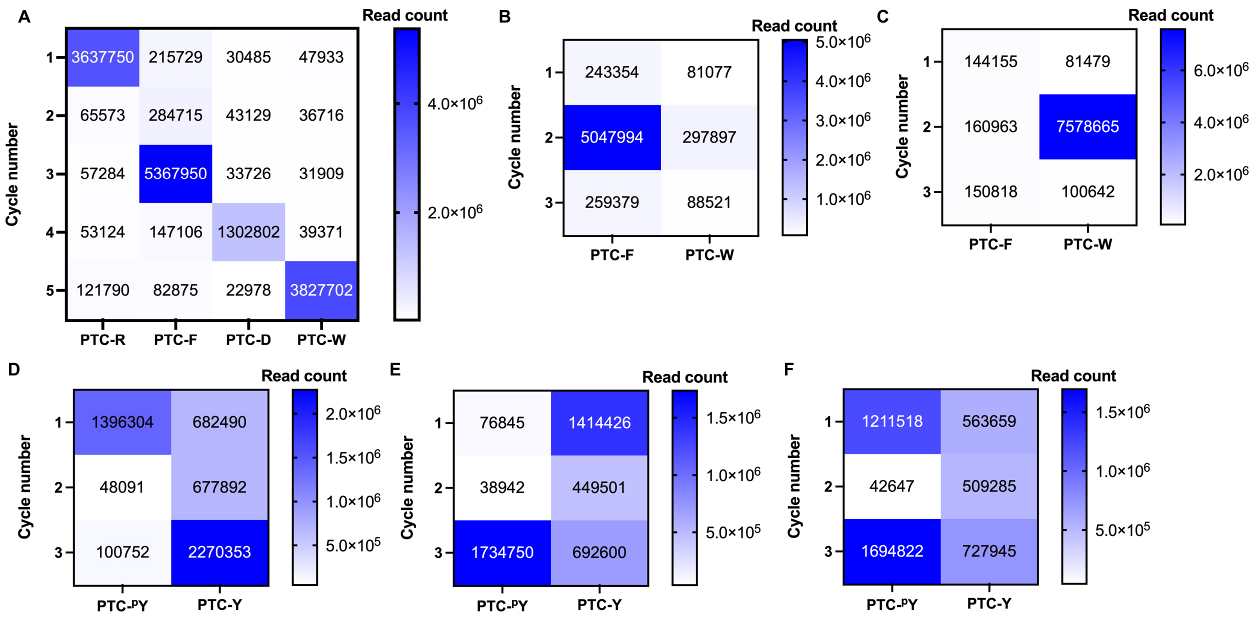


**Fig. S12. Raw DNA sequencing read counts during peptide sequencing. (A)** Heatmap of raw barcode read counts from the first five residues of RGFDWGX. (**B** to **C**) Heatmap of raw barcode read counts for peptide sequences AFG (B) and AWG (C). (**D** to **F**) Heatmap of raw barcode read counts for peptide sequences $^{p}$YGY (D), YG$^{p}$Y (E), and $^{p}$YG$^{p}$Y (F).

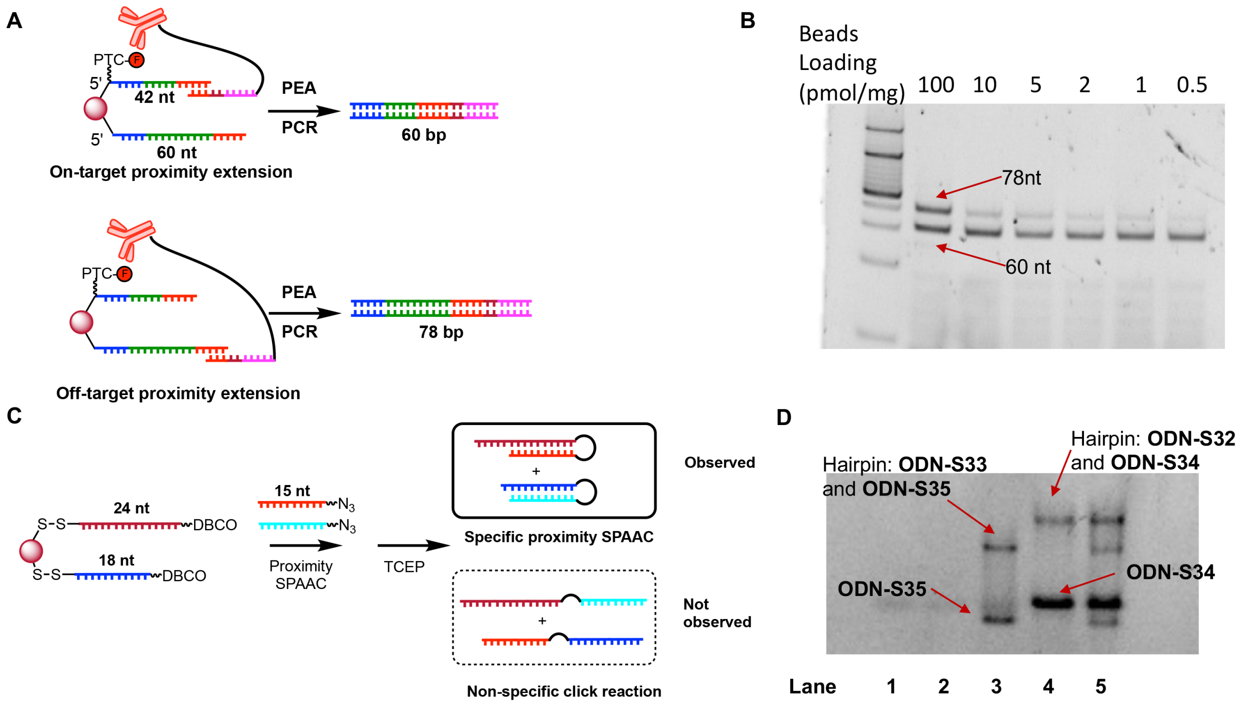


**Fig. S13. Probing barcode crosstalk during PEA and proximity SPAAC.** (**A**) Scheme of experiment to measure barcode crosstalk during PEA. A 1:1 mixture of 42 nt biotinylated DNA barcoded PTC-F (**ODN-6-PTC-F**) and a biotinylated 60 nt barcoding DNA without PTC-F (**ODN-S31**) was immobilized on SA beads. After PEA, on-target proximity extension generates a 60 bp DNA duplex whereas off-target proximity extension yields a 78 bp DNA duplex. (**B**) Native PAGE analysis of on- and off-target proximity extension at varying loading densities. Off-target proximity extension is greatly reduced by reducing the template loading density. (**C**) Scheme of experiment to measure barcode crosstalk during proximity SPAAC. A 1:1 mixture of DBCO-modified 24 nt DNA (**ODN-S32**) and 18 nt DNA (**ODN-S33**) was immobilized on carboxylic acid beads. The beads were reacted with a mixture of azide-modified 15 nt complementary strands **ODN-S34** and **ODN-S35**. **ODN-S35** contains one less spacer18 modification, producing a mobility difference. After the click reaction, the DNA on the beads is released with TCEP and analyzed by native PAGE. Specific proximity SPAAC gives rise to two different hairpins, whereas a non-specific click reaction generates two linear products. (**D**) Barcode crosstalk during proximity SPAAC based on analysis of TCEP-released DNA by native PAGE. Lane 1: DNA released from **ODN-S33-**modified beads. Lane 2: DNA released from **ODN-S32-**modified beads. No DNA is visible in these lanes because the DNA was not fluorescently labeled. Lane 3: DNA released by **ODN-S33-**modified beads reacted with azide-modified **ODN-S35**. Lane 4: DNA released from **ODN-S32-**modified beads reacted with azide-modified **ODN-S34**. Lane 5: DNA released from magnetic beads modified with **ODN-S32** and **ODN-S33** and reacted with a mixture of azide-modified **ODN-S34** and **ODN-S35**. Only hairpins formed by specific proximity SPAAC were observed.

### Table S1. Oligonucleotides used in the study

| | Sequence | Note | MS |
|---|---|---|---|
| **ODN-1** | 5'-d(ATC TGA CTG GCA ATG ATC GCT) | Native DNA stability test | |
| **ODN-2** | 5'-d(c$^7$ATC Tc$^7$Gc$^7$A CTc$^7$G c$^7$GCc$^7$A c$^7$ATc$^7$G c$^7$ATC c$^7$GCT) | Deazapurine-substituted DNA stability test | **Fig. 2** |
| **ODN-3** | 5'-**3222**-d(c$^7$ACT CCC CTc$^7$A TCT CTc$^7$A c$^7$ATC c$^7$ACc$^7$A T) | Model peptide barcode anchor DNA | **p. 37** |
| **ODN-4** | 5'-$^{2-}O_3PO$-d(TCC c$^7$ATT Cc$^7$AT CTc$^7$A TTC c$^7$Ac$^7$AT CTC TCc$^7$A)-**2224** | Model peptide barcode DNA | **p. 37** |
| **ODN-5** | 5'-**42522**-d(TTT TTT c$^7$Gc$^7$Ac$^7$G c$^7$Ac$^7$Gc$^7$A TTc$^7$G c$^7$Ac$^7$AT c$^7$Ac$^7$Gc$^7$A T) | DBCO-modified biotinylated primer | **p. 38** |
| **ODN-6** | 5'-**42522**-d(c$^7$Gc$^7$GC c$^7$Ac$^7$AC c$^7$Gc$^7$GC c$^7$ATT TTC c$^7$Gc$^7$Ac$^7$G CCc$^7$A c$^7$GTc$^7$A c$^7$ATc$^7$A c$^7$Ac$^7$Gc$^7$A c$^7$Gc$^7$AC Tc$^7$GC CTc$^7$A CCc$^7$G) | PEA template | **p. 38** |
| **ODN-6-PTC-F** | 5'-**S3**-**42522**-d(c$^7$Gc$^7$GC c$^7$Ac$^7$AC c$^7$Gc$^7$GC c$^7$ATT TTC c$^7$Gc$^7$Ac$^7$G CCc$^7$A c$^7$GTc$^7$A c$^7$ATc$^7$A c$^7$Ac$^7$Gc$^7$A c$^7$Gc$^7$AC Tc$^7$GC CTc$^7$A CCc$^7$G) | PEA templated-conjugated PTC-F | **p. 39** |
| **ODN-6-PTC-W** | 5'-**S4**-**42522**-d(c$^7$Gc$^7$GC c$^7$Ac$^7$AC c$^7$Gc$^7$GC c$^7$ATT TTC c$^7$Gc$^7$Ac$^7$G CCc$^7$A c$^7$GTc$^7$A c$^7$ATc$^7$A c$^7$Ac$^7$Gc$^7$A c$^7$Gc$^7$AC Tc$^7$GC CTc$^7$A CCc$^7$G) | PEA templated-conjugated PTC-W | **p. 39** |
| **ODN-6-PTC-Y** | 5'-**S5**-**42522**-d(c$^7$Gc$^7$GC c$^7$Ac$^7$AC c$^7$Gc$^7$GC c$^7$ATT TTC c$^7$Gc$^7$Ac$^7$G CCc$^7$A c$^7$GTc$^7$A c$^7$ATc$^7$A c$^7$Ac$^7$Gc$^7$A c$^7$Gc$^7$AC Tc$^7$GC CTc$^7$A CCc$^7$G) | PEA templated-conjugated PTC-Y | **p. 40** |
| **ODN-6-PTC-D** | 5'-**S11**-**42522**-d(c$^7$Gc$^7$GC c$^7$Ac$^7$AC c$^7$Gc$^7$GC c$^7$ATT TTC c$^7$Gc$^7$Ac$^7$G CCc$^7$A c$^7$GTc$^7$A c$^7$ATc$^7$A c$^7$Ac$^7$Gc$^7$A c$^7$Gc$^7$AC Tc$^7$GC CTc$^7$A CCc$^7$G) | PEA templated-conjugated PTC-D | **p. 40** |
| **ODN-6-PTC-R** | 5'-**S9**-**42522**-d(c$^7$Gc$^7$GC c$^7$Ac$^7$AC c$^7$Gc$^7$GC c$^7$ATT TTC c$^7$Gc$^7$Ac$^7$G CCc$^7$A c$^7$GTc$^7$A c$^7$ATc$^7$A c$^7$Ac$^7$Gc$^7$A c$^7$Gc$^7$AC Tc$^7$GC CTc$^7$A CCc$^7$G) | PEA templated-conjugated PTC-R | **p. 41** |
| **ODN-6-PTC-$^p$Y** | 5'-**S6**-**42522**-d(c$^7$Gc$^7$GC c$^7$Ac$^7$AC c$^7$Gc$^7$GC c$^7$ATT TTC c$^7$Gc$^7$Ac$^7$G CCc$^7$A c$^7$GTc$^7$A c$^7$ATc$^7$A c$^7$Ac$^7$Gc$^7$A c$^7$Gc$^7$AC Tc$^7$GC CTc$^7$A CCc$^7$G) | PEA templated-conjugated PTC-$^p$Y | **p. 41** |
| **ODN-7** | 5'-**6222**-d(TAC TGT ACC CTC TGT GCG CGG TA) | 5 nt PEA primer | **p. 42** |
| **ODN-8** | 5'-**6222**-d(TAC TGT ACC CTC TGT GCG CGG TAG) | 6 nt PEA primer | **p. 42** |
| **ODN-9** | 5'-**6222**-d(TAC TGT ACC CTC TGT GCG CGG TAG G) | 7 nt PEA primer | **p. 43** |
| **ODN-10** | 5'-d(TAC TGT ACC CTC TGT GCG CGG TAG GCA GTC) | 12 nt primer for PEA positive control | |

### Table S1 (cont'd)

| | Sequence | Note | MS |
|---|---|---|---|
| **ODN-11** | 5'-**3222**-d(Cc$^7$Gc$^7$G Tc$^7$Ac$^7$G c$^7$GCc$^7$A c$^7$GTC TC) | Peptide barcode anchor DNA | **p. 43** |
| **ODN-12** | 5'-$^{2-}$O$_3$PO-d(TTc$^7$A TTc$^7$A CNN NNN NNN NNT c$^7$Gc$^7$GC TCc$^7$G c$^7$Ac$^7$Ac$^7$A c$^7$ATc$^7$G CCc$^7$G TTc$^7$G CC)-**2224** | Peptide barcode DNA 1 | |
| **ODN-13** | 5'-**6222**-d(TAC TGT ACC CTC TGT GCG AGA CGG TAG) | Antibody tag 1 | **p. 44** |
| **ODN-14** | 5'-d(TCG TCG GCA GCG TCA GAT GTG TAT AAG AGA CAG NNN AAG GCA ACG GCA TTT TCG AG) | Adaptor FP cycle 1 | |
| **ODN-S1** | 5'-d(TTT TTT TTT TTT TTT ATT TT) | Native DNA stability test (single dA) | **Fig. S1** |
| **ODN-S2** | 5'-d(TTT TTT TTT TTT TTT GTT TT) | Native DNA stability test (single dG) | **Fig. S1** |
| **ODN-S3** | 5'-**3222**-d(c$^7$GCc$^7$G c$^7$ATC c$^7$ATT c$^7$GCC c$^7$Ac$^7$GT Cc$^7$Ac$^7$G c$^7$AT) | DNA stability test anchor | **p. 44** |
| **ODN-S4** | 5'-d(c$^7$Ac$^7$Ac$^7$A c$^7$Ac$^7$Ac$^7$A c$^7$Ac$^7$Ac$^7$A c$^7$Ac$^7$Ac$^7$A c$^7$Ac$^7$A)-**2223** | DNA stability test anchor poly-dA | **p. 45** |
| **ODN-S5** | 5'-d(c$^7$GTC c$^7$ATc$^7$G CTc$^7$A c$^7$Gc$^7$GT CTC c$^7$GCC c$^7$Ac$^7$Gc$^7$G c$^7$ACC TCc$^7$A TCC c$^7$Ac$^7$GC c$^7$AT) | Polymerase screen template | **p. 45** |
| **ODN-S6** | 5'-Cy3-d(c$^7$ATc$^7$G CTc$^7$G c$^7$Gc$^7$AT c$^7$Gc$^7$Ac$^7$G c$^7$GTC CTc$^7$G c$^7$GC) | Polymerase screen deazapurine primer | **p. 46** |
| **ODN-S7** | 5'-Cy3-d(ATG CTG GAT GAG GTC CTG GC) | Polymerase screen native primer | **p. 46** |
| **ODN-S8** | 5'-Cy3-d(c$^7$ATc$^7$G CTc$^7$G c$^7$Gc$^7$AT c$^7$Gc$^7$Ac$^7$G c$^7$GTC CTc$^7$G c$^7$GCc$^7$G c$^7$Ac$^7$Gc$^7$A CCT c$^7$Ac$^7$GC c$^7$ATc$^7$G c$^7$AC) | Polymerase screen expected product | **p. 47** |
| **ODN-S9** | 5'-d(ATG AAT GGA ATG TGA TTA GAG ATA G) | Splint | |
| **ODN-S10** | 5'-**3(S15)2**-Cy3-**2**-d(c$^7$ACT CCC CTc$^7$A TCT CTc$^7$A c$^7$ATC c$^7$ACc$^7$A T) | Disulfide-modified anchor DNA | **p. 47** |
| **ODN-S11** | 5'-**4222**-d(TTT TTT TTT TTT TTT) | Non-hybridizing DBCO-modified poly-dT | **p. 48** |
| **ODN-S12** | 5'-Cy3-d(CAT CTA TTC AAT CTC TCA AAA AA) | Cy3-labeled primer complementary strand | **p. 48** |
| **ODN-S13** | 5'-**4222**-d(Tc$^7$Gc$^7$A c$^7$Gc$^7$Ac$^7$G c$^7$ATT c$^7$Gc$^7$Ac$^7$A Tc$^7$Ac$^7$G c$^7$AT)-Cy3 | Cy3-labeled primer | **p. 49** |
| **ODN-S14** | 5'-**4222**-d(TTT TTT TTT TTT TTT)-Cy3 | Cy3-labeled non-hybridizing DBCO-modified poly-dT | **p. 49** |

**Table S1 (cont'd)**

| | Sequence | Note | MS |
|---|---|---|---|
| **ODN-S15** | 5'-(**S16**)**2**-d(TTT TTT TTT TTT TTT)-**224** | PTC amino acid attachment DNA for BLI | **p. 50** |
| **ODN-S15-PTC-F** | 5'-(**S16**)**2**-d(TTT TTT TTT TTT TTT)-**224-S3** | Biotinylated DNA-conjugated PTC-F | **p. 50** |
| **ODN-S15-PTC-Y** | 5'-(**S16**)**2**-d(TTT TTT TTT TTT TTT)-**224-S5** | Biotinylated DNA-conjugated PTC-Y | **p. 51** |
| **ODN-S15-PTC-W** | 5'-(**S16**)**2**-d(TTT TTT TTT TTT TTT)-**224-S4** | Biotinylated DNA-conjugated PTC-W | **p. 51** |
| **ODN-S15-PTC-$^{p}$Y** | 5'-(**S16**)**2**-d(TTT TTT TTT TTT TTT)-**224-S6** | Biotinylated DNA-conjugated PTC-$^{p}$Y | **p. 52** |
| **ODN-S15-PTC-D** | 5'-(**S16**)**2**-d(TTT TTT TTT TTT TTT)-**224-S11** | Biotinylated DNA-conjugated PTC-D | **p. 52** |
| **ODN-S15-PTC-E** | 5'-(**S16**)**2**-d(TTT TTT TTT TTT TTT)-**224-S7** | Biotinylated DNA-conjugated PTC-E | **p. 53** |
| **ODN-S15-PTC-N** | 5'-(**S16**)**2**-d(TTT TTT TTT TTT TTT)-**224-S10** | Biotinylated DNA-conjugated PTC-N | **p. 53** |
| **ODN-S15-PTC-R** | 5'-(**S16**)**2**-d(TTT TTT TTT TTT TTT)-**224-S9** | Biotinylated DNA-conjugated PTC-R | **p. 54** |
| **ODN-S15-PTC-Q** | 5'-(**S16**)**2**-d(TTT TTT TTT TTT TTT)-**224-S8** | Biotinylated DNA-conjugated PTC-Q | **p. 54** |
| **ODN-S15-PTC-ADMA** | 5'-(**S16**)**2**-d(TTT TTT TTT TTT TTT)-**224-S13** | Biotinylated DNA-conjugated PTC-ADMA | **p. 55** |
| **ODN-S15-PTC-$^{p}$S** | 5'-(**S16**)**2**-d(TTT TTT TTT TTT TTT)-**224-S12** | Biotinylated DNA-conjugated PTC-$^{p}$S | **p. 55** |
| **ODN-S15-PTC-$^{Ac}$K** | 5'-(**S16**)**2**-d(TTT TTT TTT TTT TTT)-**224-S14** | Biotinylated DNA-conjugated PTC-$^{Ac}$K | **p. 56** |
| **ODN-S16** | 5'-**32522**-d($c^7$G$c^7$GC $c^7$A$c^7$AC $c^7$G$c^7$GC $c^7$ATT TTC $c^7$G$c^7$A$c^7$G CC$c^7$A $c^7$GT$c^7$A $c^7$AT$c^7$A $c^7$A$c^7$G$c^7$A $c^7$G$c^7$AC T$c^7$GC CT$c^7$A CC$c^7$G C$c^7$GC $c^7$AC$c^7$A $c^7$G$c^7$A$c^7$G $c^7$G$c^7$GT $c^7$AC$c^7$A $c^7$GT$c^7$A) | Expected product of PEA | **p. 56** |
| **ODN-S17** | 5'-d(GGC AAC GGC ATT TTC GAG) | FP | |
| **ODN-S18** | 5'-d(TAC TGT ACC CTC TGT GCG) | RP | |
| **ODN-S19** | 5'-$^{2-}O_3$PO-d(TT$c^7$A TT$c^7$A CNN NNN NNN NNT C$c^7$GC TC$c^7$G $c^7$A$c^7$A$c^7$A $c^7$AT$c^7$G CC$c^7$G TT$c^7$G CC)-**2224** | Peptide barcode DNA 2 | |

**Table S1 (cont'd)**

| | **Sequence** | **Note** | **MS** |
|---|---|---|---|
| **ODN-S20** | 5'-$^{2-}O_3PO$-d(TTc$^7$A TTc$^7$A CNN NNN NNN NNT Cc$^7$AC TCc$^7$G c$^7$Ac$^7$Ac$^7$A c$^7$ATc$^7$G CCc$^7$G TTc$^7$G CC)-**2224** | Peptide barcode DNA 3 | |
| **ODN-S21** | 5'-d(GTA ATA AGA GAC TGC CTA CCG) | Sequencing splint | |
| **ODN-S22** | 5'-**42522**-d(c$^7$Gc$^7$GC c$^7$Ac$^7$AC c$^7$Gc$^7$GC c$^7$ATT TTC) | DBCO-modified biotinylated sequencing primer | **p. 57** |
| **ODN-S23** | 5'-**6222**-d(TAC TGT ACC CTC TGT GCG TAT CGG TAG) | Antibody tag 2 | **p. 57** |
| **ODN-S24** | 5'-**6222**-d(TAC TGT ACC CTC TGT GCG ACA CGG TAG) | Antibody tag 3 | **p. 58** |
| **ODN-S25** | 5'-**6222**-d(TAC TGT ACC CTC TGT GCG ATA CGG TAG) | Antibody tag 4 | **p. 58** |
| **ODN-S26** | 5'-d(TCG TCG GCA GCG TCA GAT GTG TAT AAG AGA CAG NNN ATG GCAA CGG CAT TTT CGA G) | Adaptor FP cycle 2 | |
| **ODN-S27** | 5'-d(TCG TCG GCA GCG TCA GAT GTG TAT AAG AGA CAG NNN ACG GCAA CGG CAT TTT CGA G) | Adaptor FP cycle 3 | |
| **ODN-S28** | 5'-d(TCG TCG GCA GCG TCA GAT GTG TAT AAG AGA CAG NNN GCG GCAA CGG CAT TTT CGA G) | Adaptor FP cycle 4 | |
| **ODN-S29** | 5'-d(TCG TCG GCA GCG TCA GAT GTG TAT AAG AGA CAG NNN GAG GCAA CGG CAT TTT CGA G) | Adaptor FP cycle 5 | |
| **ODN-S30** | 5'-d(GTC TCG TGG GCT CGG AGA TGT GTA TAA GAG ACA GNN NNT ACT GTA CCC TCT GTG CG) | Adaptor RP | |
| **ODN-S31** | 5'-**32522**-d(c$^7$Gc$^7$GC c$^7$Ac$^7$AC c$^7$Gc$^7$GC c$^7$ATT TTC c$^7$Gc$^7$Ac$^7$G CCT Tc$^7$Gc$^7$G Tc$^7$Ac$^7$A c$^7$ACC Tc$^7$AT c$^7$Gc$^7$AT TCc$^7$A c$^7$GTc$^7$A c$^7$ATc$^7$A c$^7$Ac$^7$Gc$^7$A c$^7$Gc$^7$AC Tc$^7$GC CTc$^7$A CCc$^7$G) | PEA template 60 nt | **p. 59** |
| **ODN-S32** | 5'-**(S17)222**-d(CTc$^7$G c$^7$Gc$^7$AC Tc$^7$GT c$^7$Gc$^7$Ac$^7$G c$^7$Ac$^7$AC TTT TTT TTT)-**(S15)22(S18)** | Hairpin anchor 1 | **p. 59** |
| **ODN-S33** | 5'-**(S17)222**-d(c$^7$Ac$^7$AT Cc$^7$GT c$^7$GTC c$^7$GTT c$^7$ACc$^7$A TTT)- **(S15)22(S18)** | Hairpin anchor 2 | **p. 60** |
| **ODN-S34** | 5'- Cy3-d(c$^7$GTT CTC c$^7$ACc$^7$A c$^7$GTC Cc$^7$Ac$^7$G)-**222(S19)** | Azide-modified hairpin forming strand 1 | **p. 60** |
| **ODN-S35** | 5'-Cy3-d(TTc$^7$A c$^7$GCc$^7$A Cc$^7$Ac$^7$G Cc$^7$Ac$^7$A Tc$^7$GT)-**22(S19)** | Azide-modified hairpin forming strand 2 | **p. 61** |

### Table S2. Oligonucleotide modifications

| | Structure | Note |
|---|---|---|
| **2** | 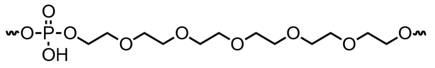 | Hexaethylene glycol (i.e., spacer18) modification |
| **3** | 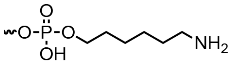 | Hexylamino modification |
| **4** | 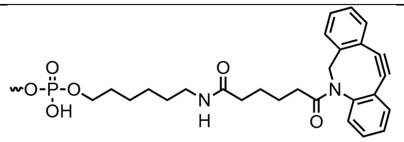 | DBCO modification |
| **5** | 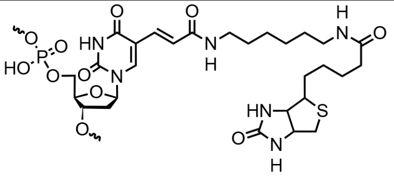 | Biotin-dT modification |
| **6** | 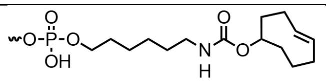 | TCO modification |
| **S15** | 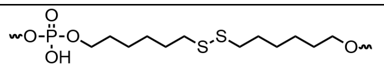 | Internal disulfide modification |
| **S16** | 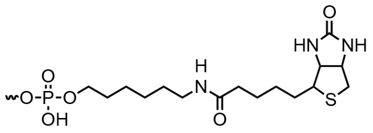 | 5′-Biotin modification |
| **S17** | 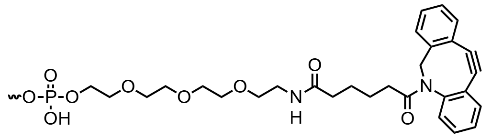 | 5′-DBCO-triethylene glycol (TEG) modification |
| **S18** | 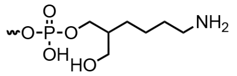 | 3′-C7 amino modification (compatible with 5′-DBCO-TEG modification) |
| **S19** | 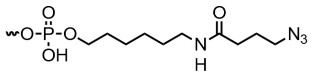 | Azide modification |

### Table S3. Peptides used in the study

| Sequence | Note |
|---|---|
| FGGGGGX | X = azidolysine here and elsewhere in table. |
| YGGGGGX | |
| WGGGGGX | |
| DGGGGGX | |
| RGGGGGX | |
| pYGGGGGX | |
| RGFDWGX | |
| AWGAGX | |
| AFGAGX | |
| pYGYGGX | |
| YGpYGGX | |
| pYGpYGGX | |
| GGGGSGGGGSX | Flexible 10 aa peptide |
| GGGGSGGGGSGGGGSGGGGSX | Flexible 20 aa peptide |
| GGGGSGGGGSGGGGSGGGGSGGGGSGGGGSX | Flexible 30 aa peptide |
| GSGSEPEPEPEPEPEPGSGSX | Rigid 20 aa peptide |
| GSGSEPEPEPEPEPEPGSGSEPEPEPEPSGX | Rigid 30 aa peptide |

### Table S4. Peptide and antibody barcodes

| Peptide or antibody | Barcoding DNA (barcode) |
|---|---|
| **Fig. 5B** | |
| RGFDWGX | Peptide barcode DNA 1 (CC) |
| Anti-PTC-R antibody | Antibody tag 1 (TCT) |
| Anti-PTC-F antibody | Antibody tag 2 (ATA) |
| Anti-PTC-D antibody | Antibody tag 3 (TGT) |
| Anti-PTC-W antibody | Antibody tag 4 (TAT) |
| **Fig. 5**, **D** and **E** | |
| AFGAGX | Peptide barcode DNA 1 (CC) |
| AWGAGX | Peptide barcode DNA 2 (CG) |
| Anti-PTC-F antibody | Antibody tag 1 (TCT) |
| Anti-PTC-W antibody | Antibody tag 2 (ATA) |
| **Fig. 5**, **F** to **H** | |
| pYGYGGX | Peptide barcode DNA 1 (CC) |
| YGpYGGX | Peptide barcode DNA 2 (CG) |
| pYGpYGGX | Peptide barcode DNA 3 (TG) |
| PY20 antibody | Antibody tag 4 (TAT) |
| Anti-PTC-Y antibody | Antibody tag 3 (TGT) |

**MS Spectra of modified oligonucleotides**

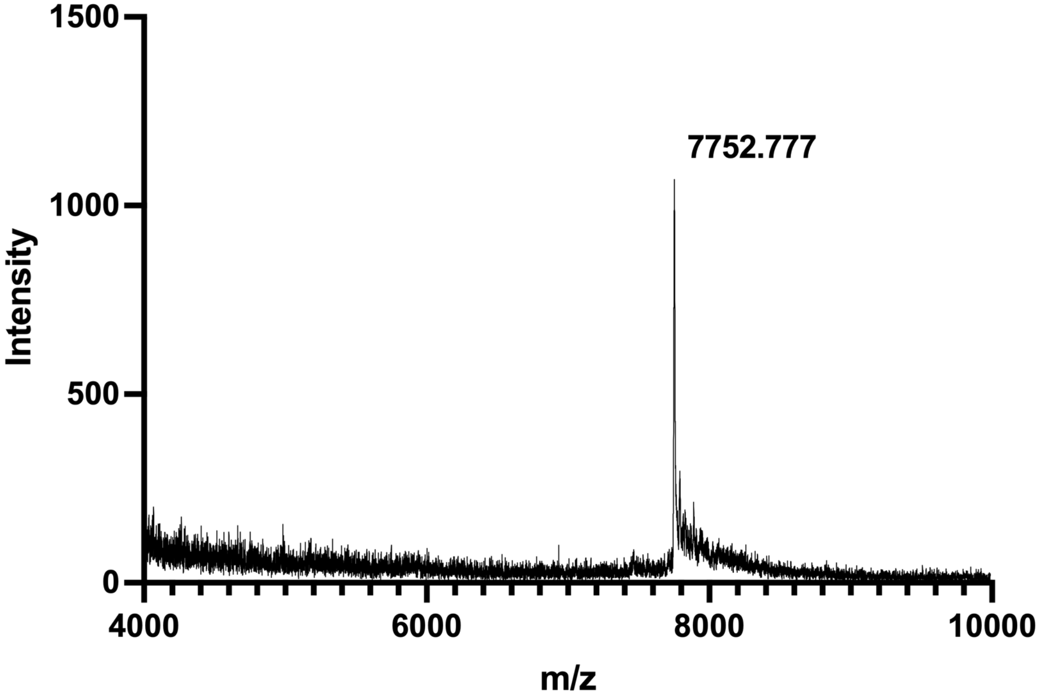


MALDI-TOF-MS spectrum of **ODN-3**. Calculated monoisotopic m/z = 7751.2.

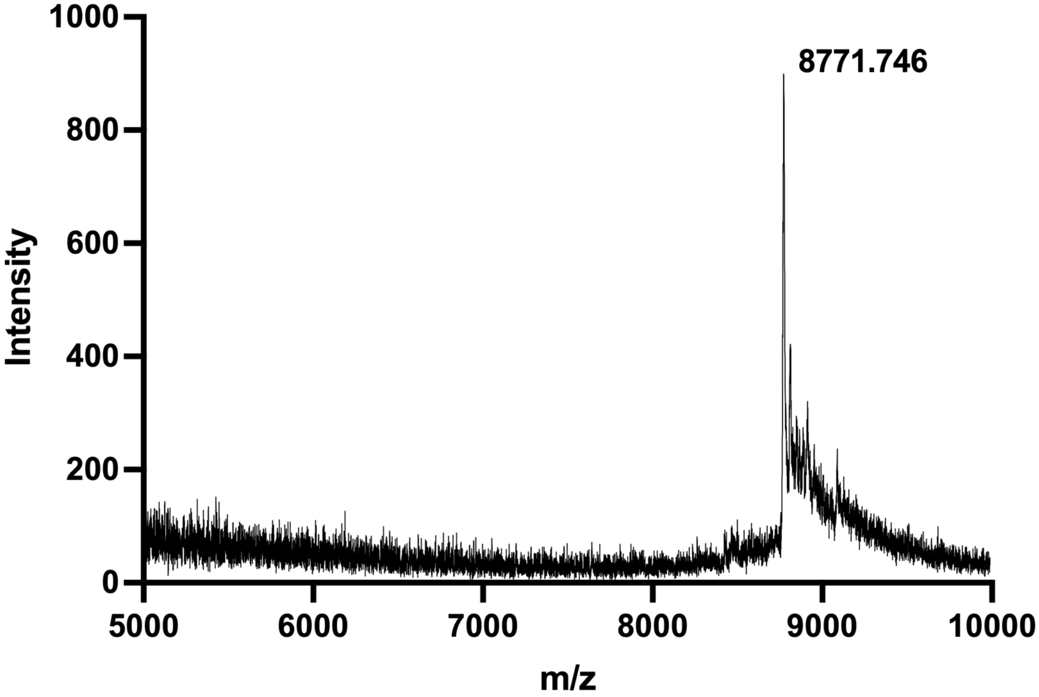


MALDI-TOF-MS spectrum of **ODN-4**. Calculated monoisotopic m/z = 8770.6.

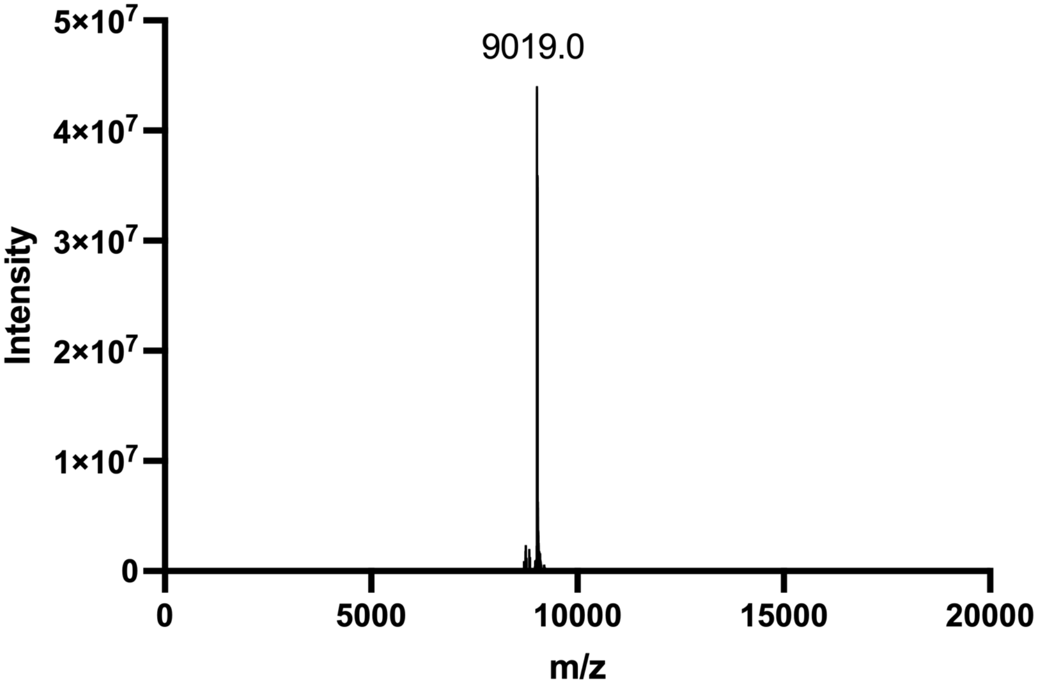


ESI-MS spectrum of **ODN-5**. Calculated MW = 9018.6.

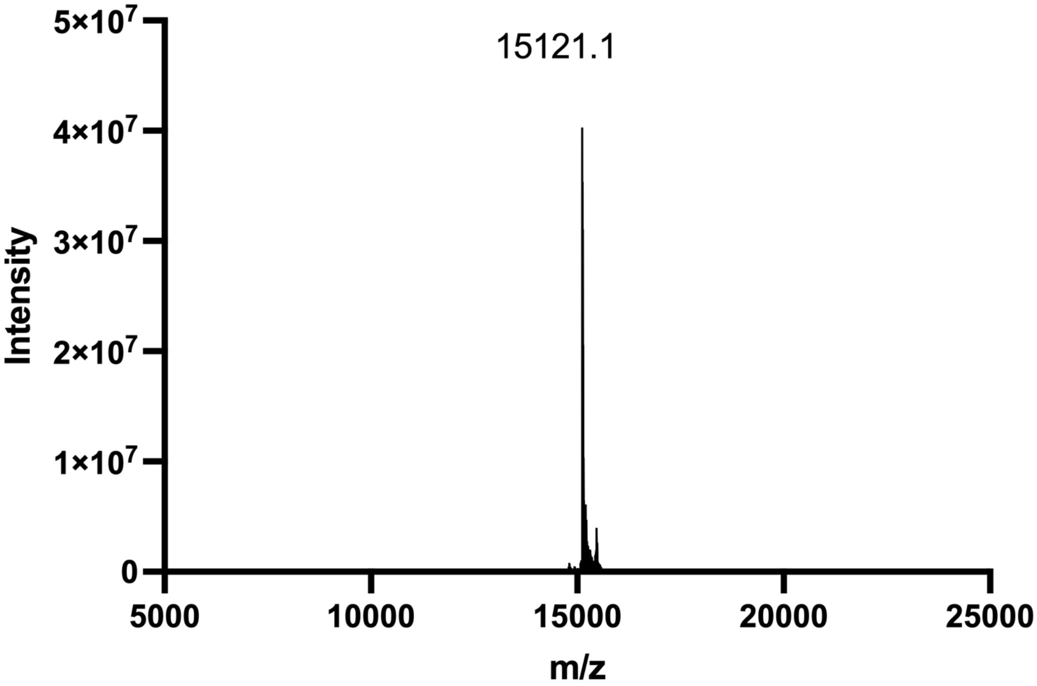


ESI-MS spectrum of **ODN-6**. Calculated MW = 15121.6.

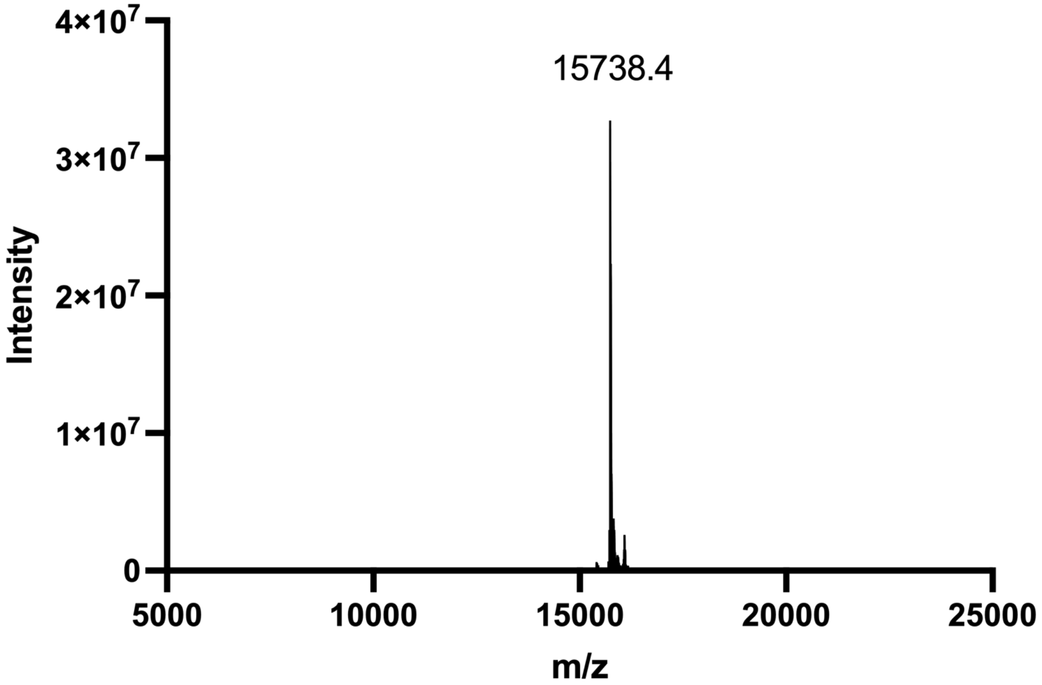


ESI-MS spectrum of **ODN-6-PTC-F**. Calculated MW = 15737.8.

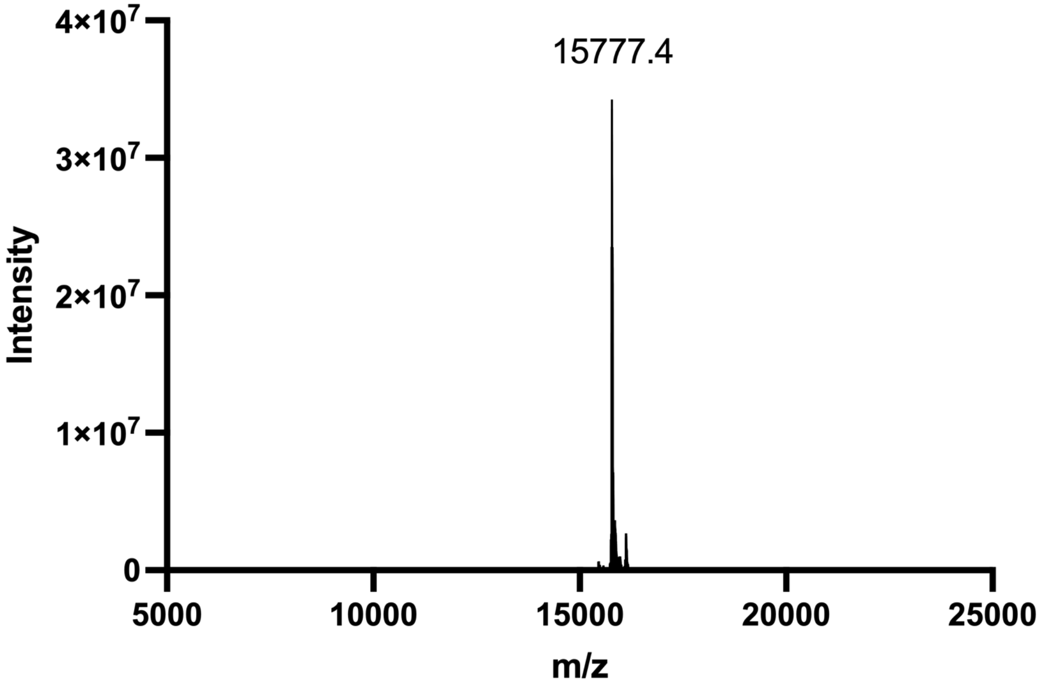


ESI-MS spectrum of **ODN-6-PTC-W**. Calculated MW = 15776.8.

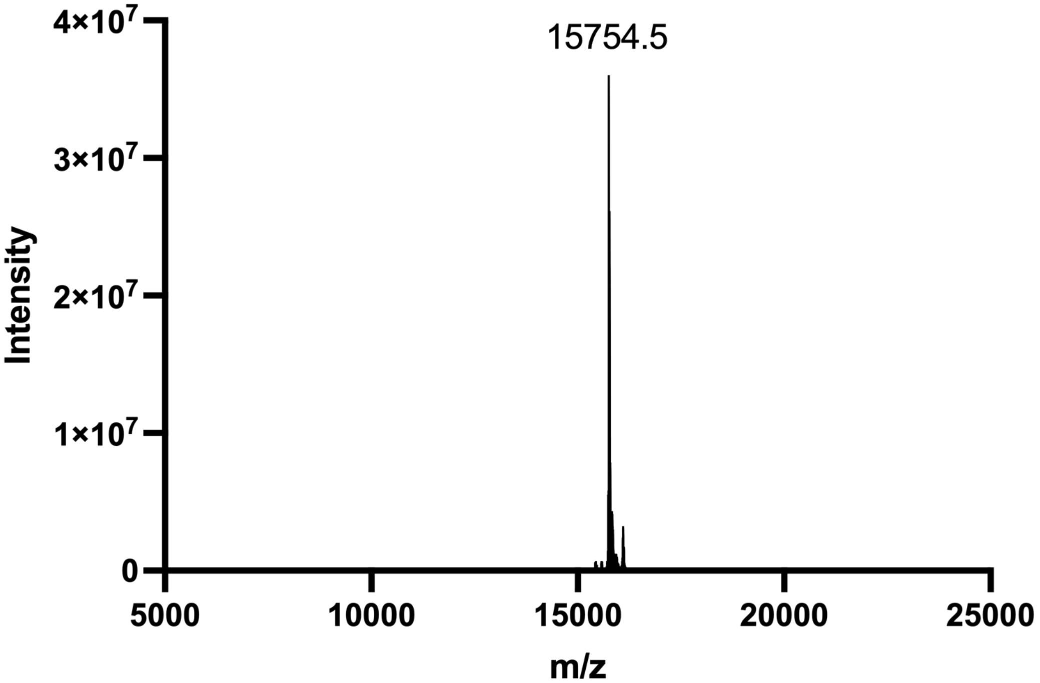


ESI-MS spectrum of **ODN-6-PTC-Y**. Calculated MW = 15753.8.

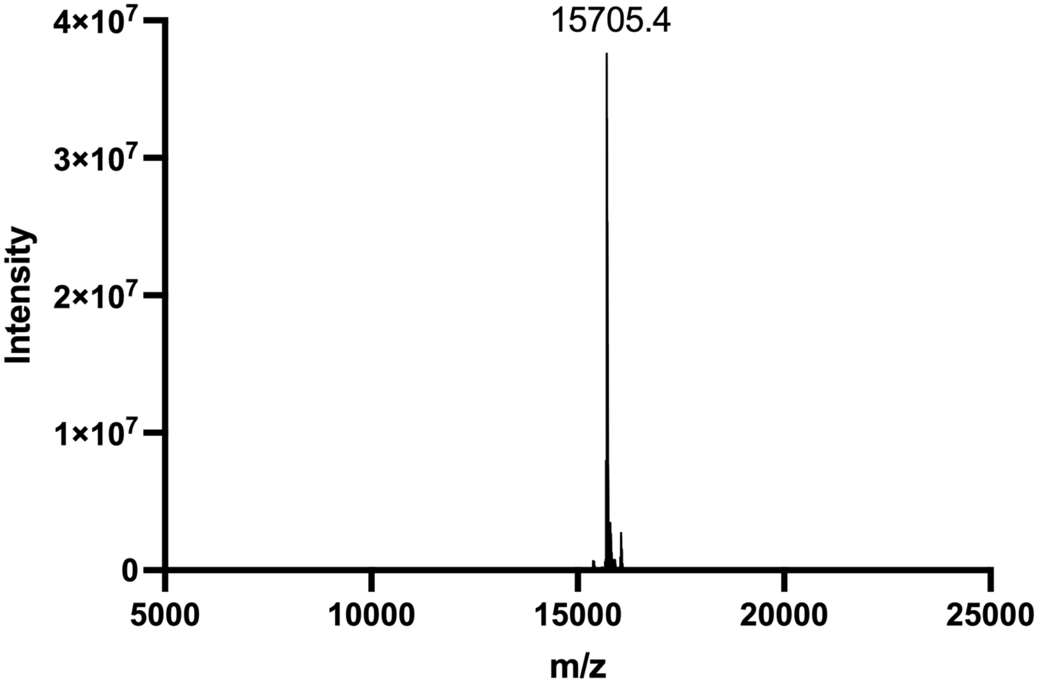


ESI-MS spectrum of **ODN-6-PTC-D**. Calculated MW = 15705.8.

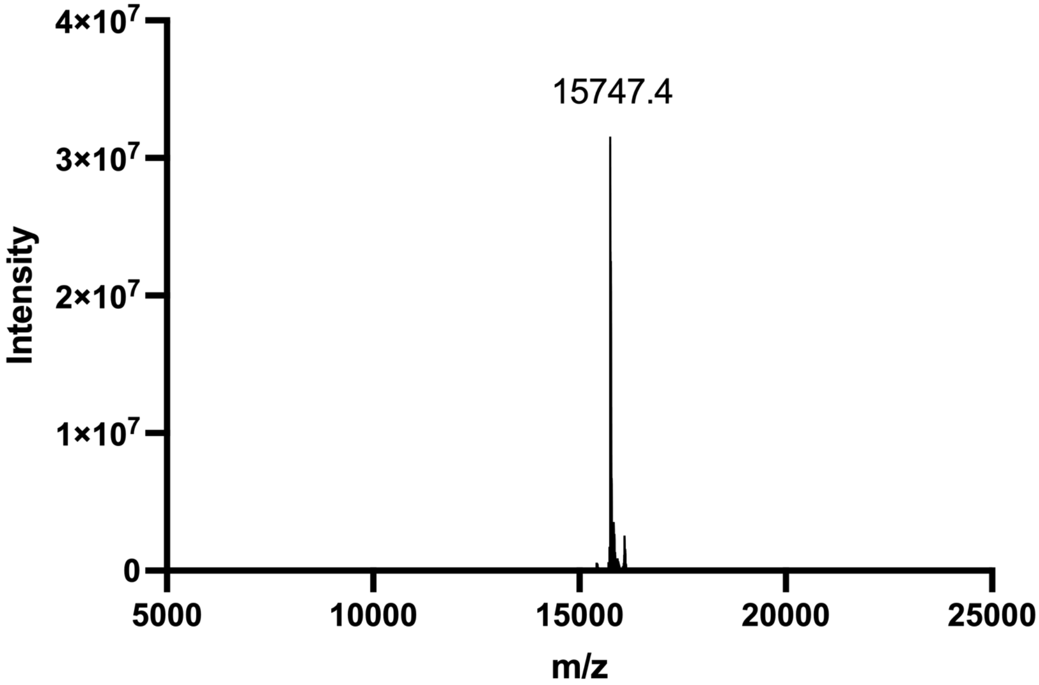


ESI-MS spectrum of **ODN-6-PTC-R**. Calculated MW = 15746.9.

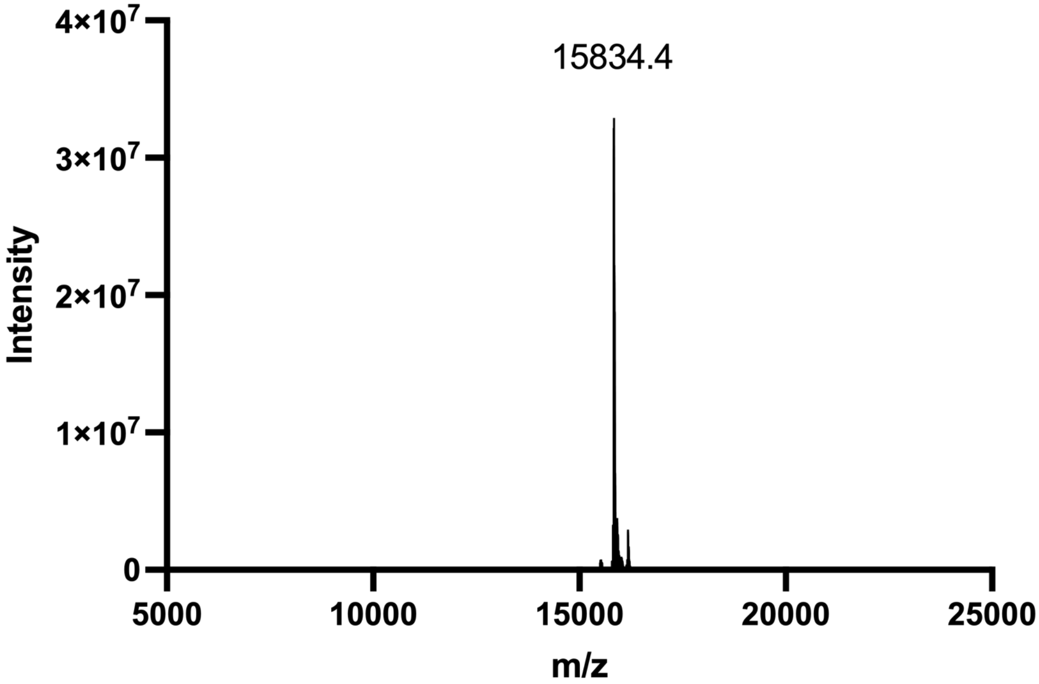


ESI-MS spectrum of **ODN-6-PTC-$^{p}$Y**. Calculated MW = 15833.9.

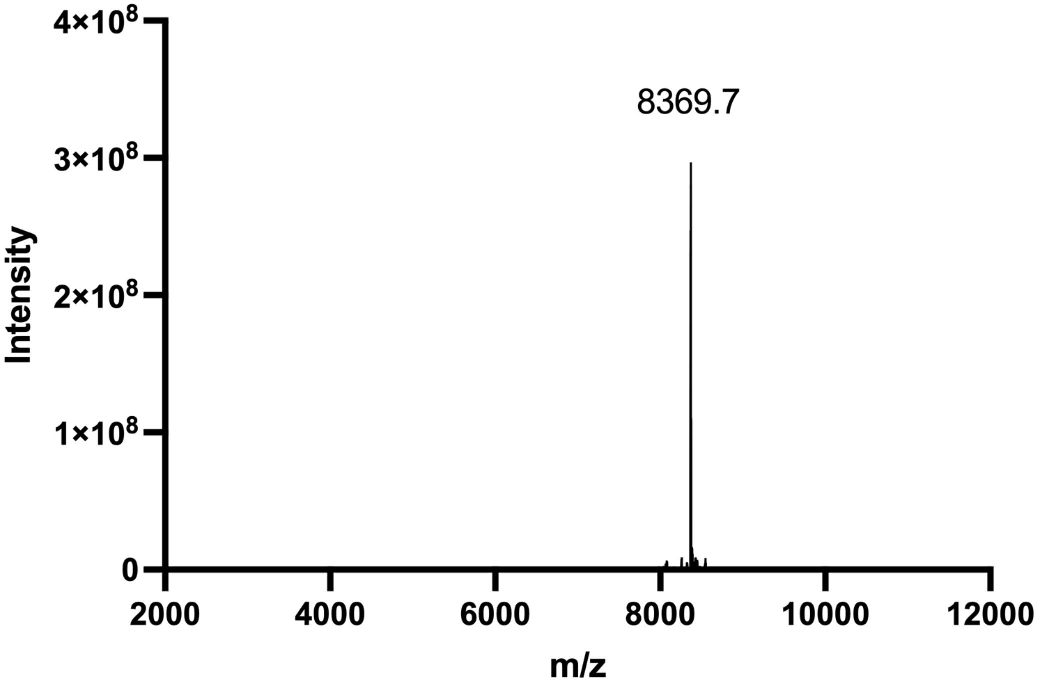


ESI-MS spectrum of **ODN-7**. Calculated MW = 8369.9.

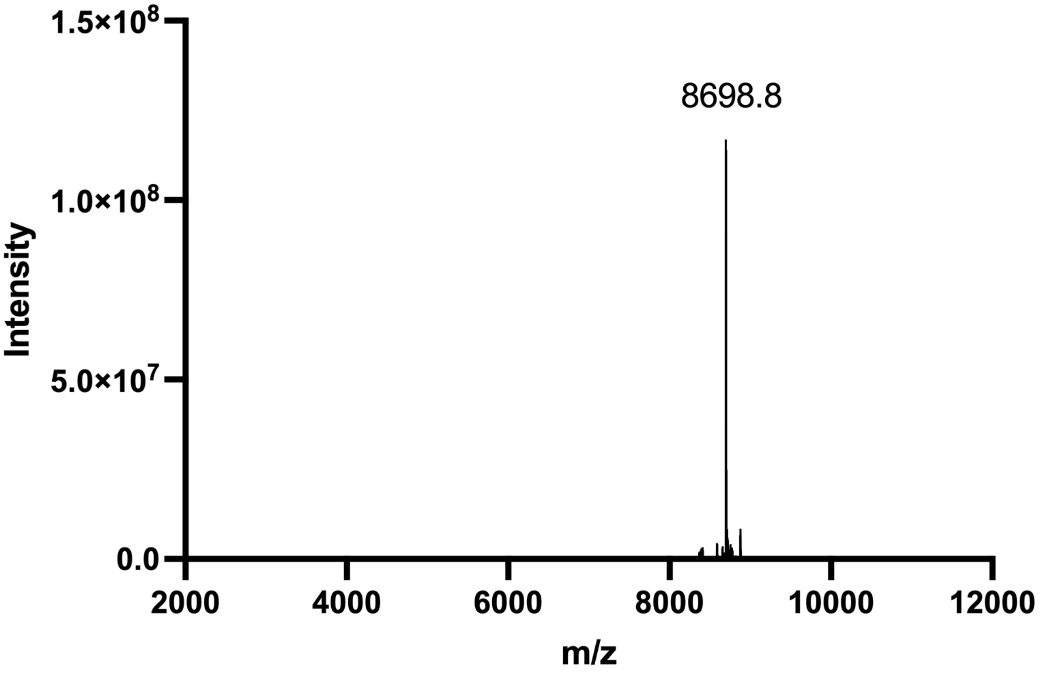


ESI-MS spectrum of **ODN-8**. Calculated MW = 8699.1.

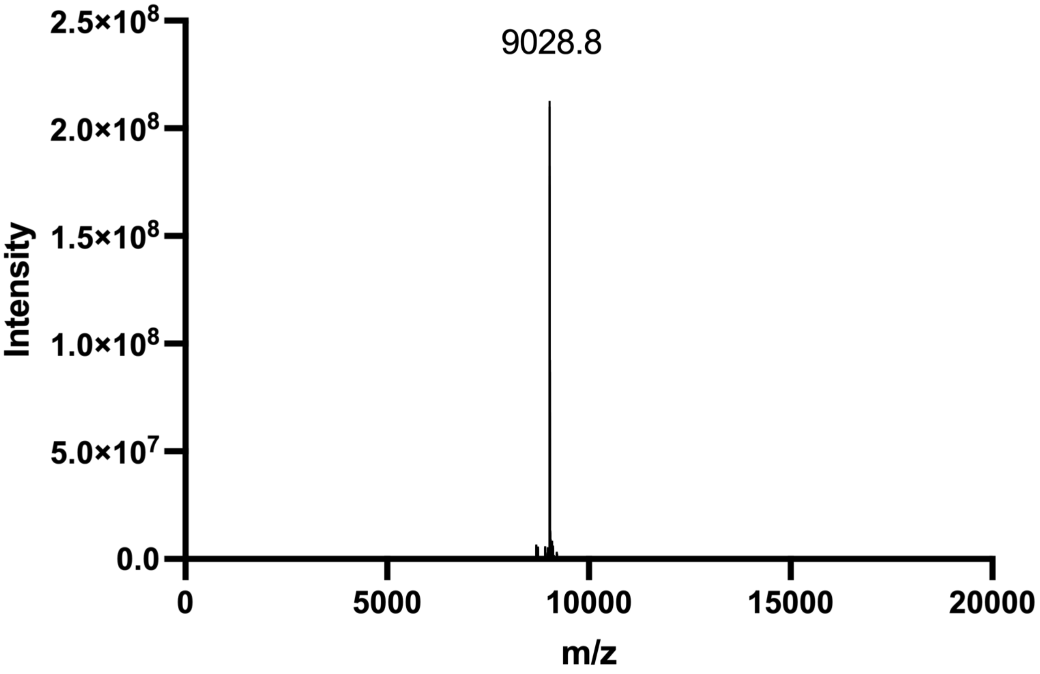


ESI-MS spectrum of **ODN-9**. Calculated MW = 9028.3.

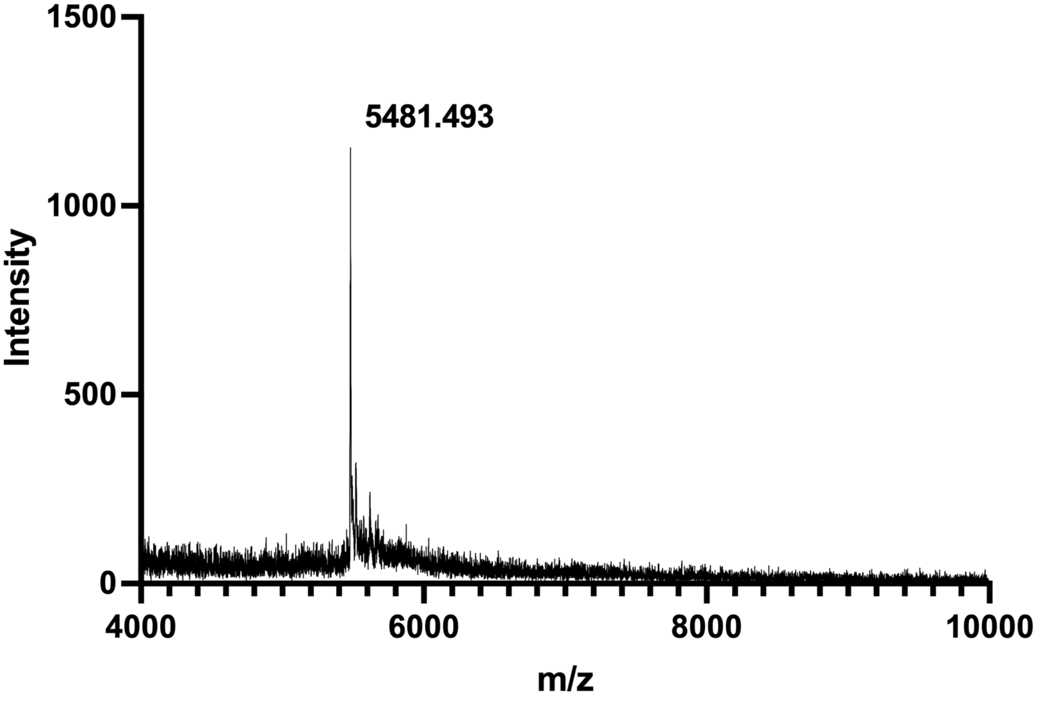


MALDI-TOF-MS spectrum of **ODN-11**. Calculated monoisotopic m/z = 5481.8.

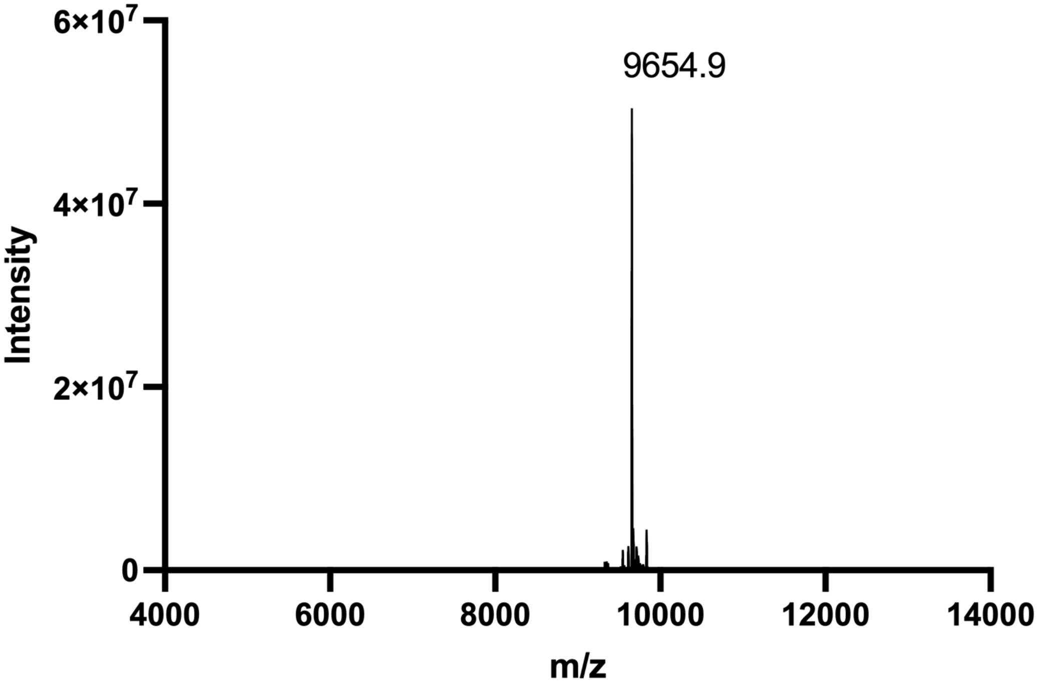


ESI-MS spectrum of **ODN-13**. Calculated MW = 9654.7.

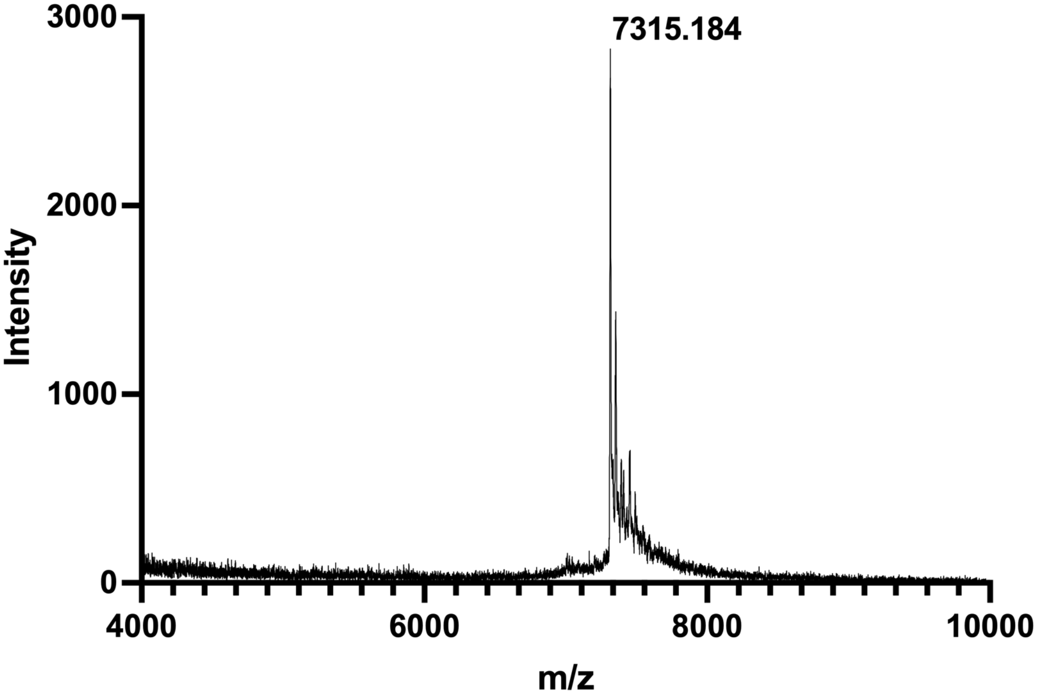


MALDI-TOF-MS spectrum of **ODN-S3**. Calculated monoisotopic m/z = 7315.1.

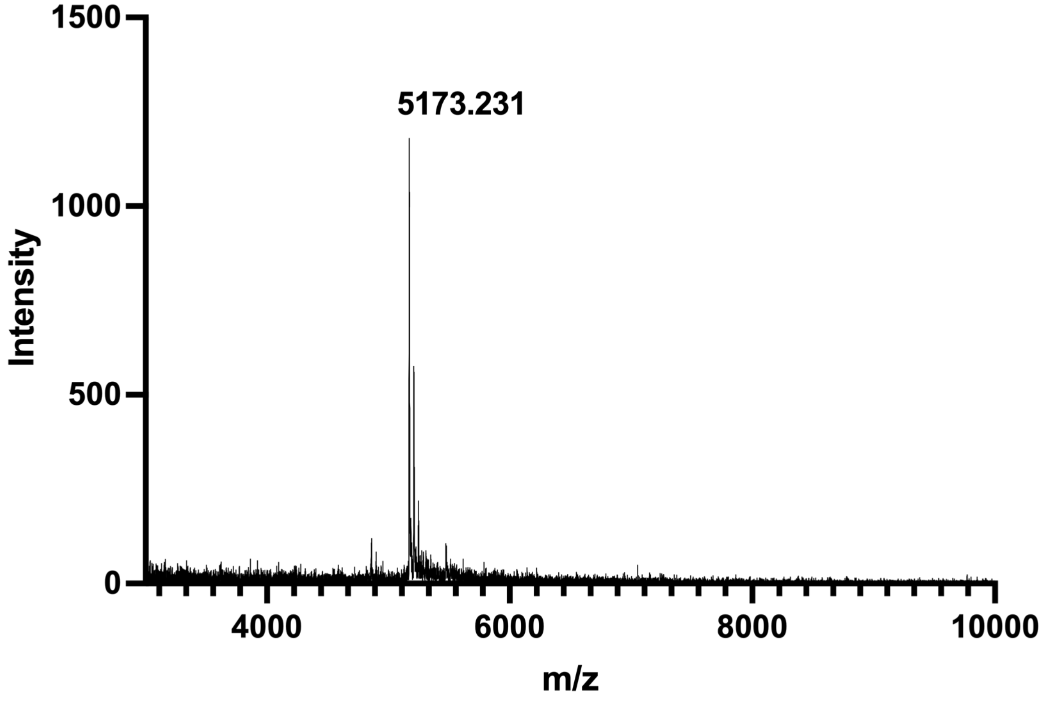


MALDI-TOF-MS spectrum of **ODN-S4**. Calculated monoisotopic m/z =5173.6.

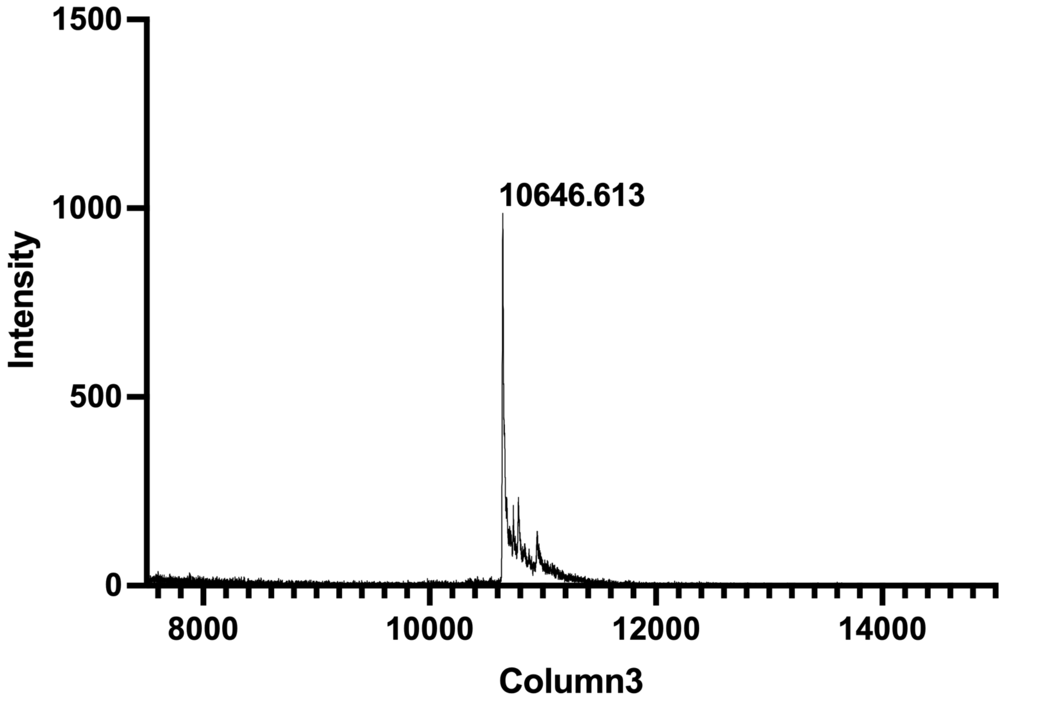


MALDI-TOF-MS spectrum of **ODN-S5**. Calculated monoisotopic m/z = 10646.8.

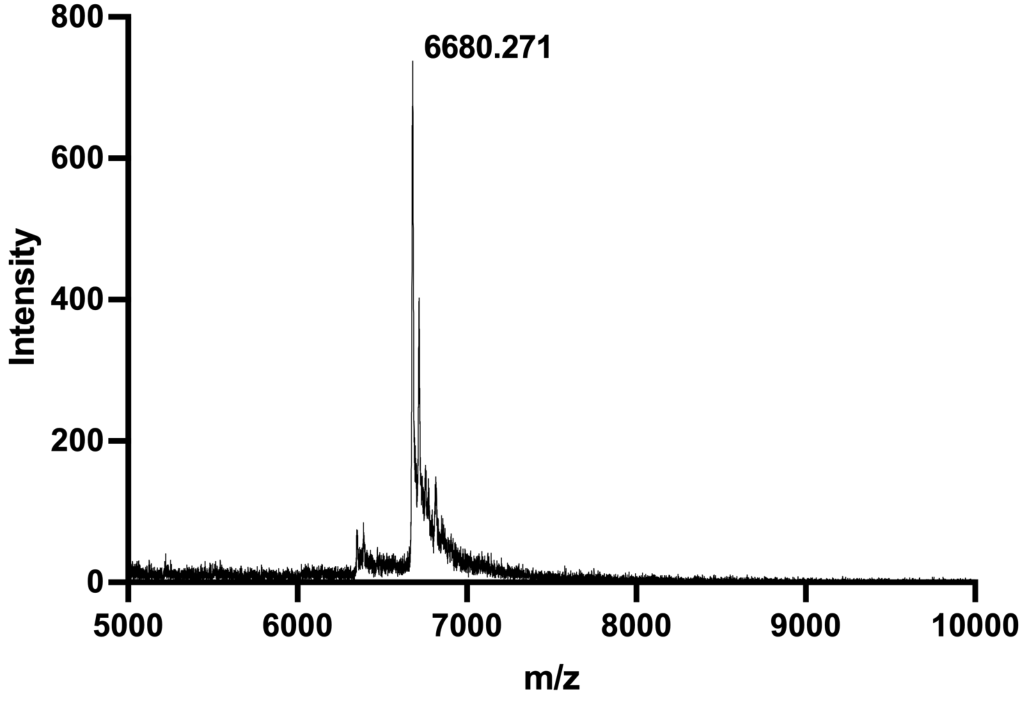


MALDI-TOF-MS spectrum of **ODN-S6**. Calculated monoisotopic m/z = 6680.6.

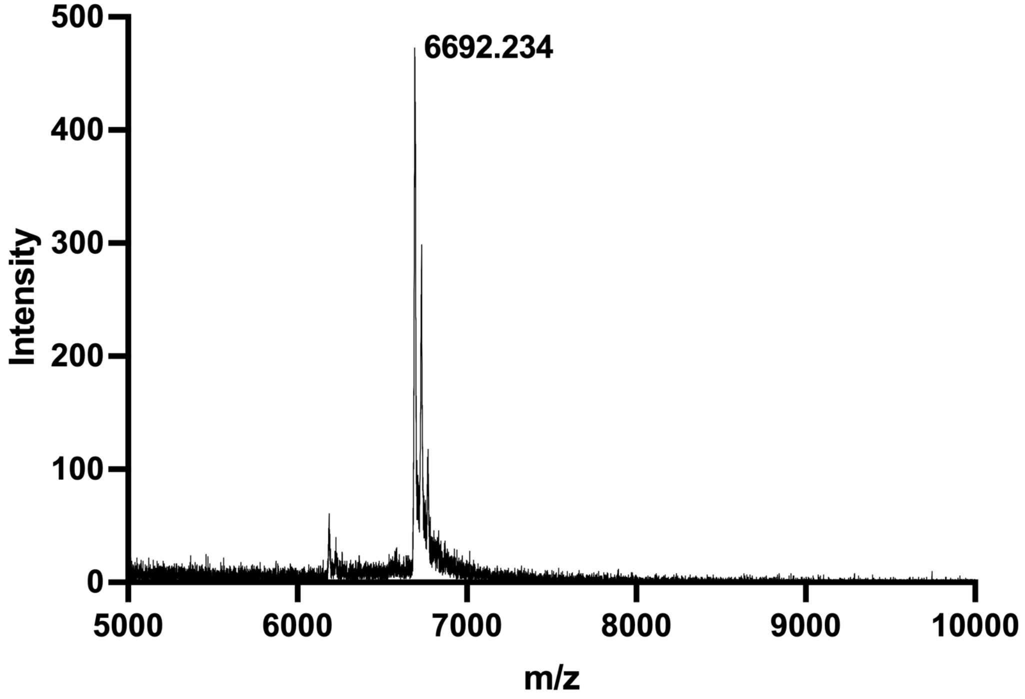


MALDI-TOF-MS spectrum of **ODN-S7**. Calculated monoisotopic m/z = 6691.6.

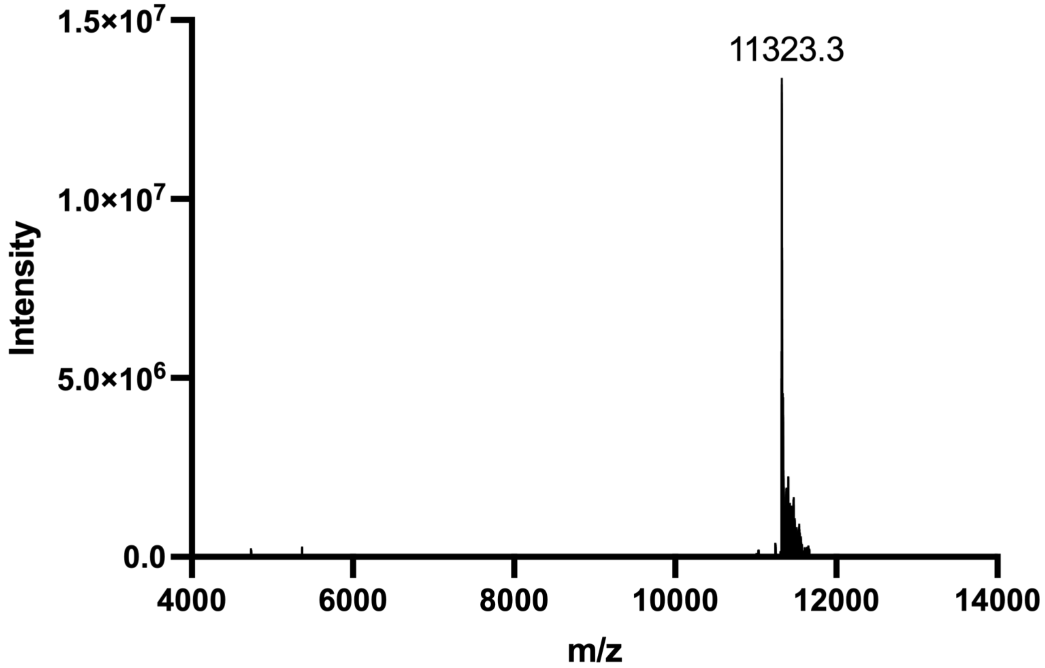


ESI-MS spectrum of **ODN-S8**. Calculated MW = 11322.7.

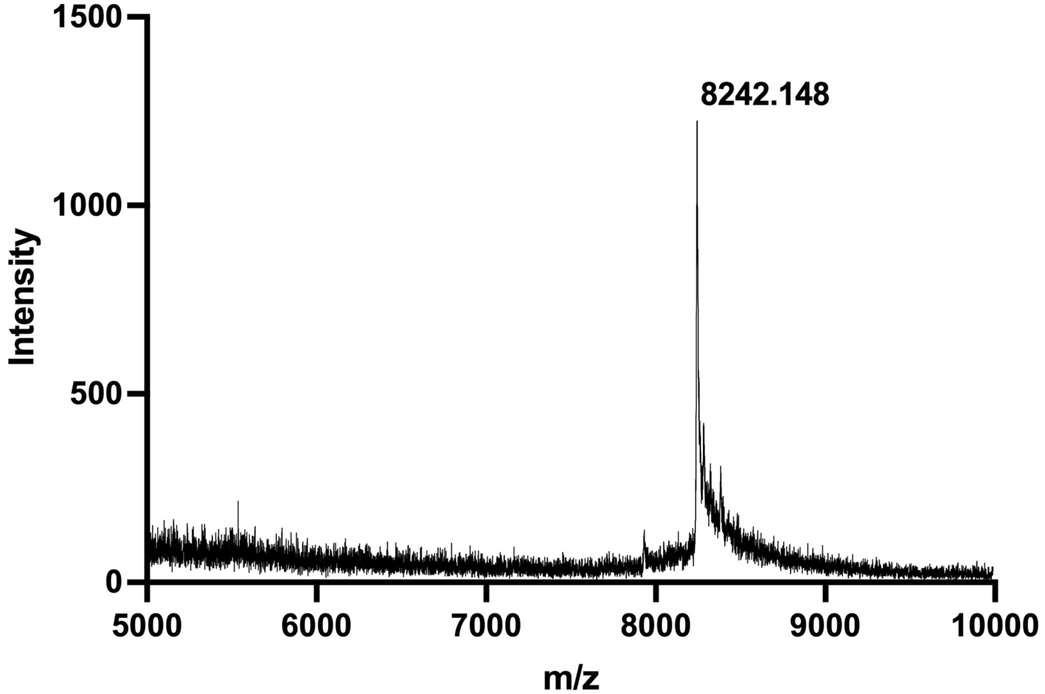


ESI-MS spectrum of **ODN-S10**. Calculated monoisotopic m/z = 8241.9.

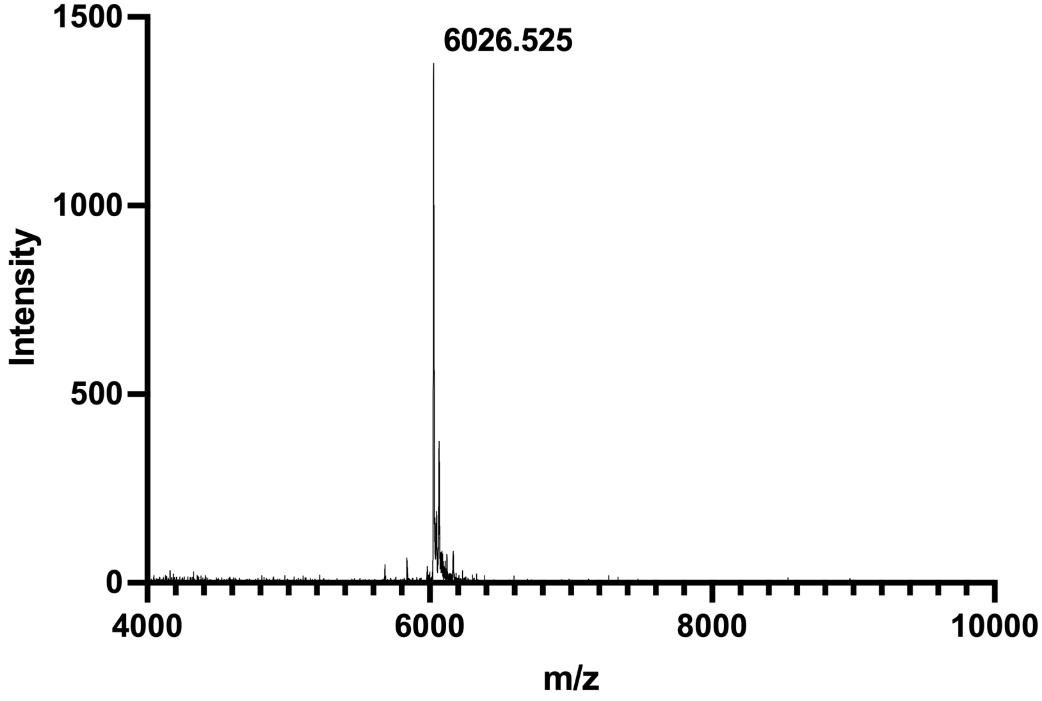


MALDI-TOF-MS spectrum of **ODN-S11**. Calculated monoisotopic m/z = 6026.2.

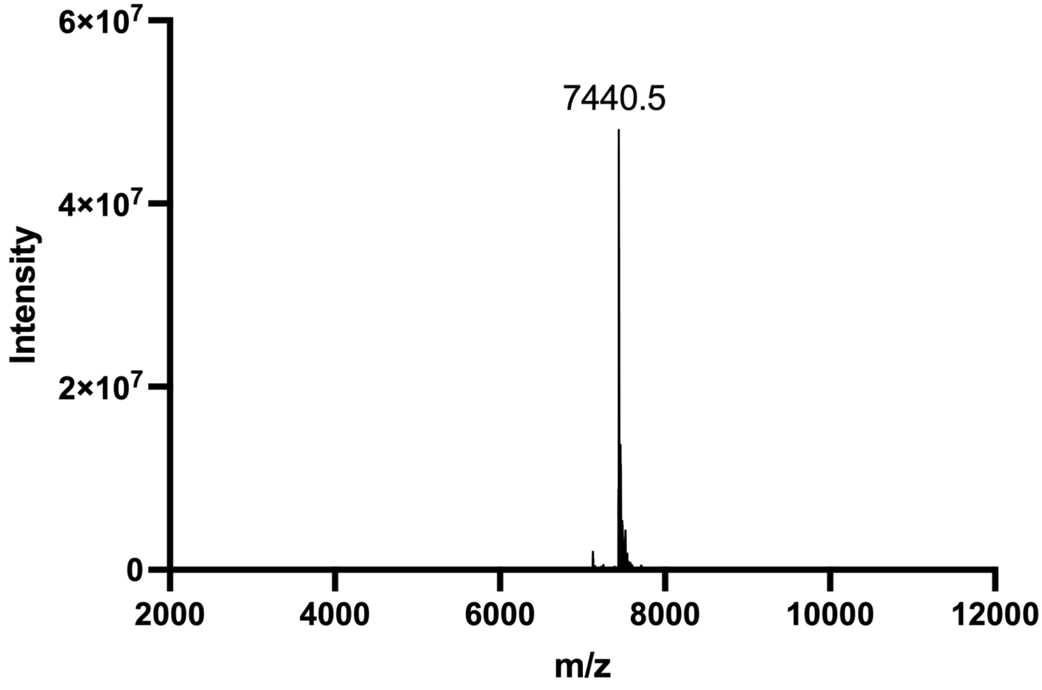


ESI-MS spectrum of **ODN-S12**. Calculated MW = 7440.6.

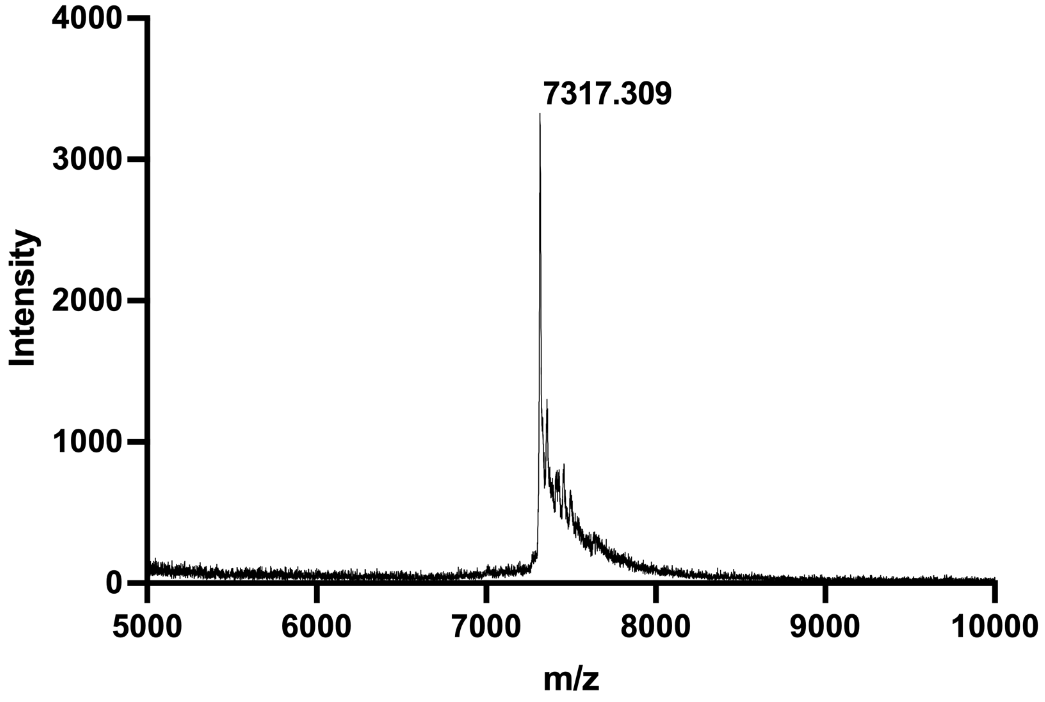


MALDI-TOF-MS spectrum of **ODN-S13**. Calculated monoisotopic m/z = 7316.9.

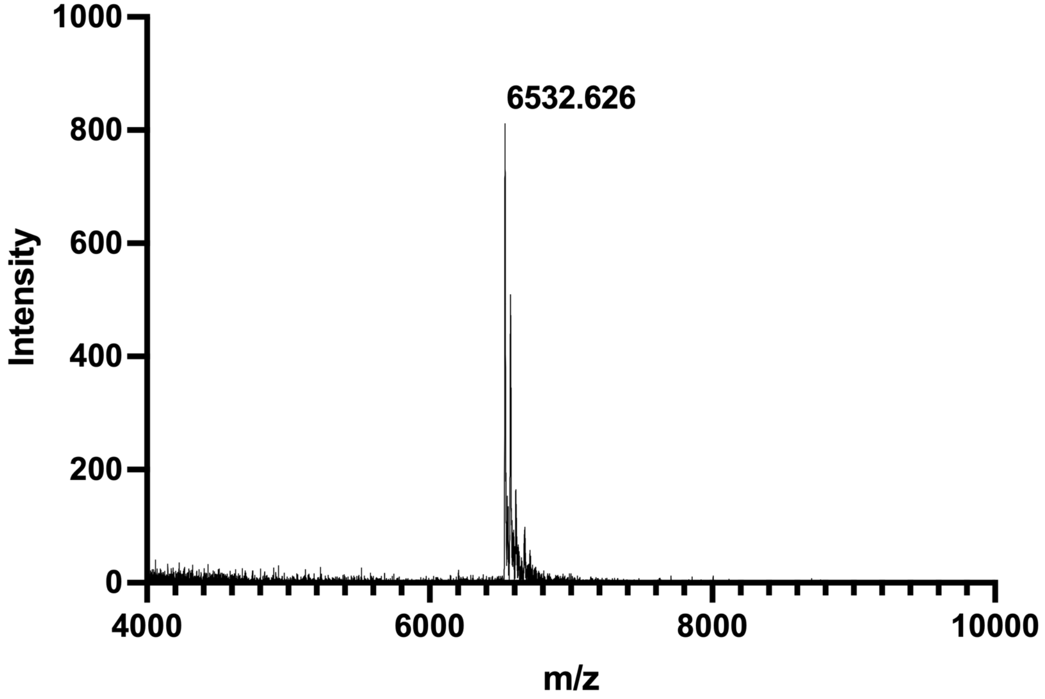


MALDI-TOF-MS spectrum of **ODN-S14**. Calculated monoisotopic m/z = 6532.7.

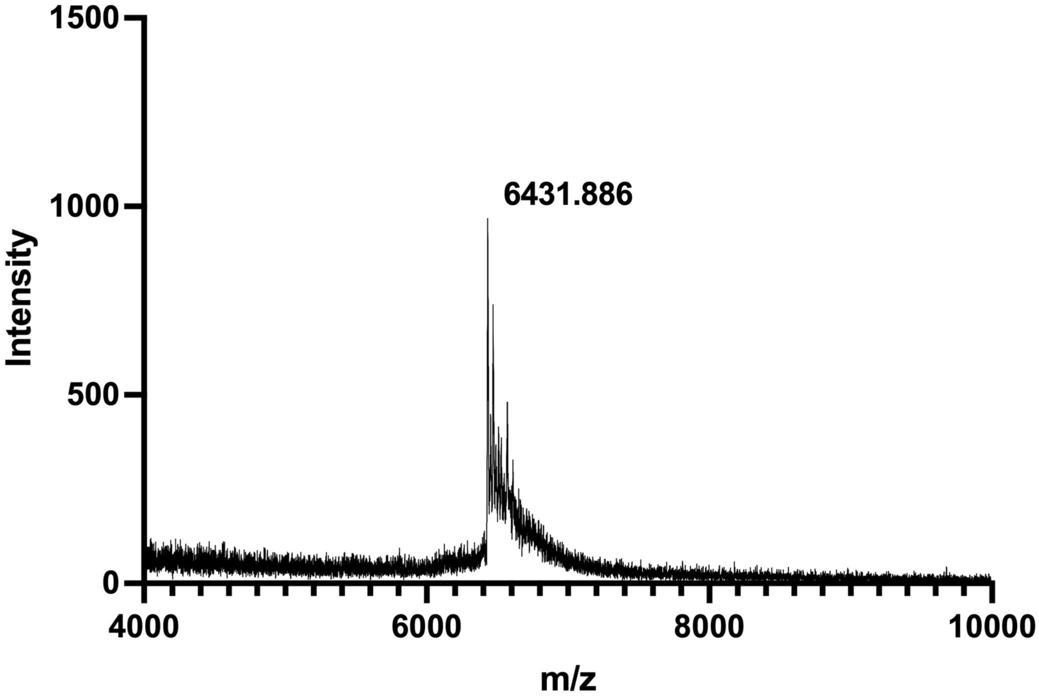

MALDI-TOF-MS spectrum of **ODN-S15**. Calculated monoisotopic m/z = 6431.6.

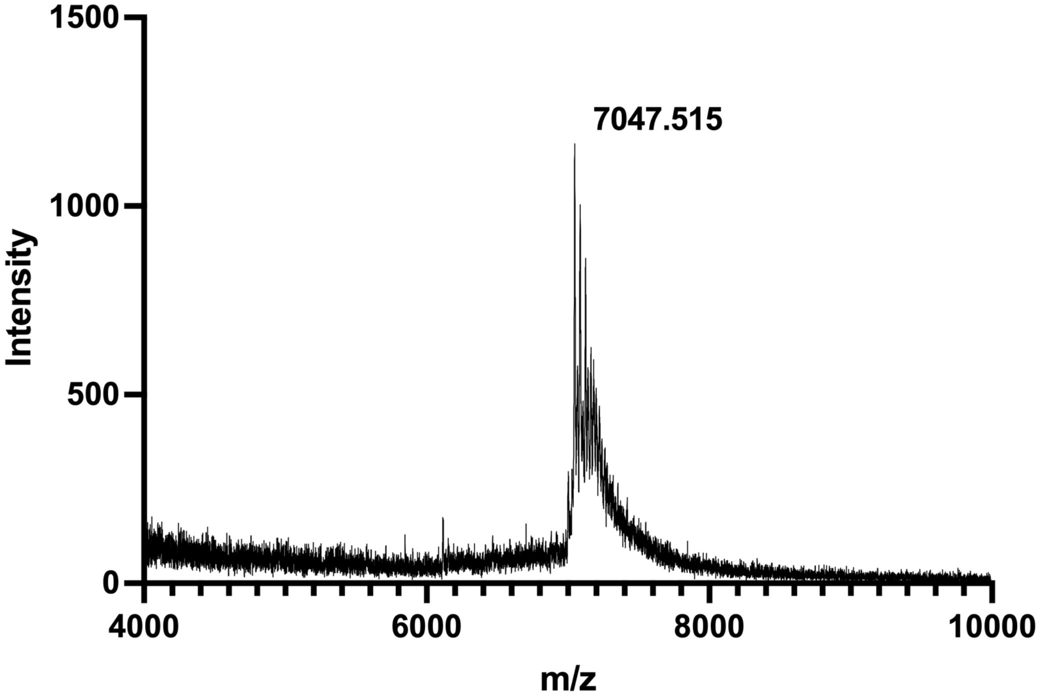

MALDI-TOF-MS spectrum of **ODN-S15-PTC-F**. Calculated monoisotopic m/z = 7047.8.

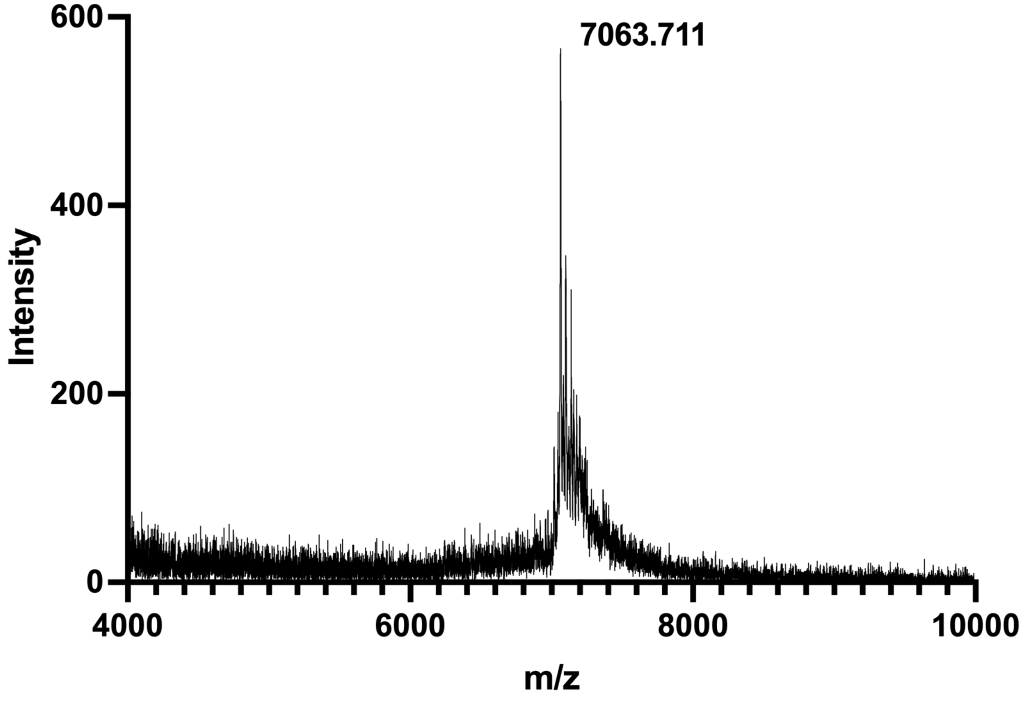

MALDI-TOF-MS spectrum of **ODN-S15-PTC-Y**. Calculated monoisotopic m/z = 7063.8.

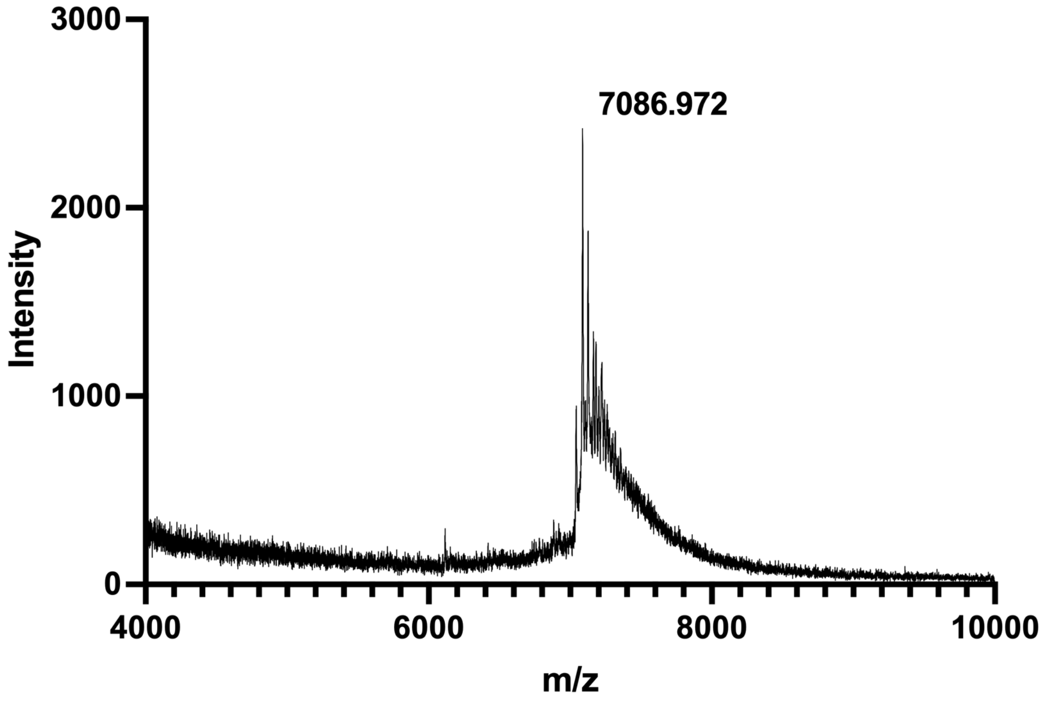

MALDI-TOF-MS spectrum of **ODN-S15-PTC-W**. Calculated monoisotopic m/z = 7086.8.

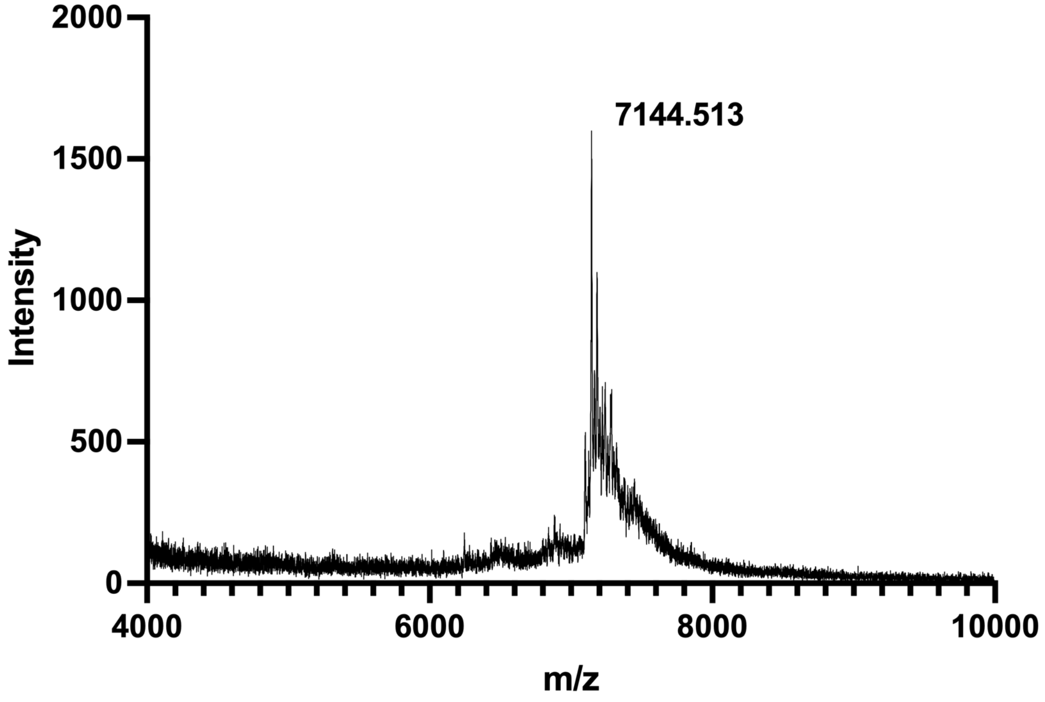


MALDI-TOF-MS spectrum of **ODN-S15-PTC-pY**. Calculated monoisotopic m/z = 7143.8.

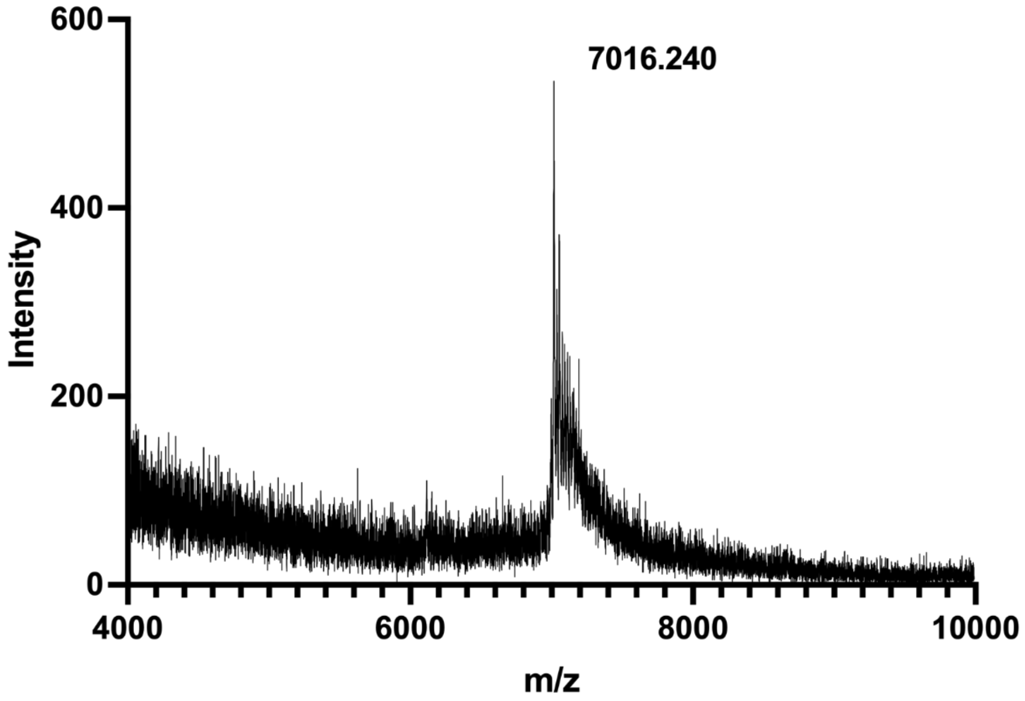


MALDI-TOF-MS spectrum of **ODN-S15-PTC-D**. Calculated monoisotopic m/z = 7015.8.

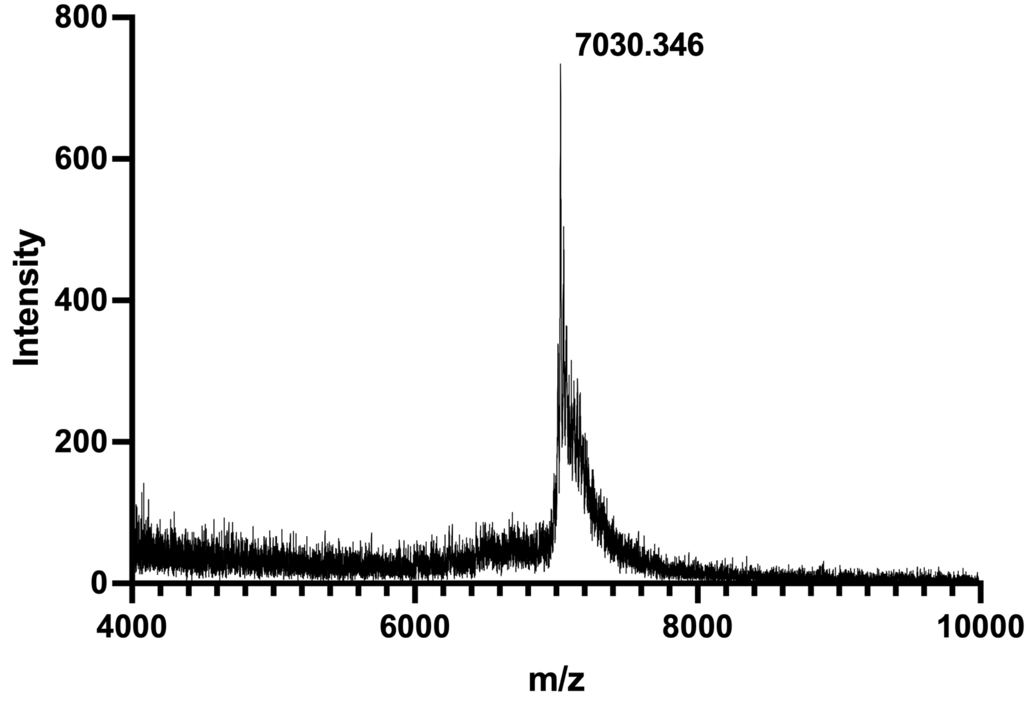

MALDI-TOF-MS spectrum of **ODN-S15-PTC-E**. Calculated monoisotopic m/z = 7030.8.

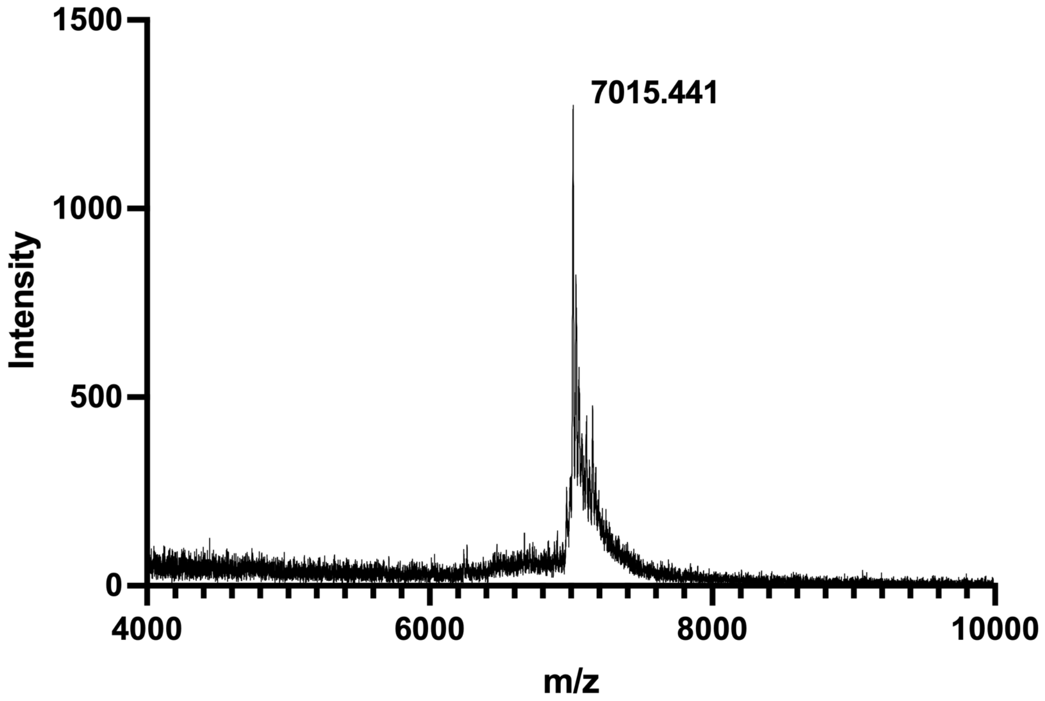

MALDI-TOF-MS spectrum of **ODN-S15-PTC-N**. Calculated monoisotopic m/z = 7014.8.

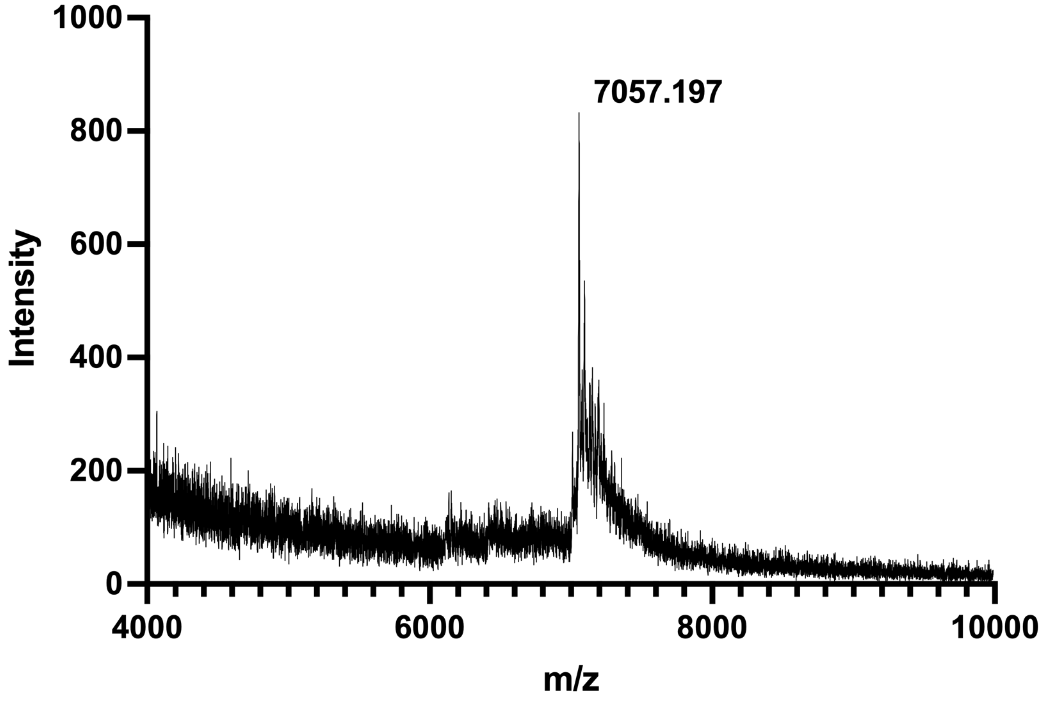


MALDI-TOF-MS spectrum of **ODN-S15-PTC-R**. Calculated monoisotopic m/z = 7056.9.

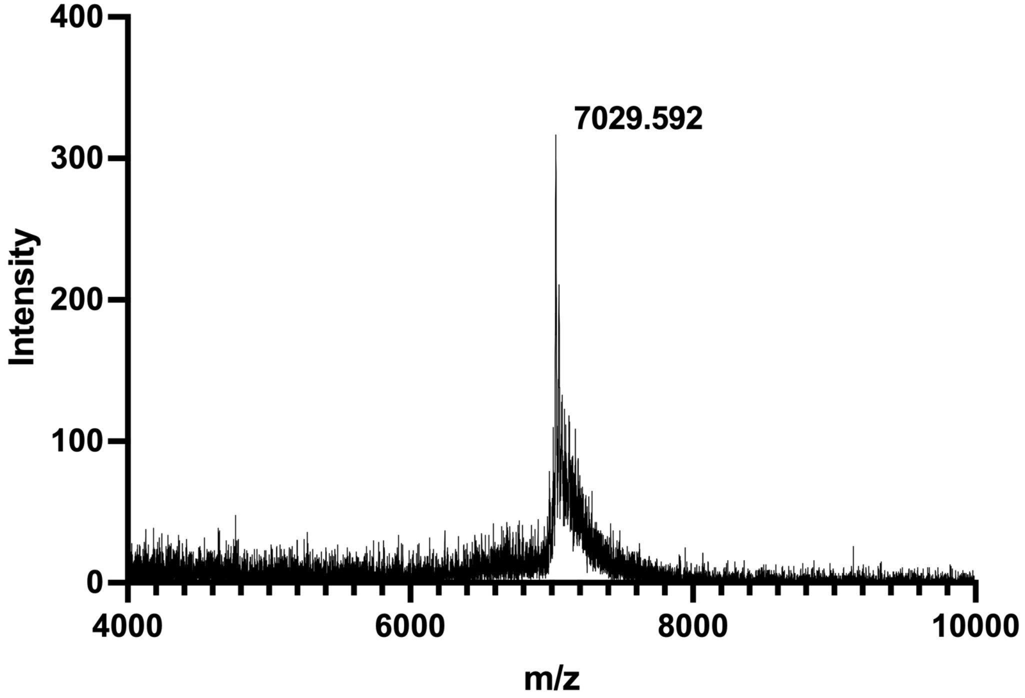


MALDI-TOF-MS spectrum of **ODN-S15-PTC-Q**. Calculated monoisotopic m/z = 7029.8.

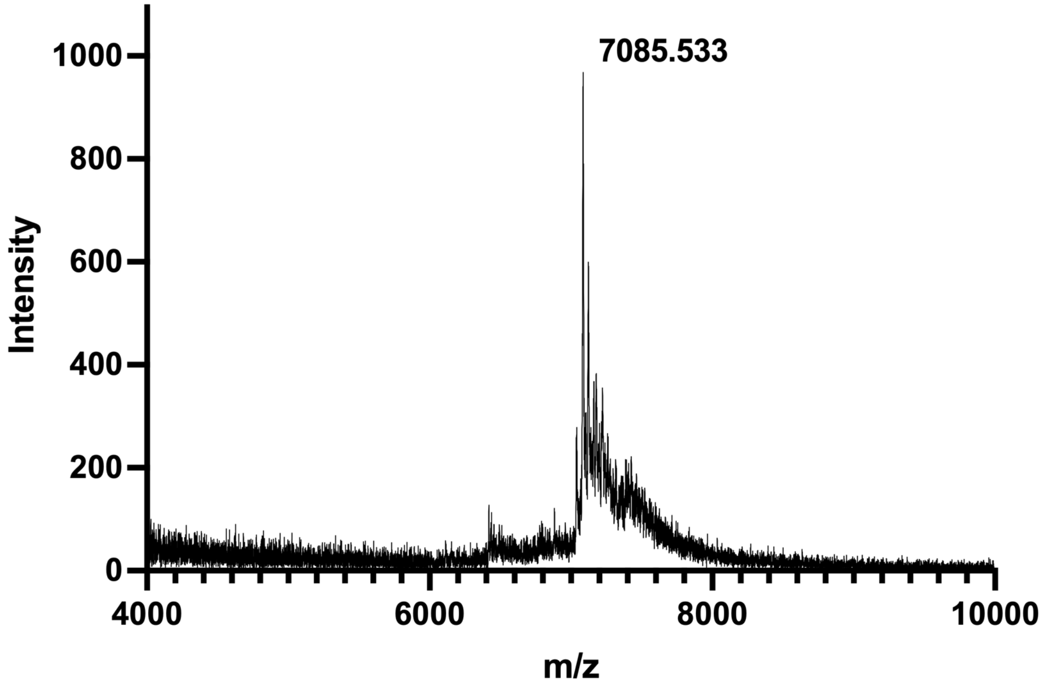


MALDI-TOF-MS spectrum of **ODN-S15-PTC-ADMA**. Calculated monoisotopic m/z = 7084.9.

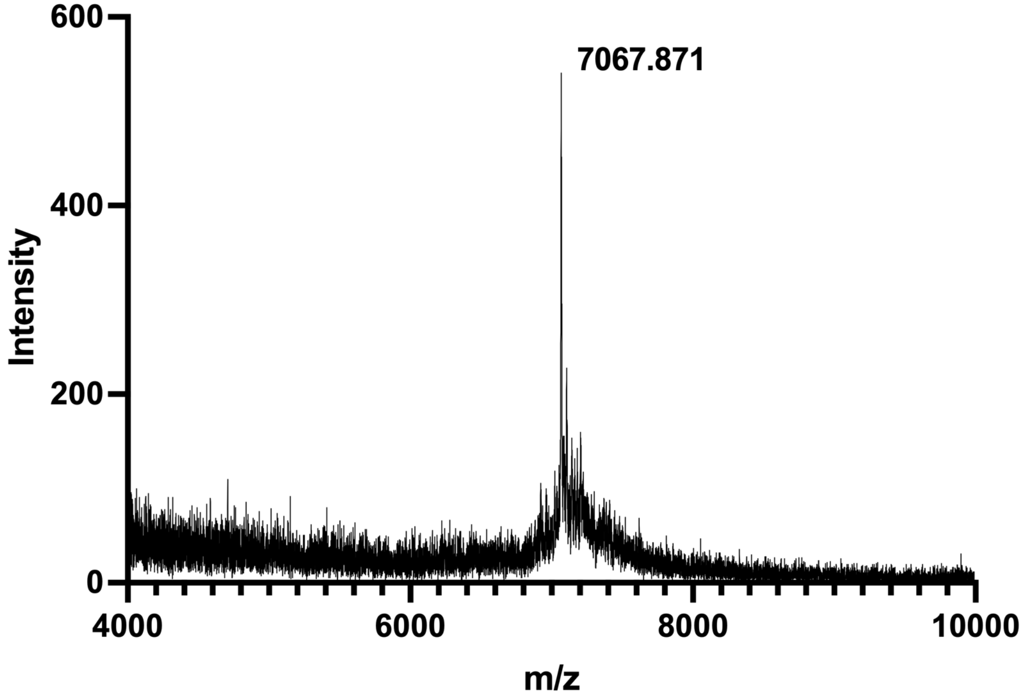


MALDI-TOF-MS spectrum of **ODN-S15-PTC-$^{p}$S**. Calculated monoisotopic m/z = 7067.8.

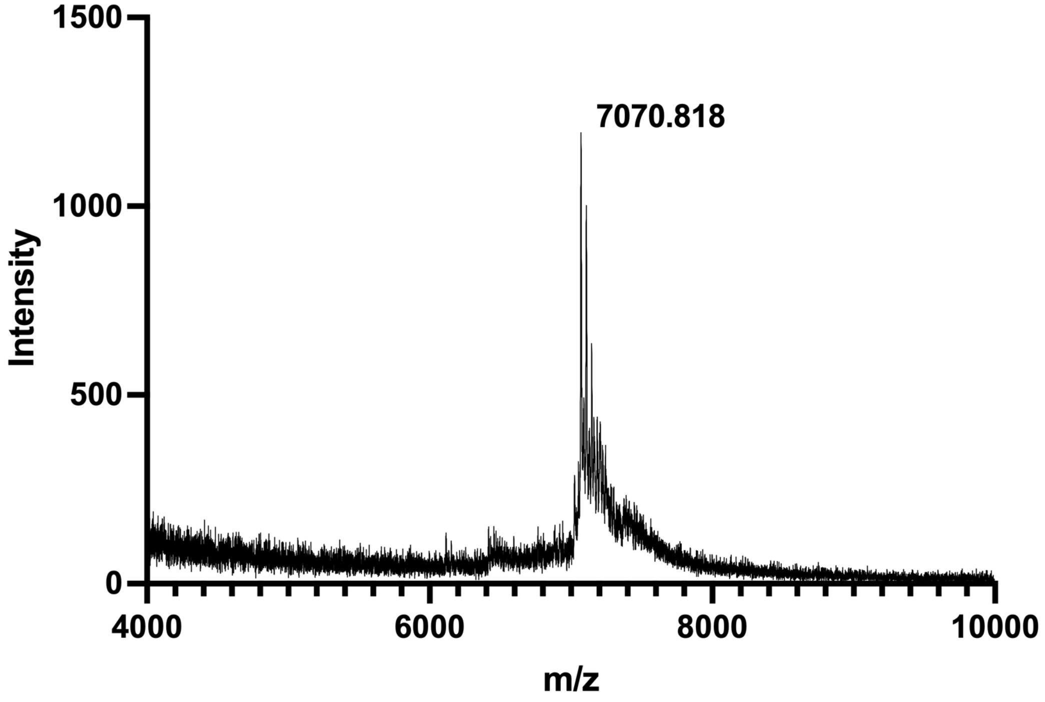


MALDI-TOF-MS spectrum of **ODN-S15-PTC-$^{Ac}$K**. Calculated monoisotopic m/z = 7070.9.

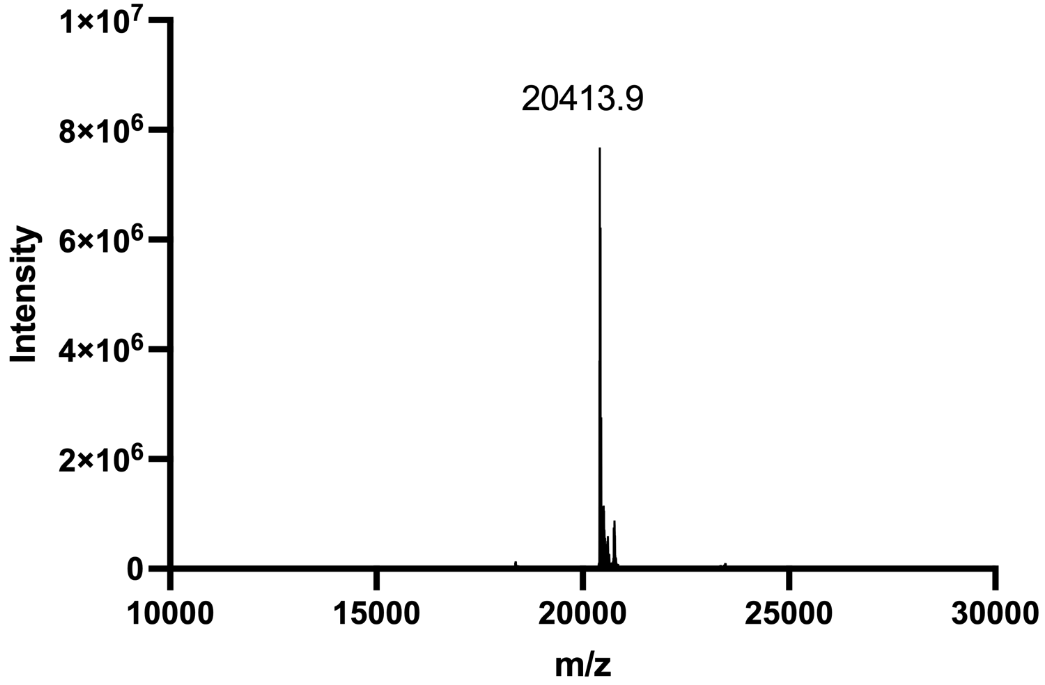


ESI-MS spectrum of **ODN-S16**. Calculated MW = 20412.9.

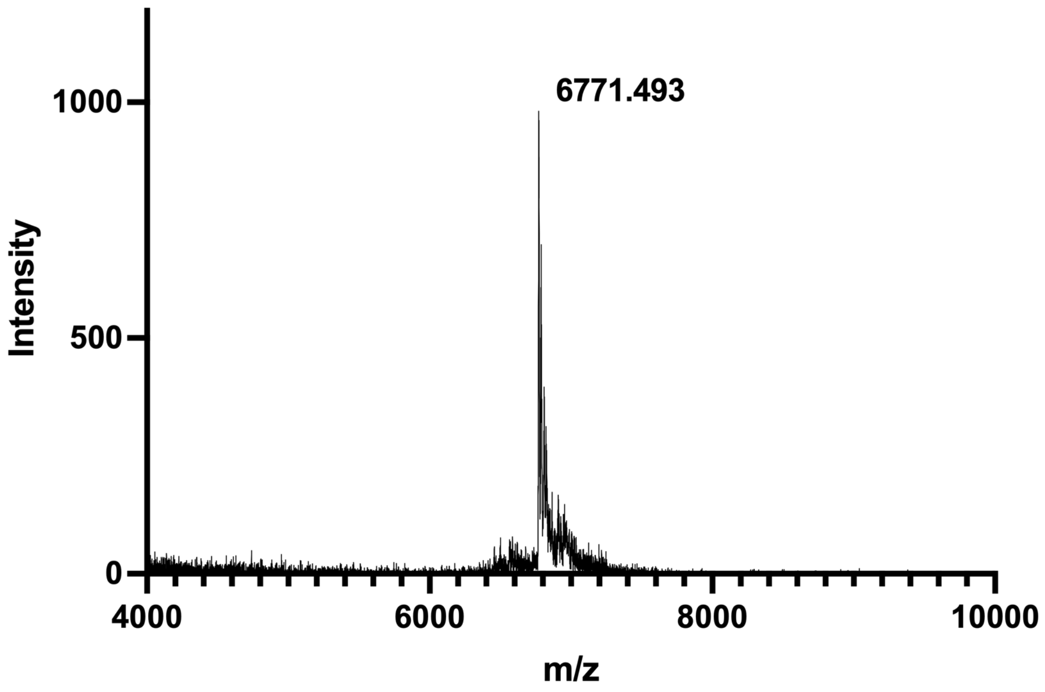


ESI-MS spectrum of **ODN-S22**. Calculated monoisotopic m/z = 6770.9.

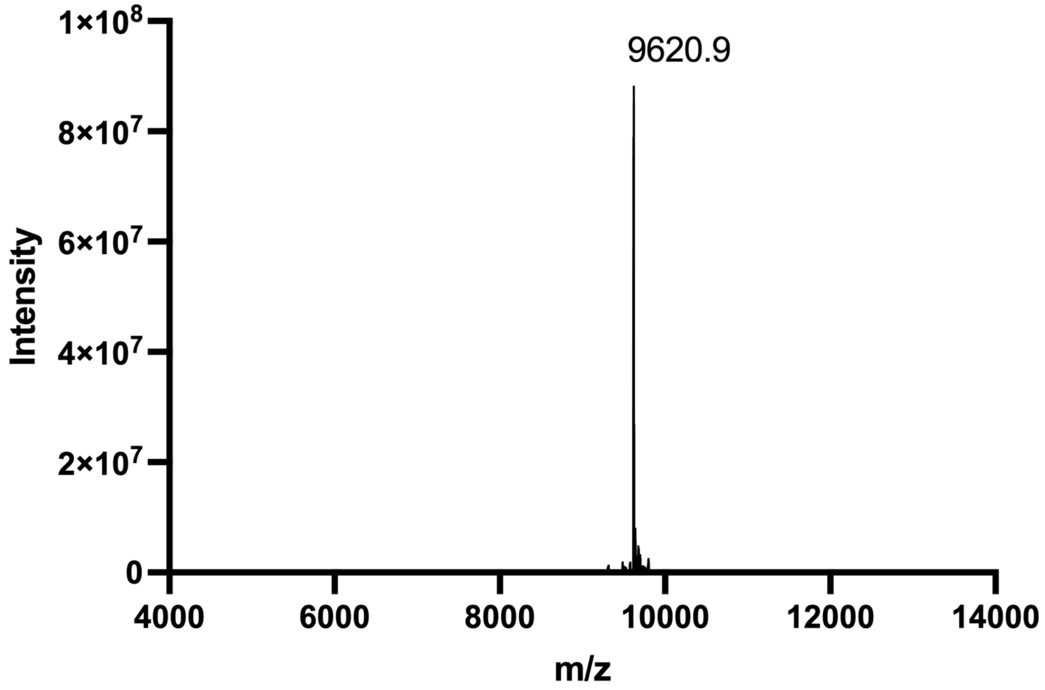


ESI-MS spectrum of **ODN-S23**. Calculated MW =9620.7.

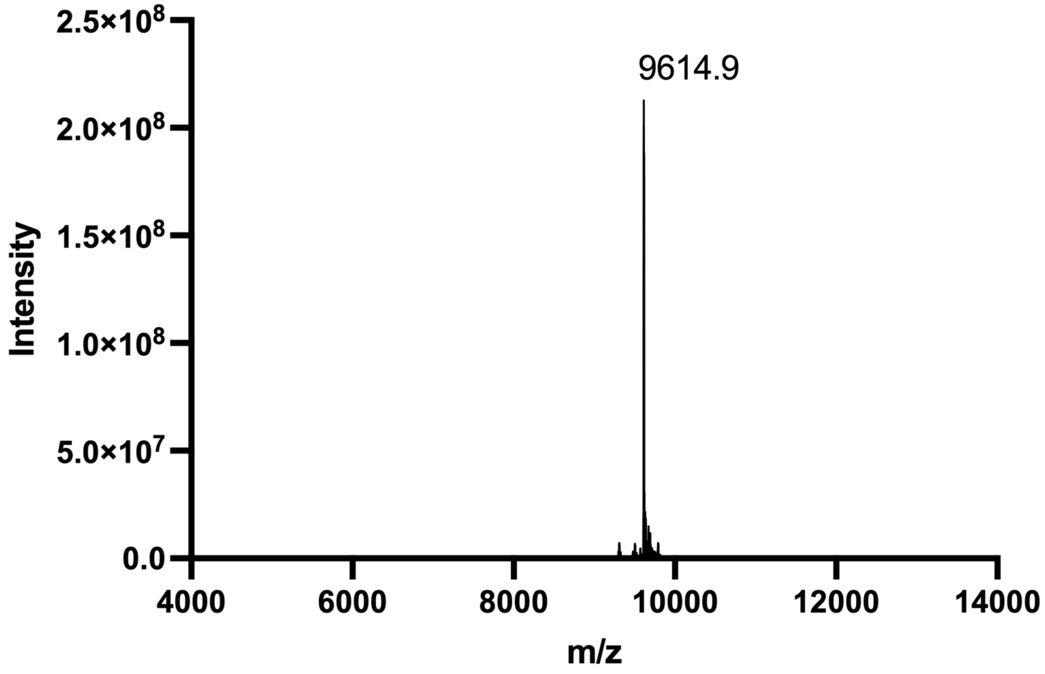


ESI-MS spectrum of **ODN-S24**. Calculated MW = 9614.7.

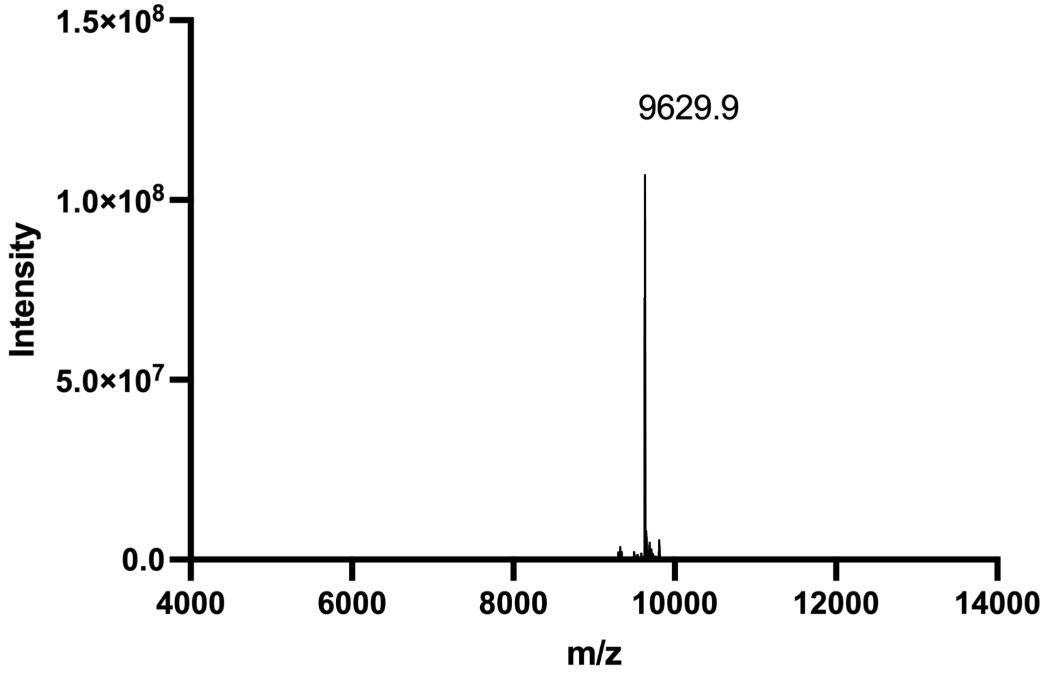


ESI-MS spectrum of **ODN-S25**. Calculated MW = 9629.7.

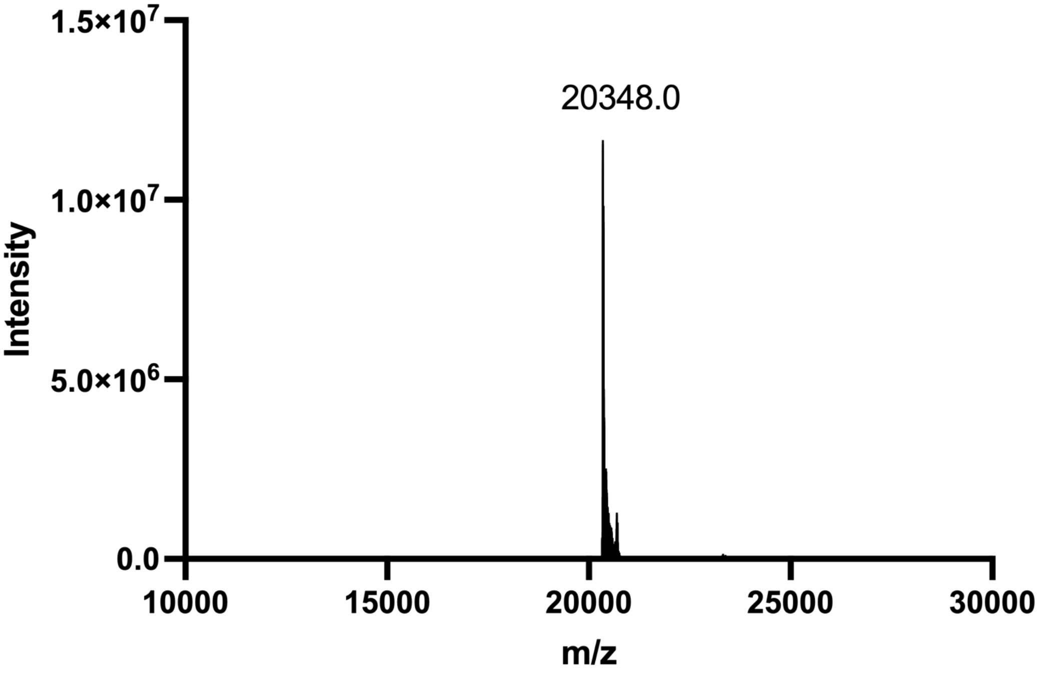


ESI-MS spectrum of **ODN-S31**. Calculated MW = 20347.8.

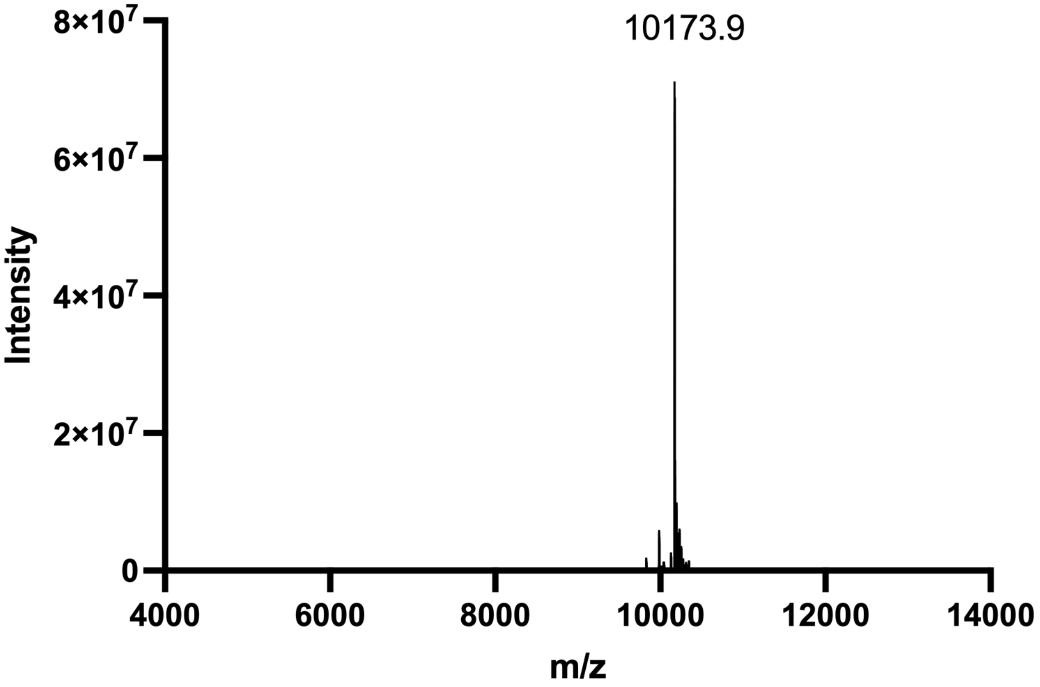


ESI-MS spectrum of **ODN-S32**. Calculated MW = 10174.5.

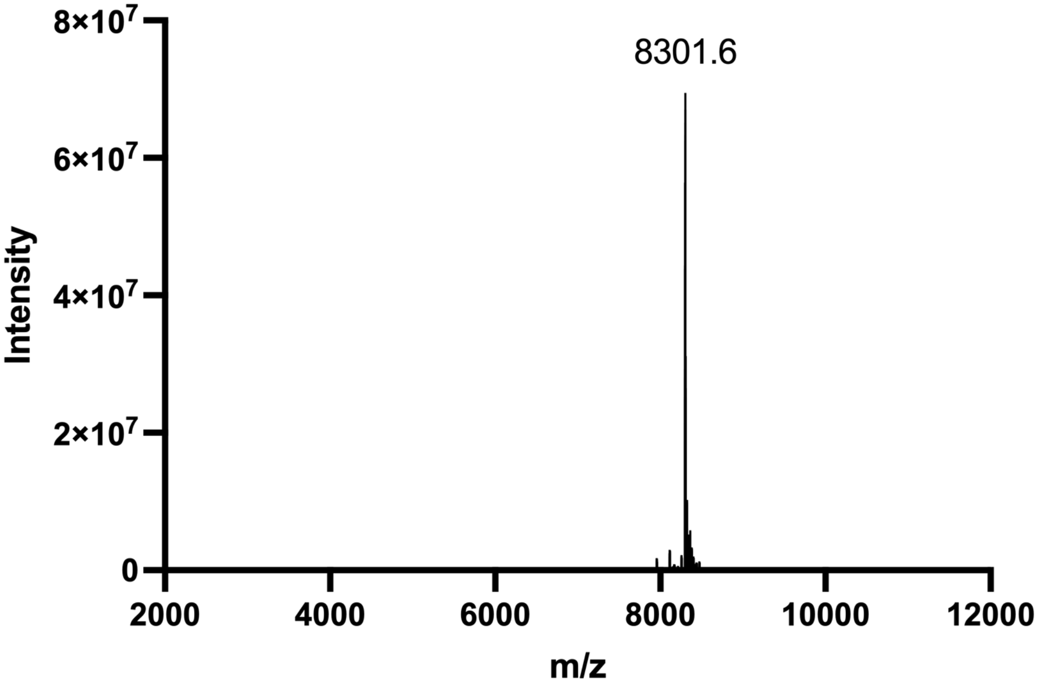


ESI-MS spectrum of **ODN-S33**. Calculated MW = 8301.3.

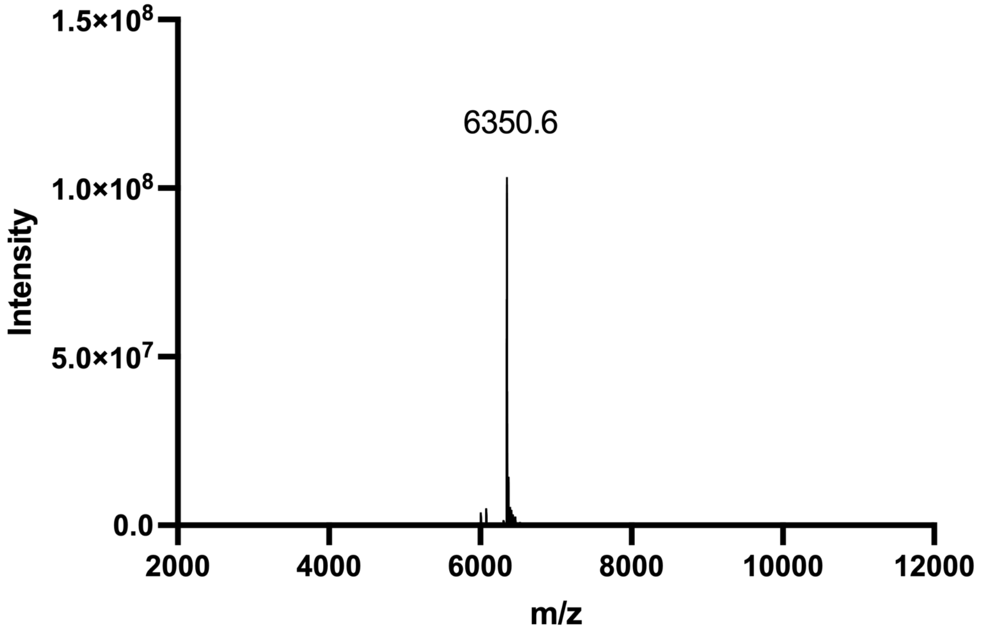


ESI-MS spectrum of **ODN-S34**. Calculated MW = 6350.8.

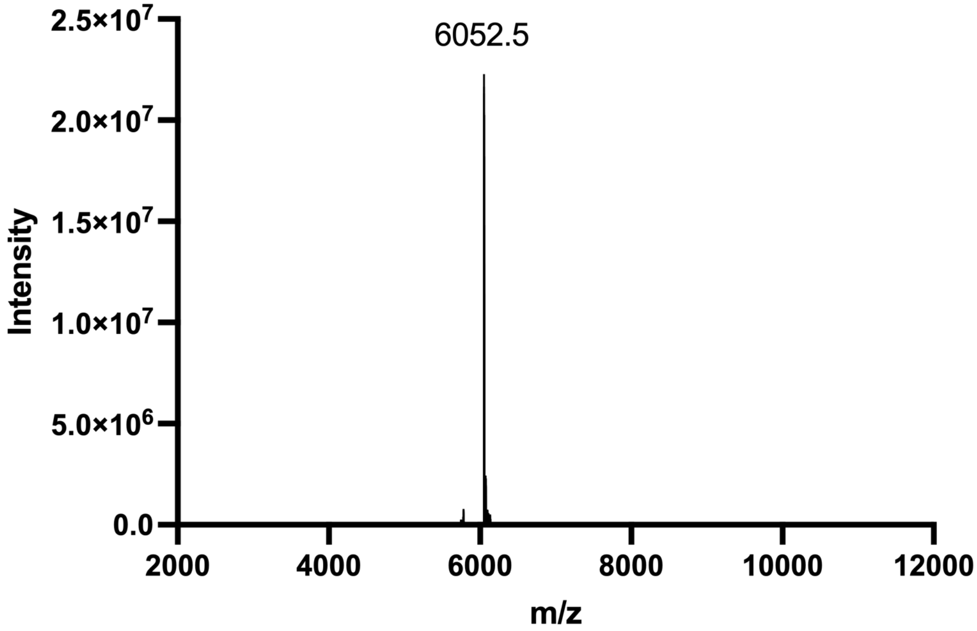


ESI-MS spectrum of **ODN-S35**. Calculated MW = 6052.5.

### NMR spectra of organic compounds

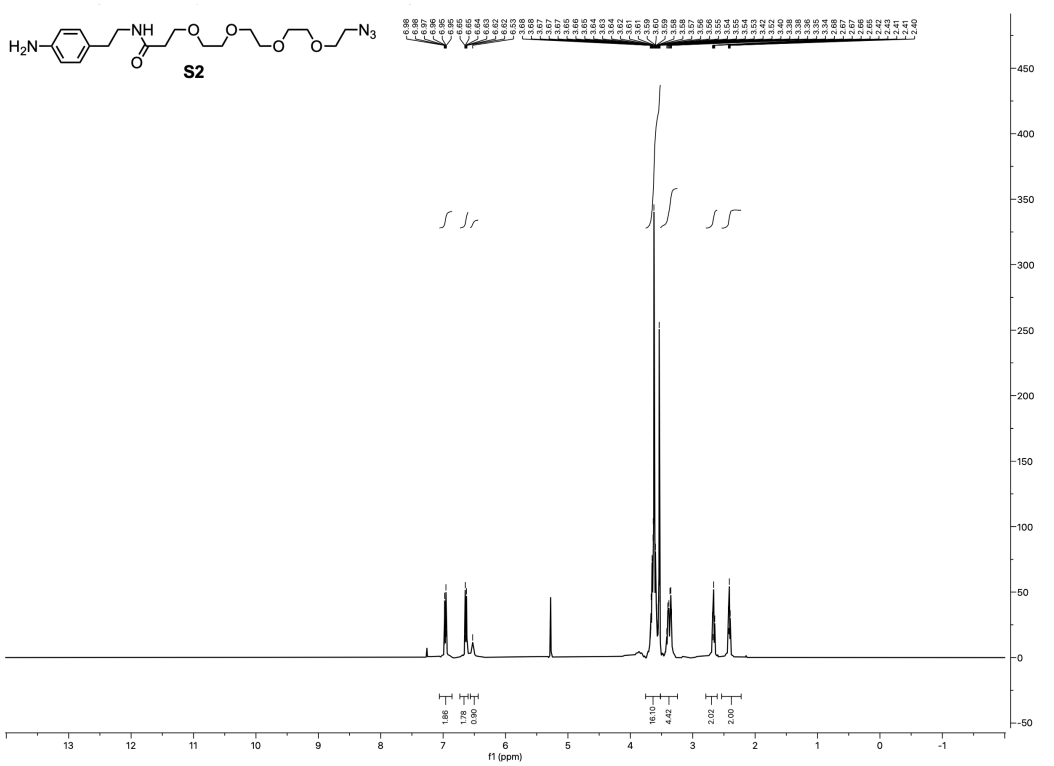


$^{1}H$ NMR (400 MHz, $CDCl_3$) spectrum of **S2**.

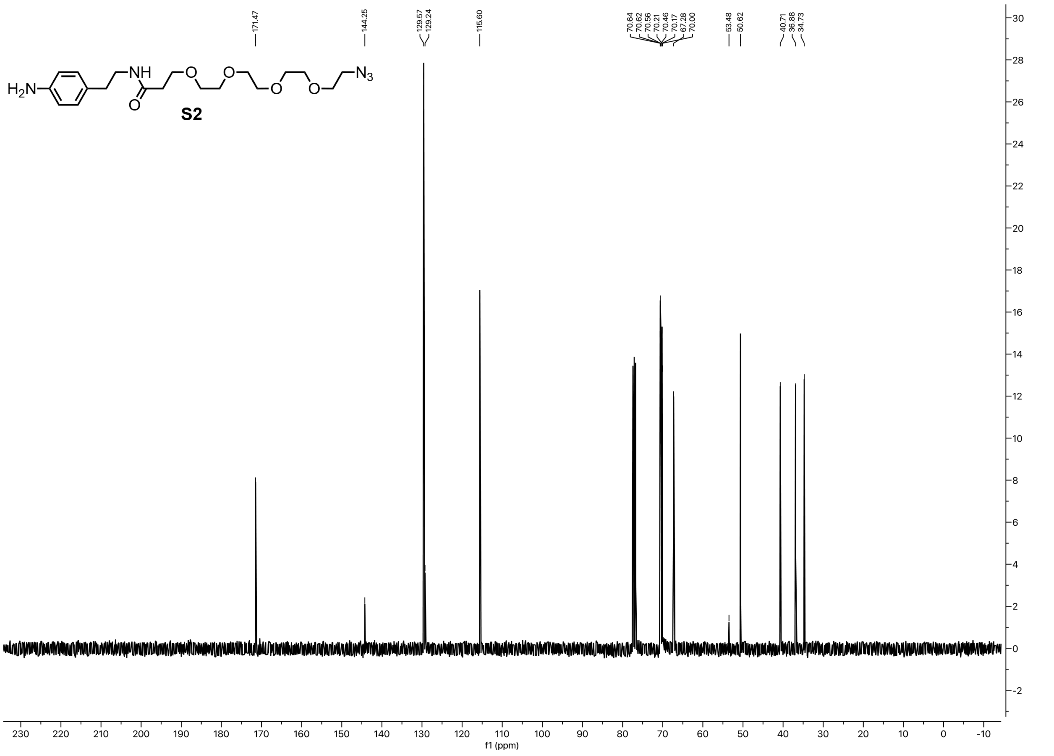


$^{13}C$ NMR (101 MHz, $CDCl_3$) spectrum of **S2**

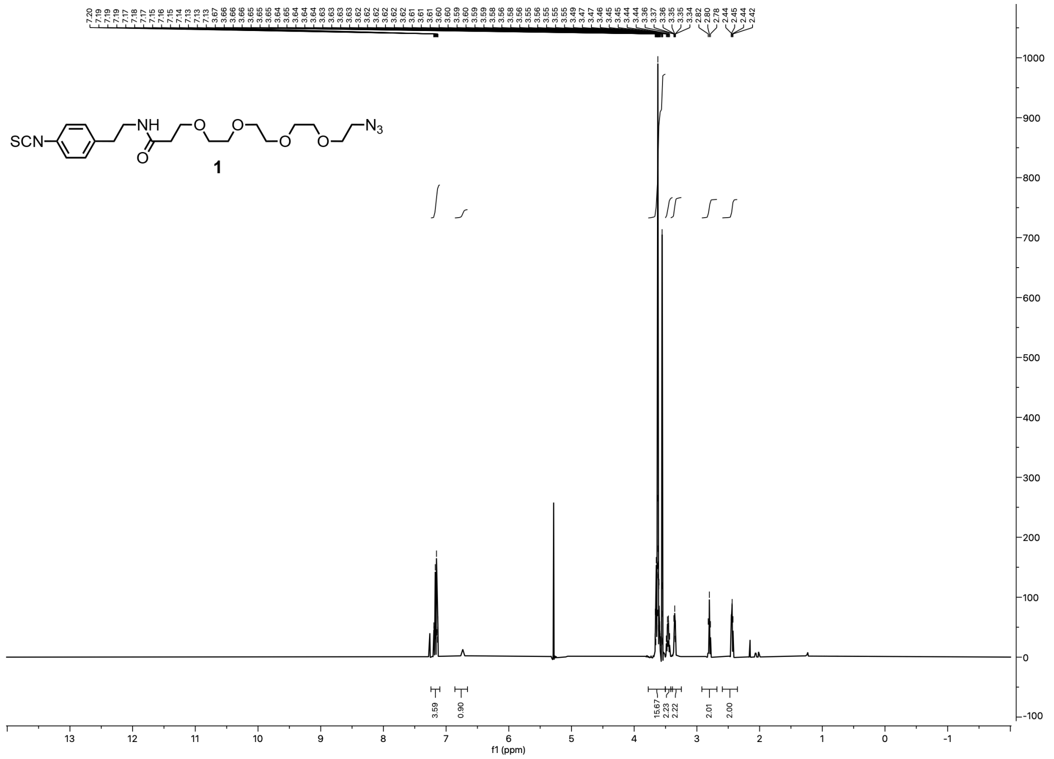

$^{1}$H NMR (400 MHz, $CDCl_3$) spectrum of **1**.

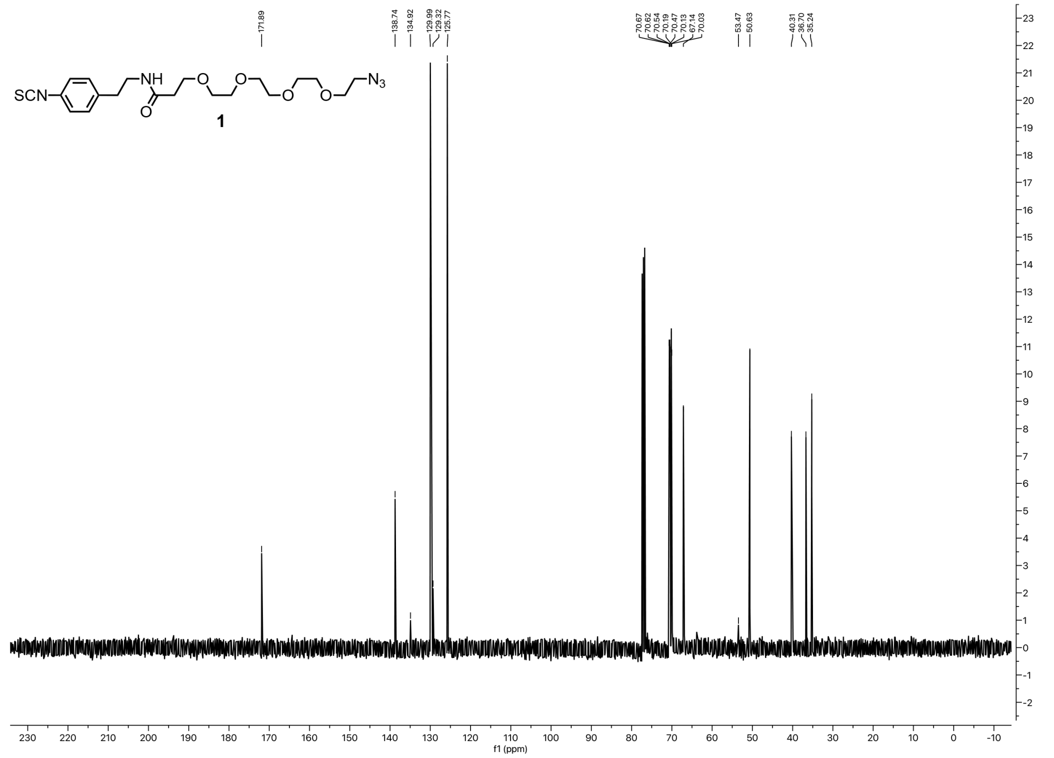

$^{13}$C NMR (101 MHz, $CDCl_3$) spectrum of **1**.

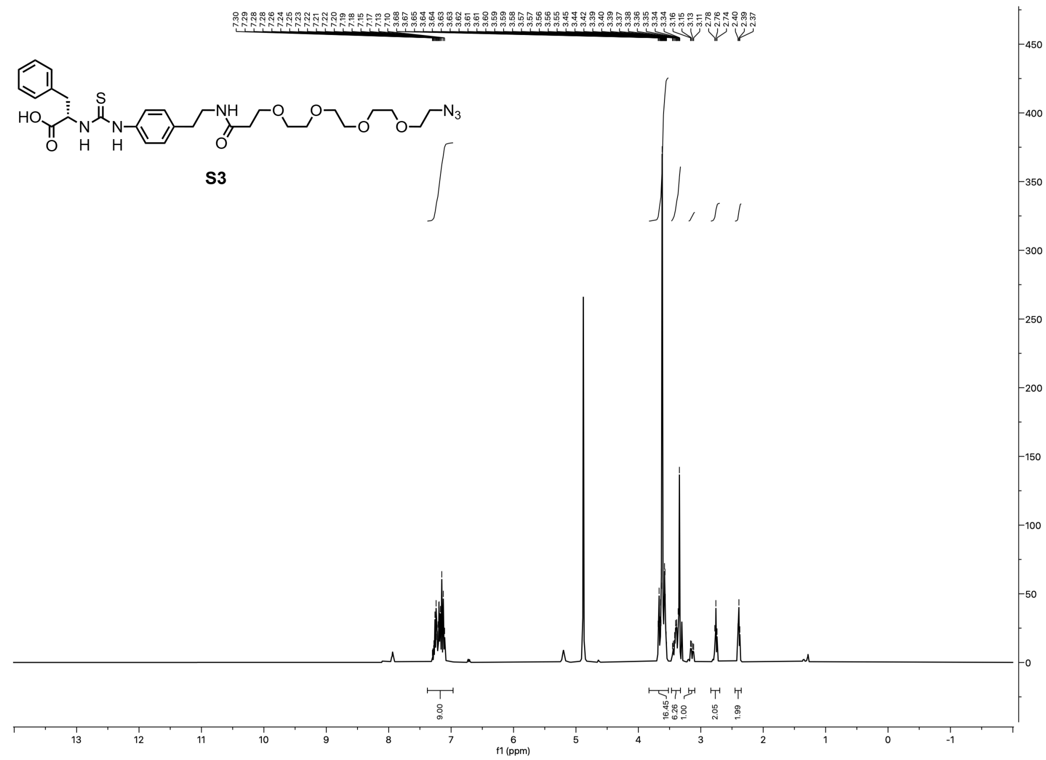

$^{1}H$ NMR (400 MHz, MeOH-d4) spectrum of **S3**.

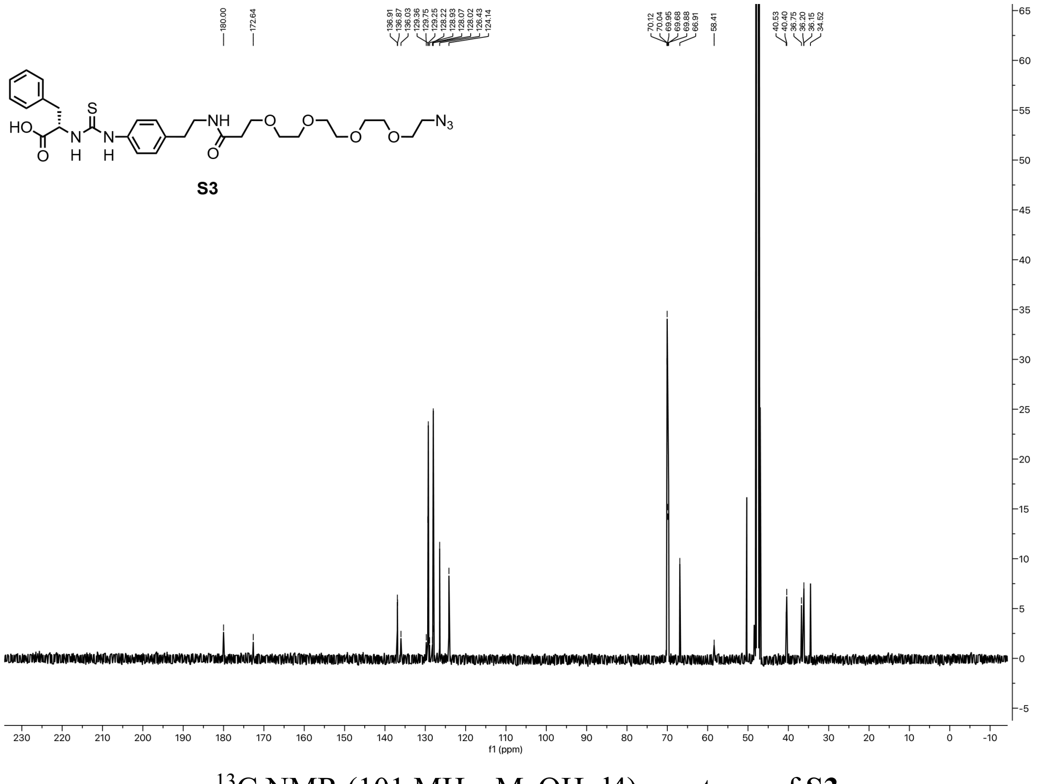

$^{13}C$ NMR (101 MHz, MeOH-d4) spectrum of **S3**.

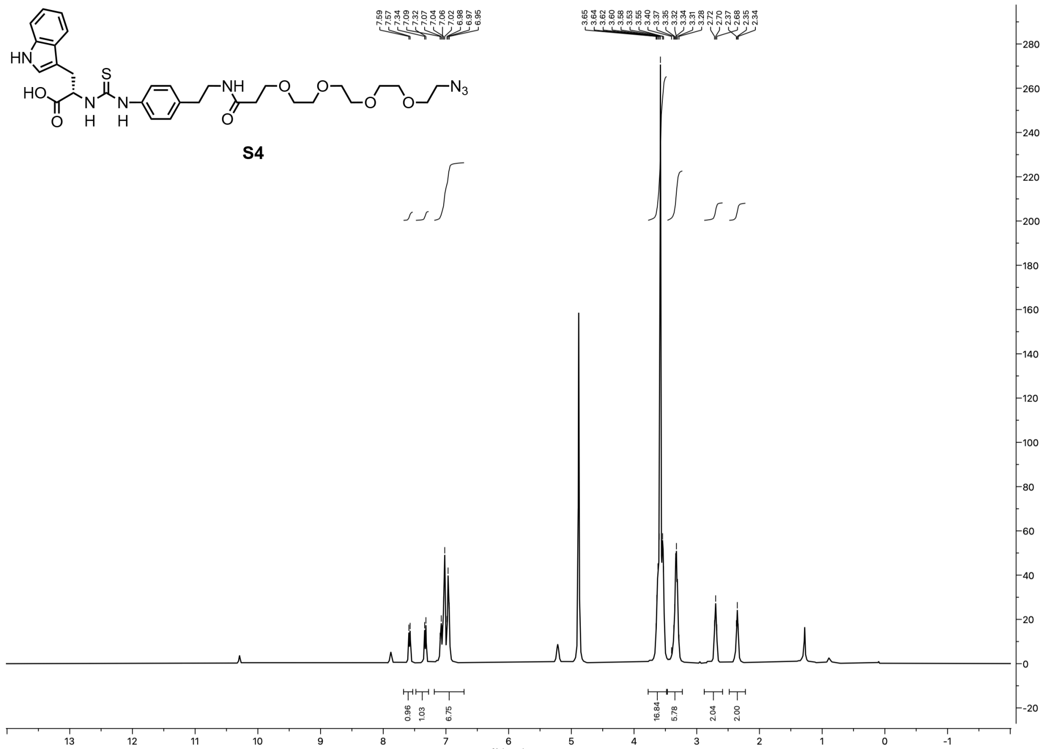

$^{1}$H NMR (400 MHz, MeOH-d4) spectrum of **S4**.

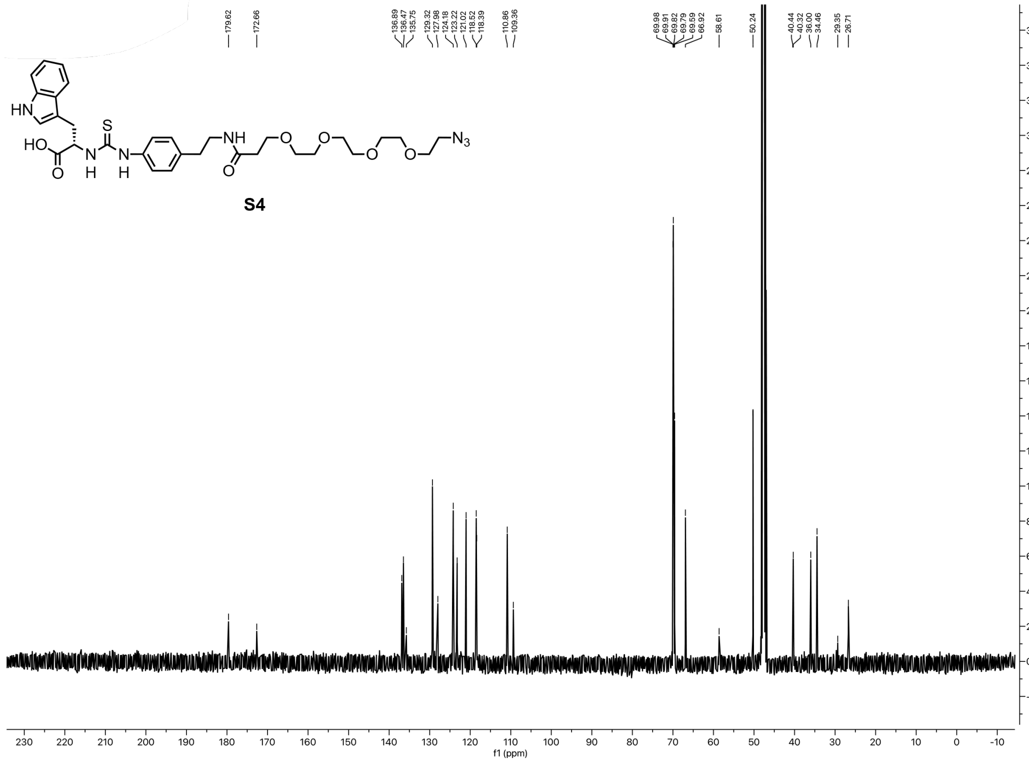

$^{13}$C NMR (101 MHz, MeOH-d4) spectrum of **S4**.

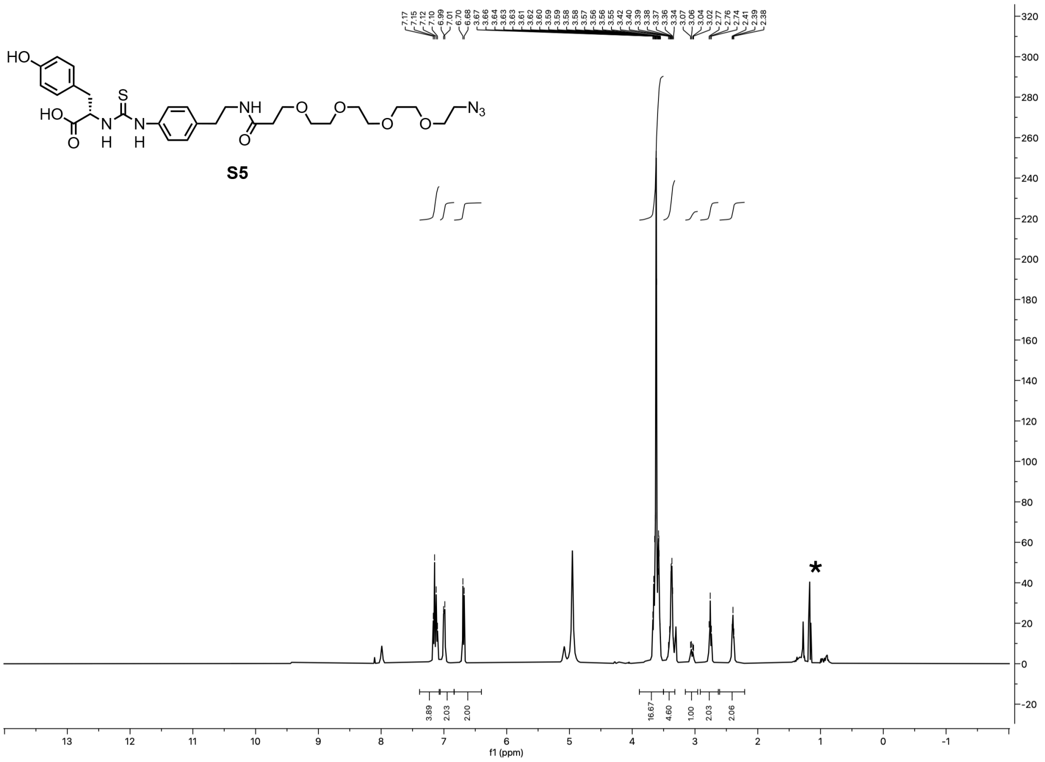

[1]H NMR (400 MHz, MeOH-d4) spectrum of **S5**. (* Impurities)

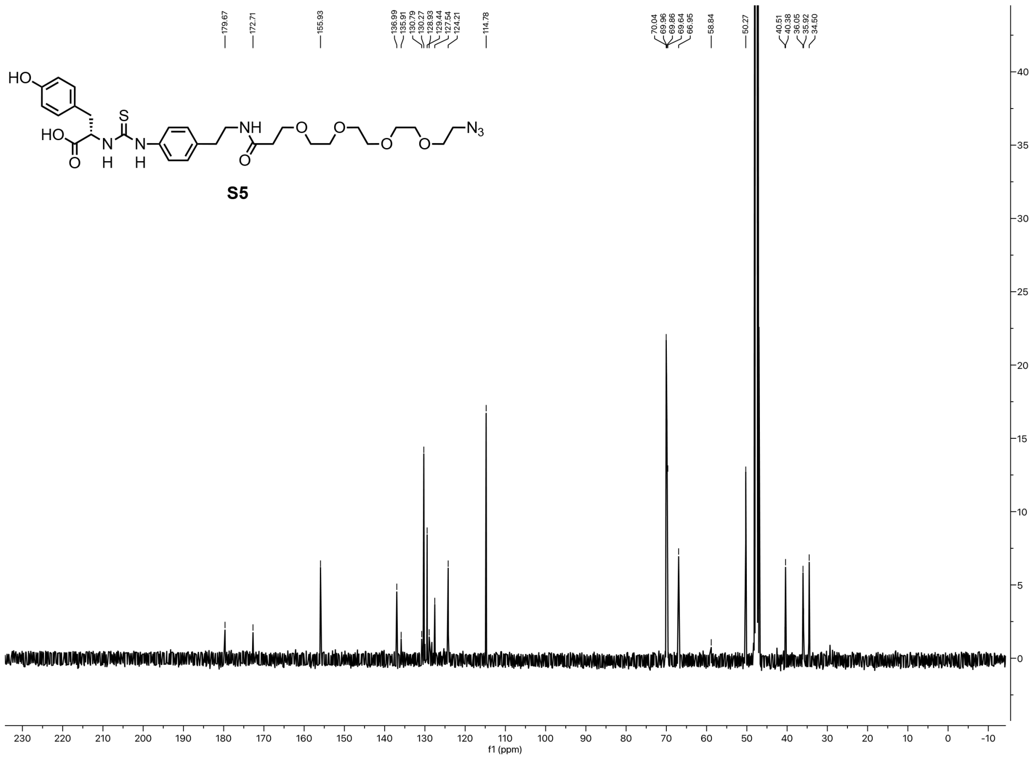

[13]C NMR (101 MHz, MeOH-d4) spectrum of **S5**.

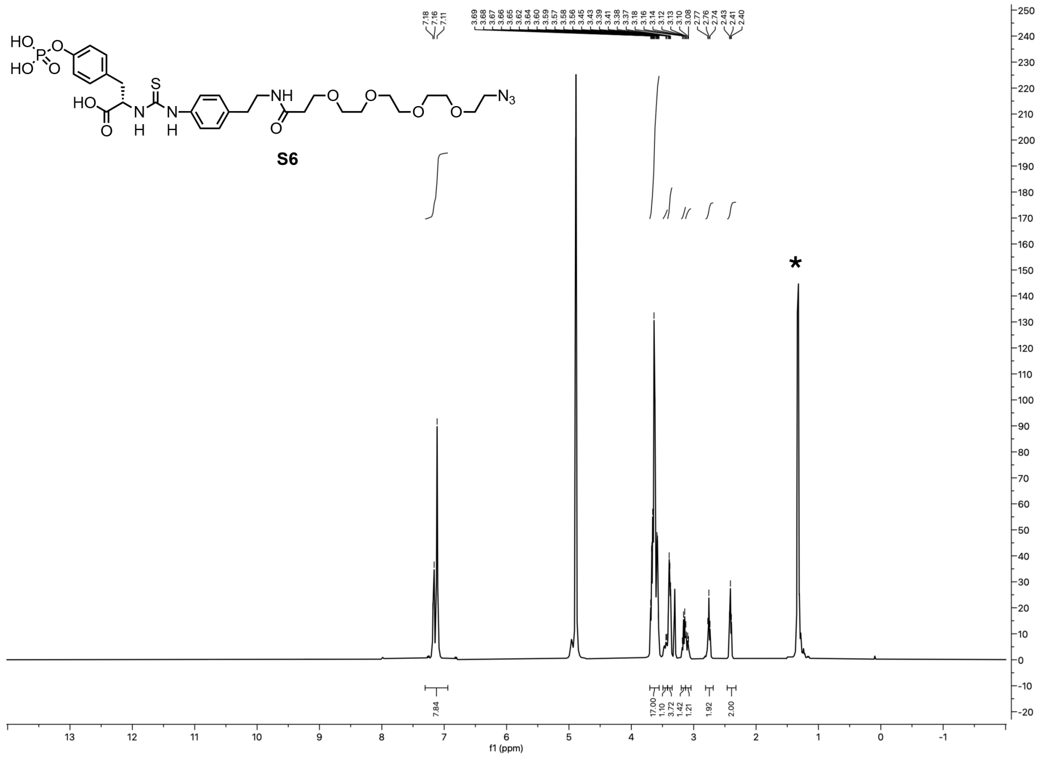


[1]H NMR (400 MHz, MeOH-d4) spectrum of **S6**. (* Impurities)

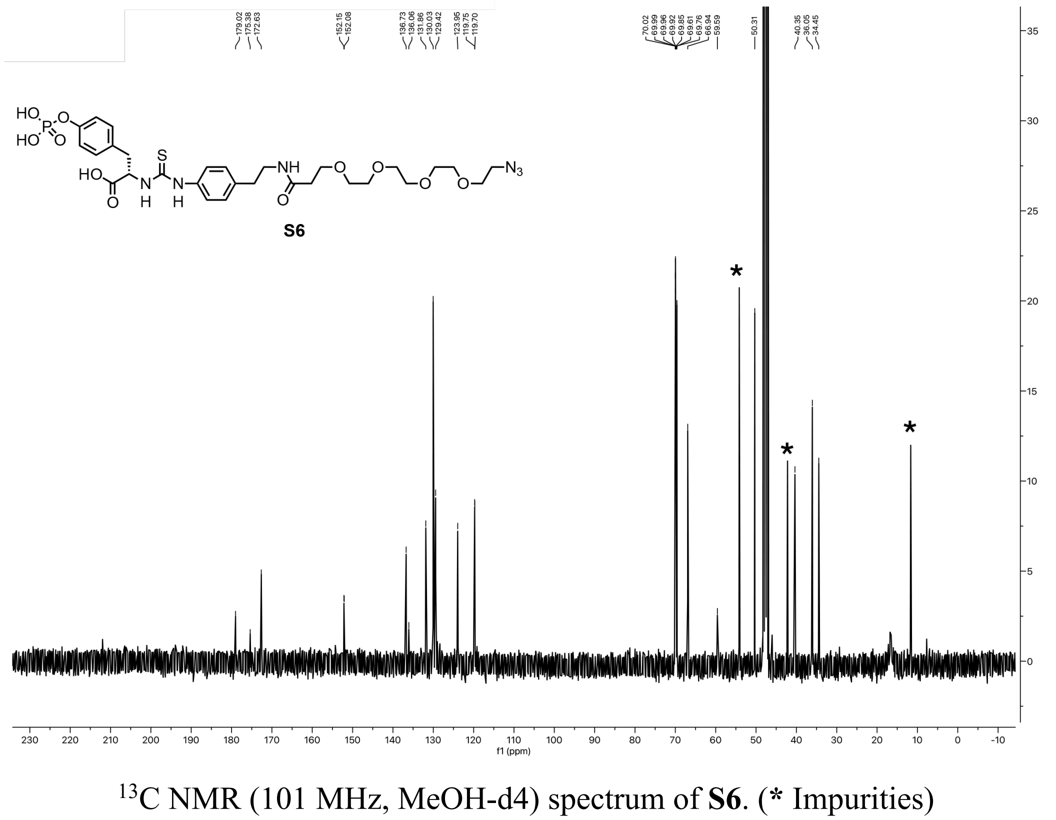


[13]C NMR (101 MHz, MeOH-d4) spectrum of **S6**. (* Impurities)

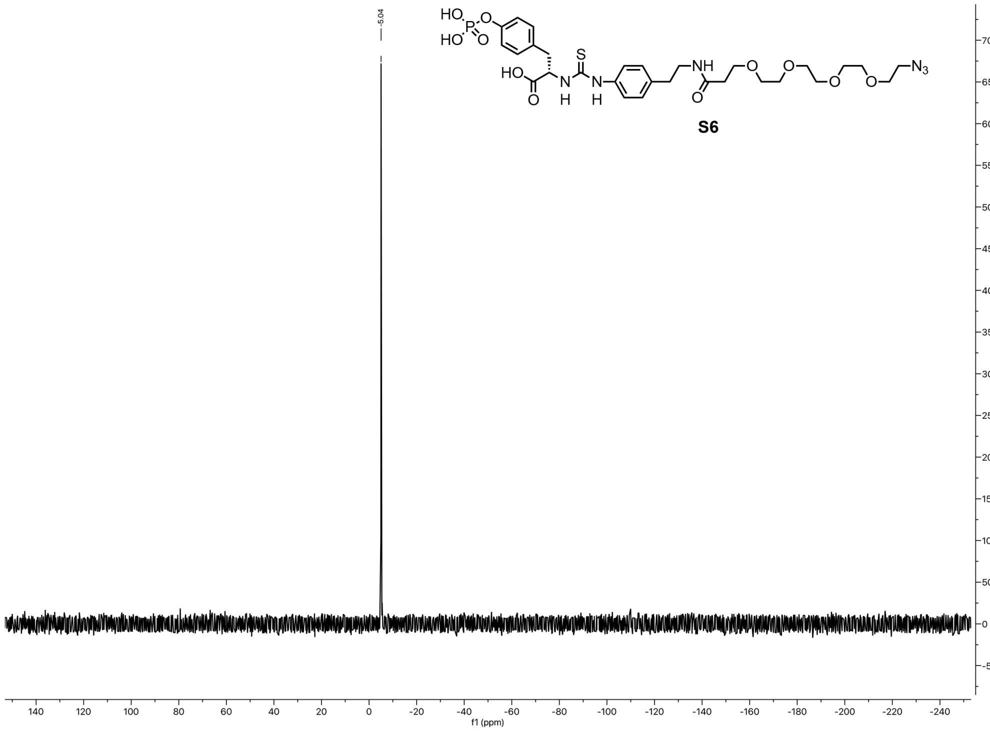

$^{31}$P NMR (162 MHz, DMSO-d6) spectrum of **S6**.

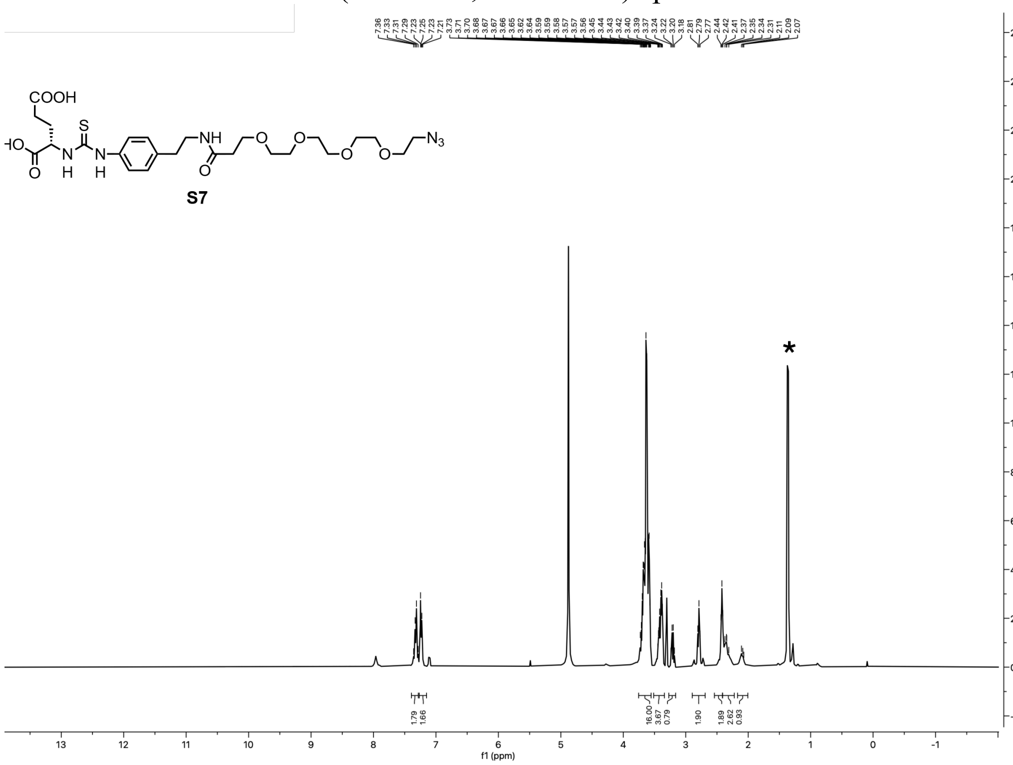

$^{1}$H NMR (400 MHz, MeOH-d4) spectrum of **S7**. (* Impurities)

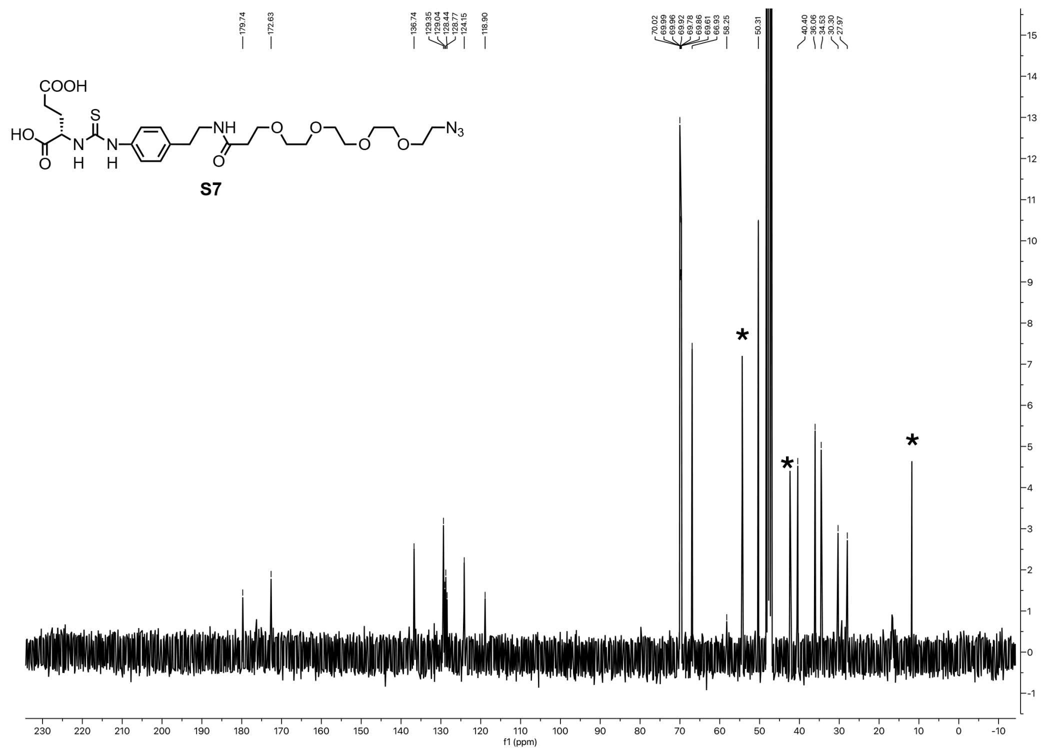

$^{13}$C NMR (101 MHz, MeOH-d4) spectrum of **S7**. (* Impurities)

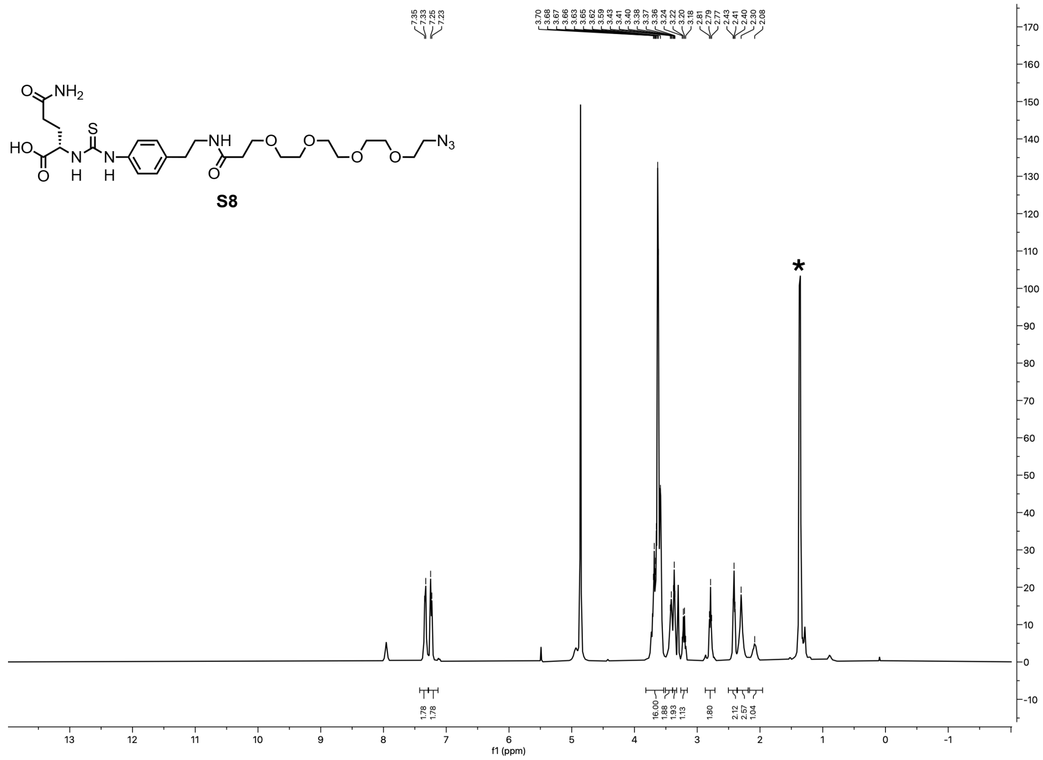

$^{1}$H NMR (400 MHz, MeOH-d4) spectrum of **S8**. (* Impurities)

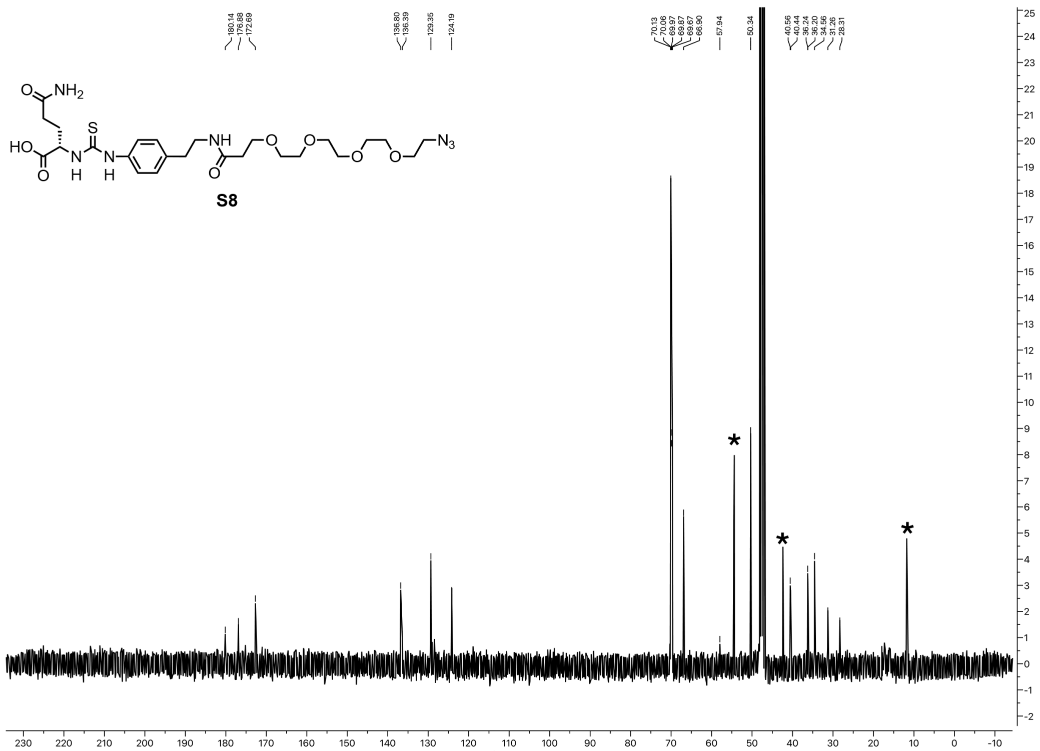

$^{13}C$ NMR (101 MHz, MeOH-d4) spectrum of **S8**. (* Impurities)

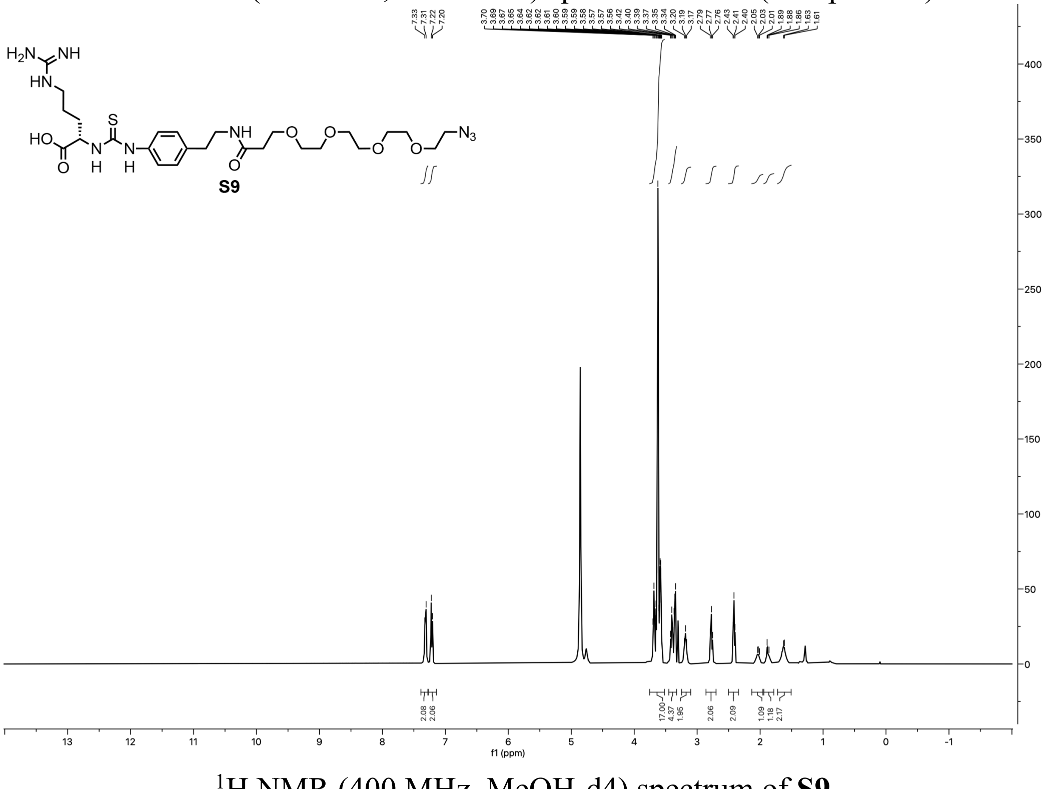

$^{1}H$ NMR (400 MHz, MeOH-d4) spectrum of **S9**.

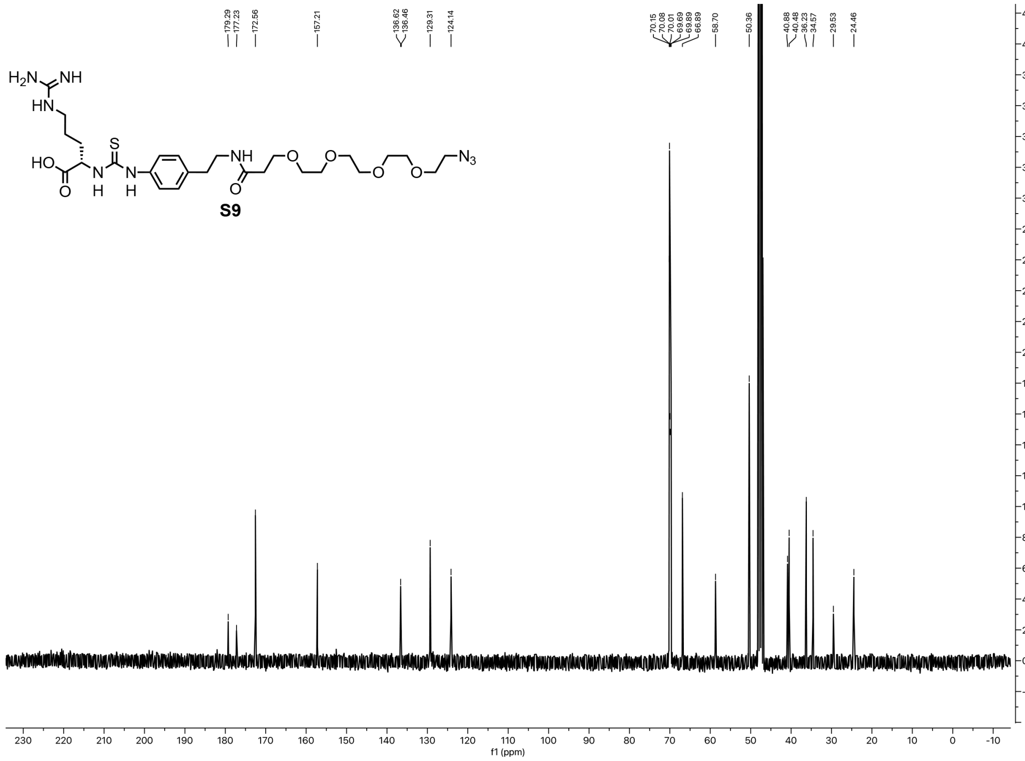

$^{13}C$ NMR (101 MHz, MeOH-d4) spectrum of **S9**.

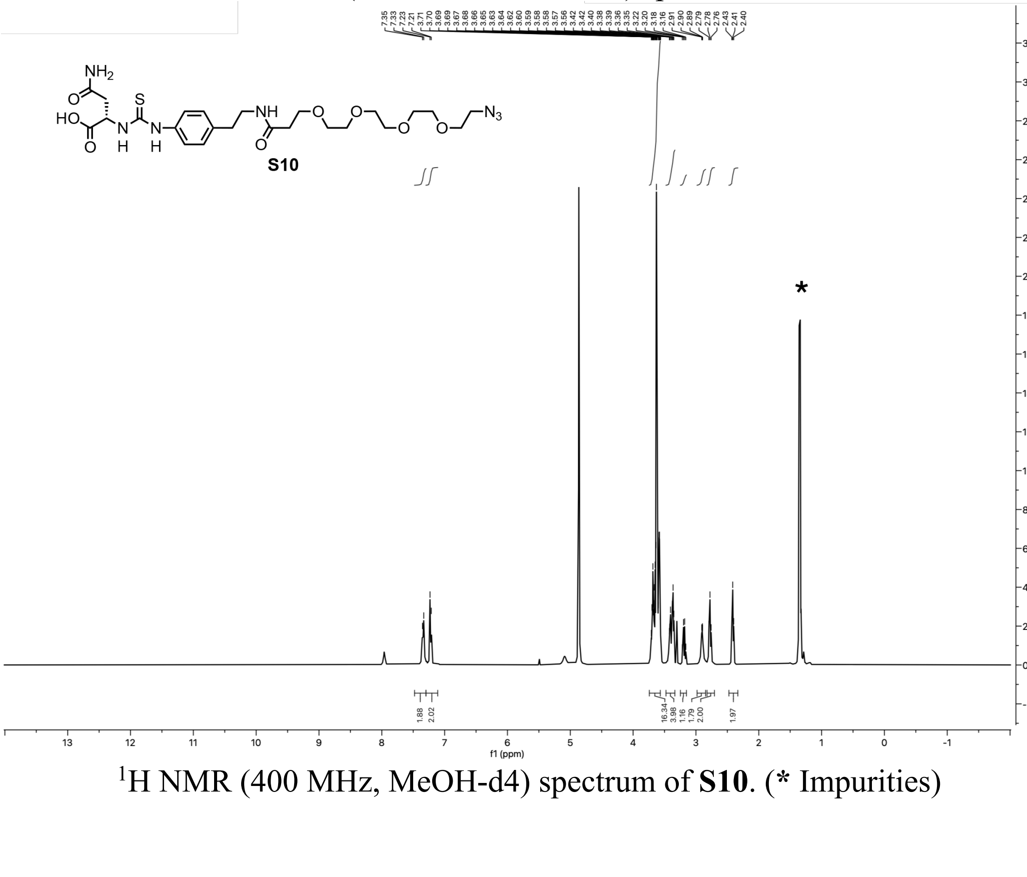

$^{1}H$ NMR (400 MHz, MeOH-d4) spectrum of **S10**. (* Impurities)

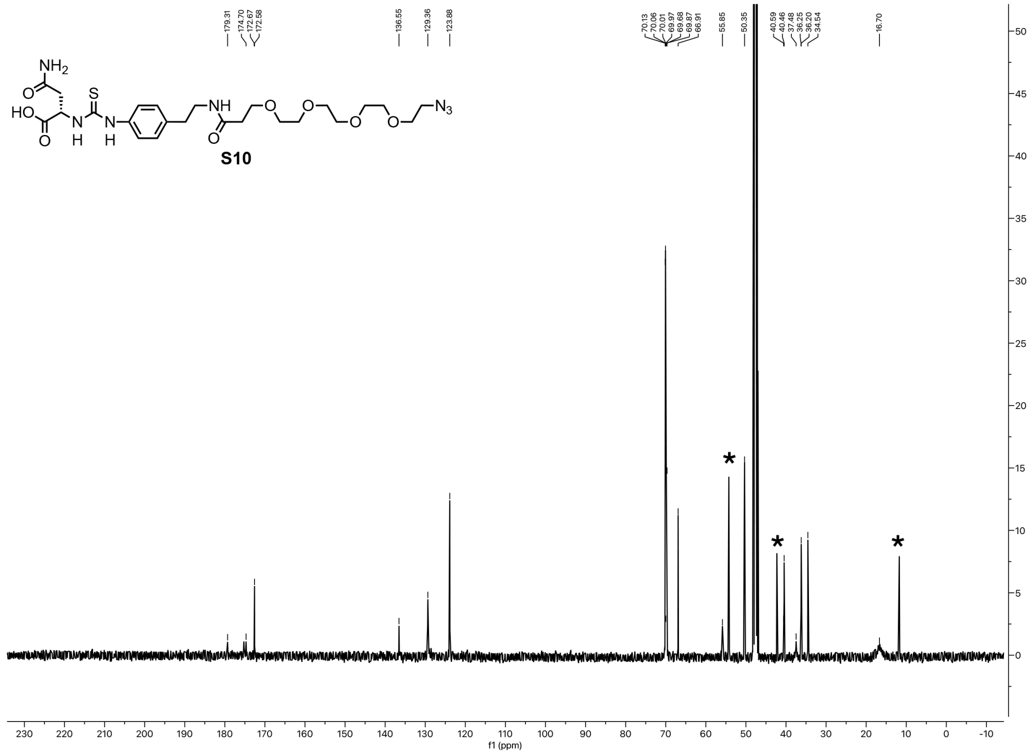

$^{13}C$ NMR (101 MHz, MeOH-d4) spectrum of **S10**. (* Impurities)

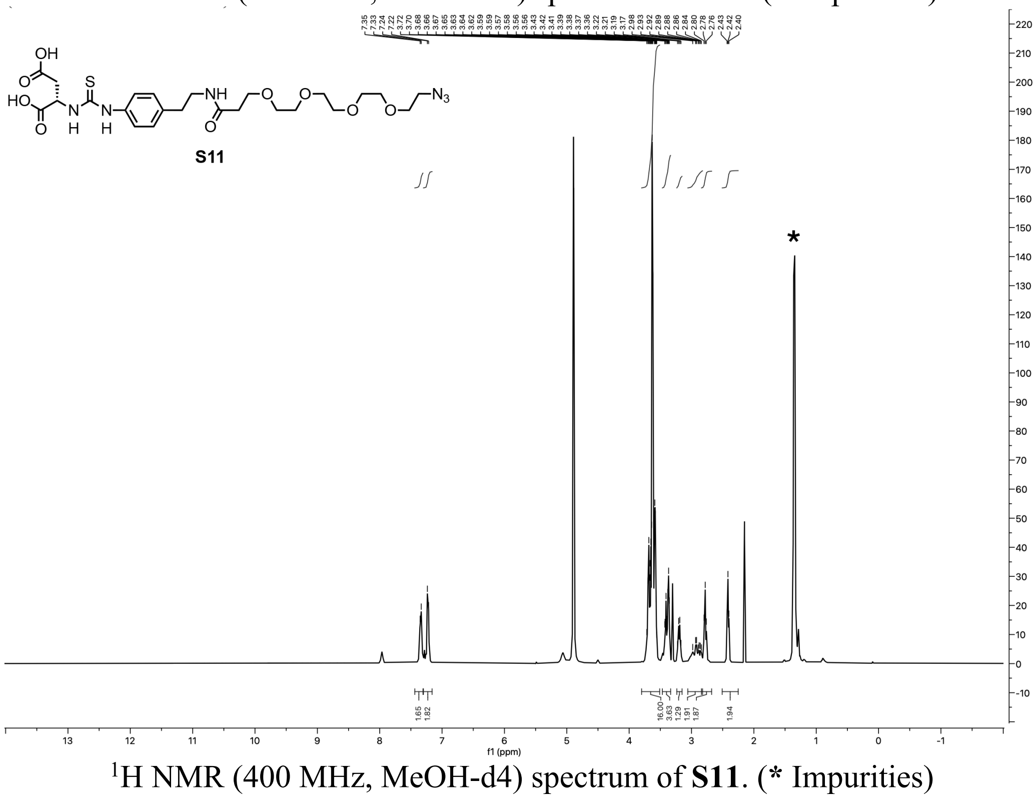

$^{1}H$ NMR (400 MHz, MeOH-d4) spectrum of **S11**. (* Impurities)

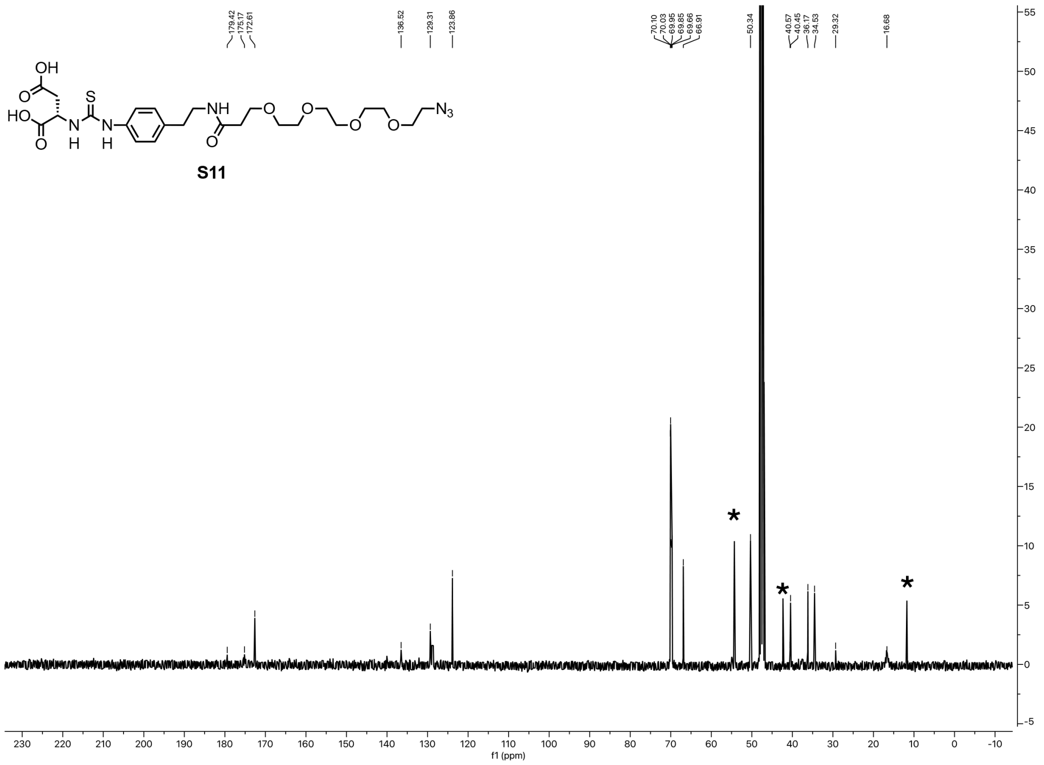

$^{13}$C NMR (101 MHz, MeOH-d4) spectrum of **S11**. (* Impurities)

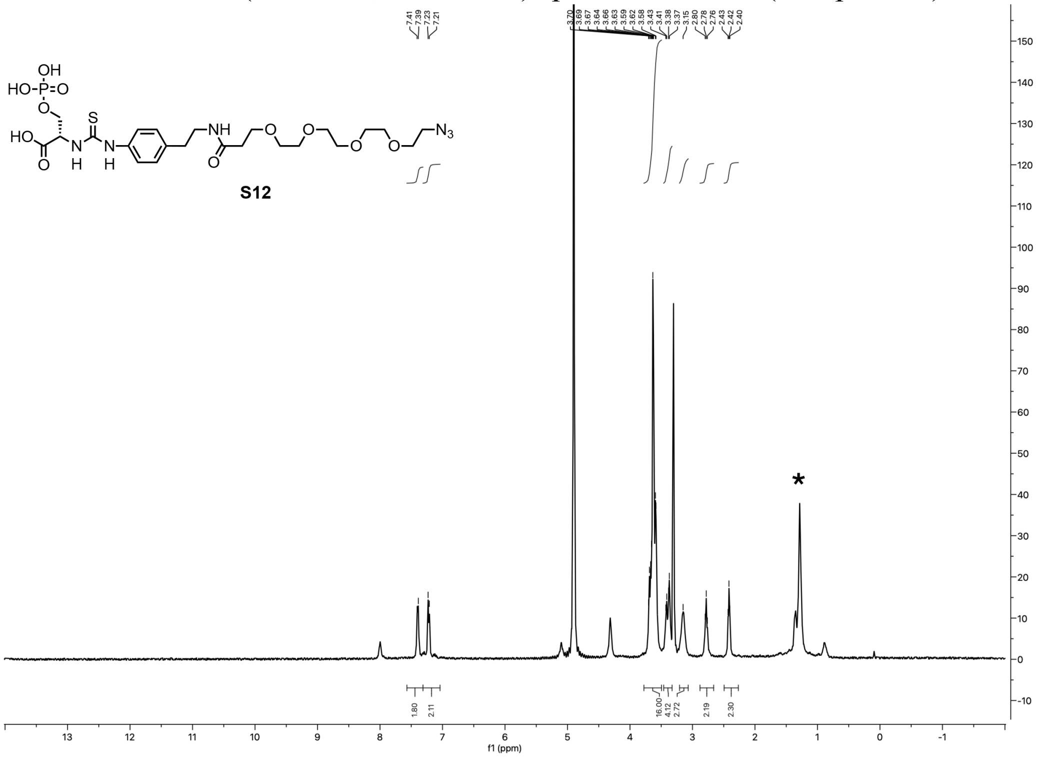

$^{1}$H NMR (400 MHz, MeOH-d4) spectrum of **S12**. (* Impurities)

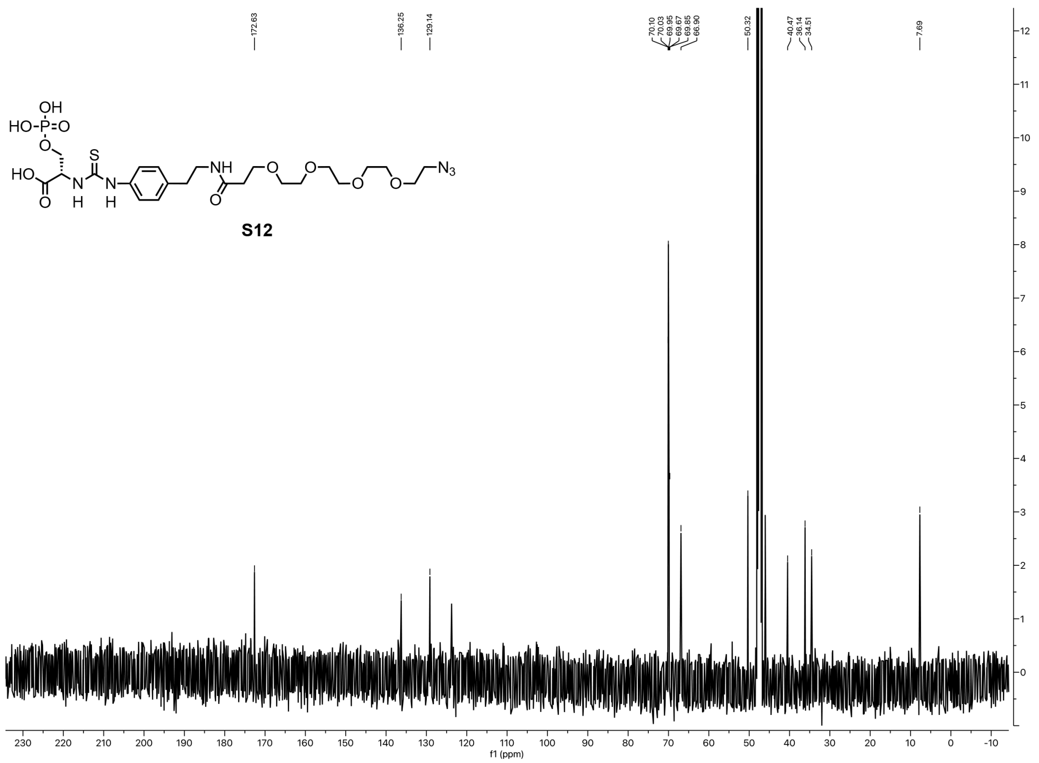


$^{13}C$ NMR (101 MHz, MeOH-d4) spectrum of **S12**.

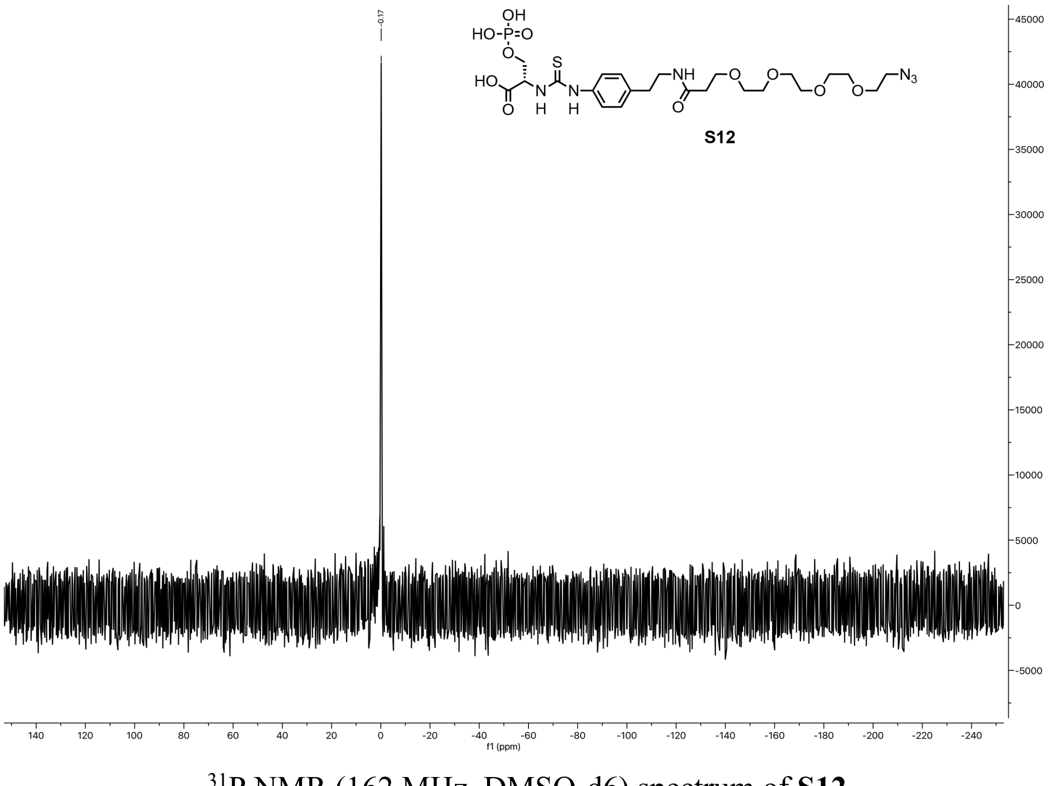


$^{31}P$ NMR (162 MHz, DMSO-d6) spectrum of **S12**.

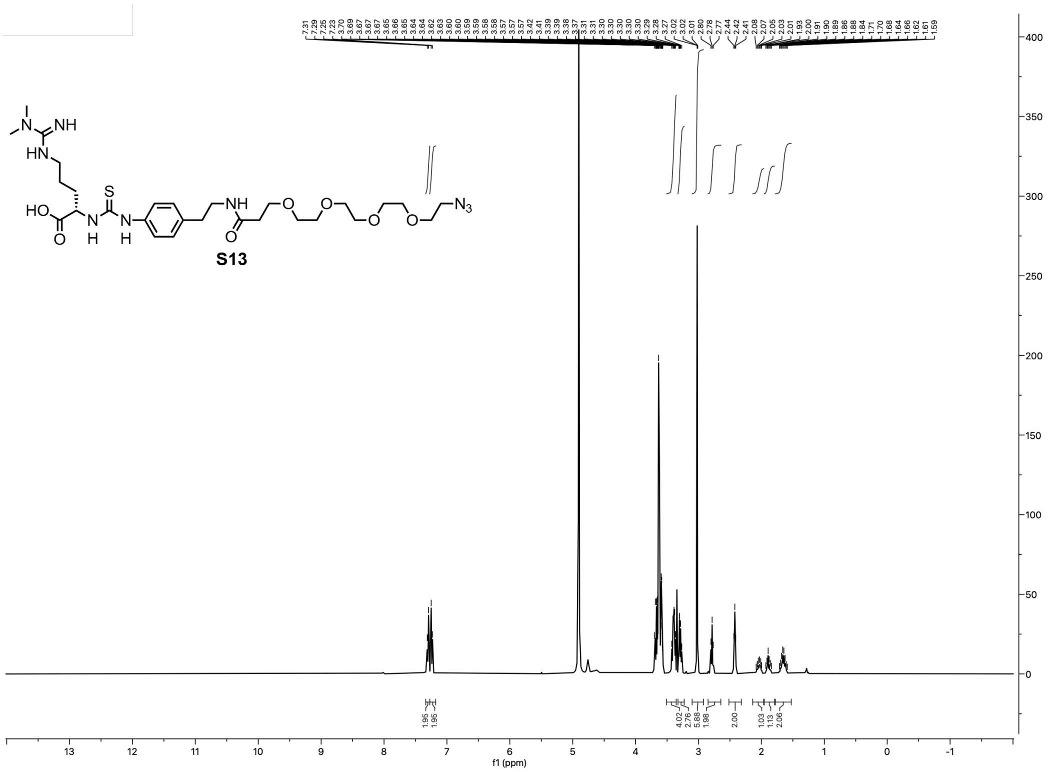


$^{1}$H NMR (400 MHz, MeOH-d4) spectrum of **S13**

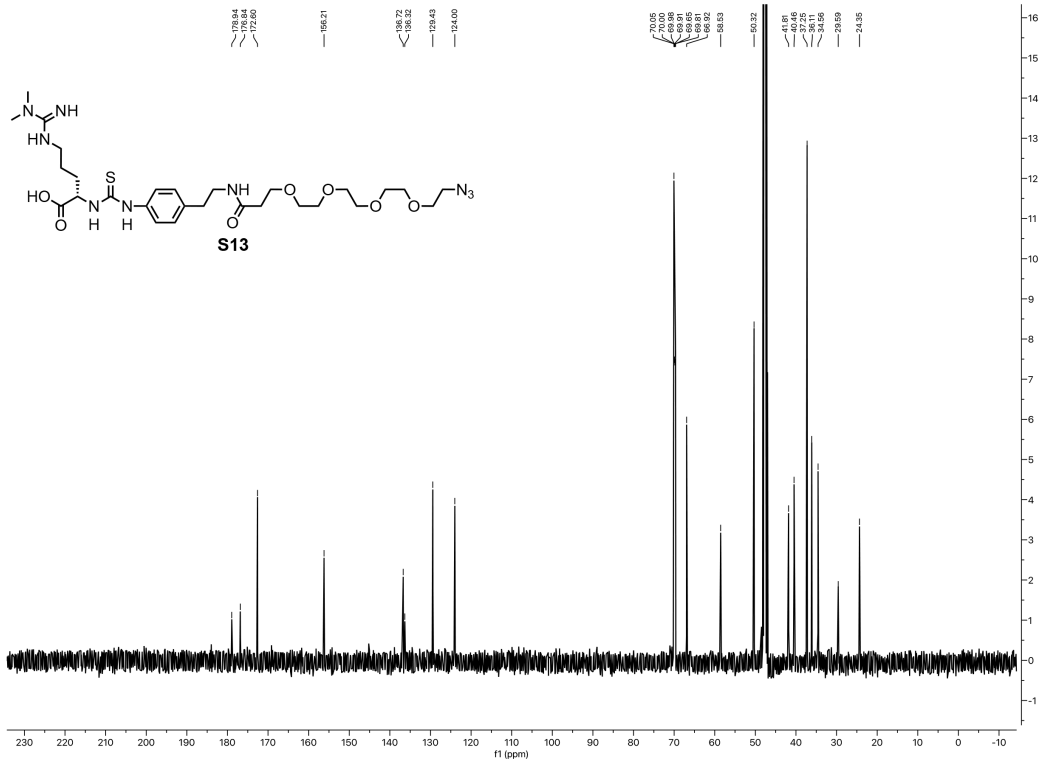


$^{13}$C NMR (101 MHz, MeOH-d4) spectrum of **S13**.

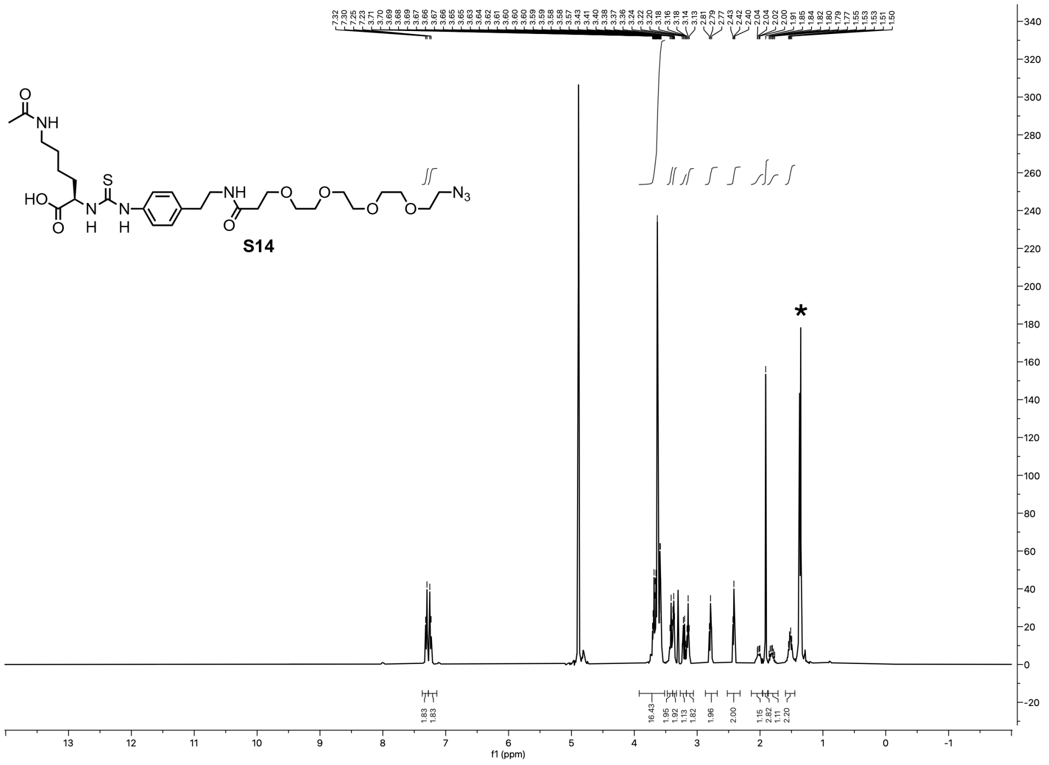

$^{1}H$ NMR (400 MHz, MeOH-d4) spectrum of **S14**. (* Impurities)

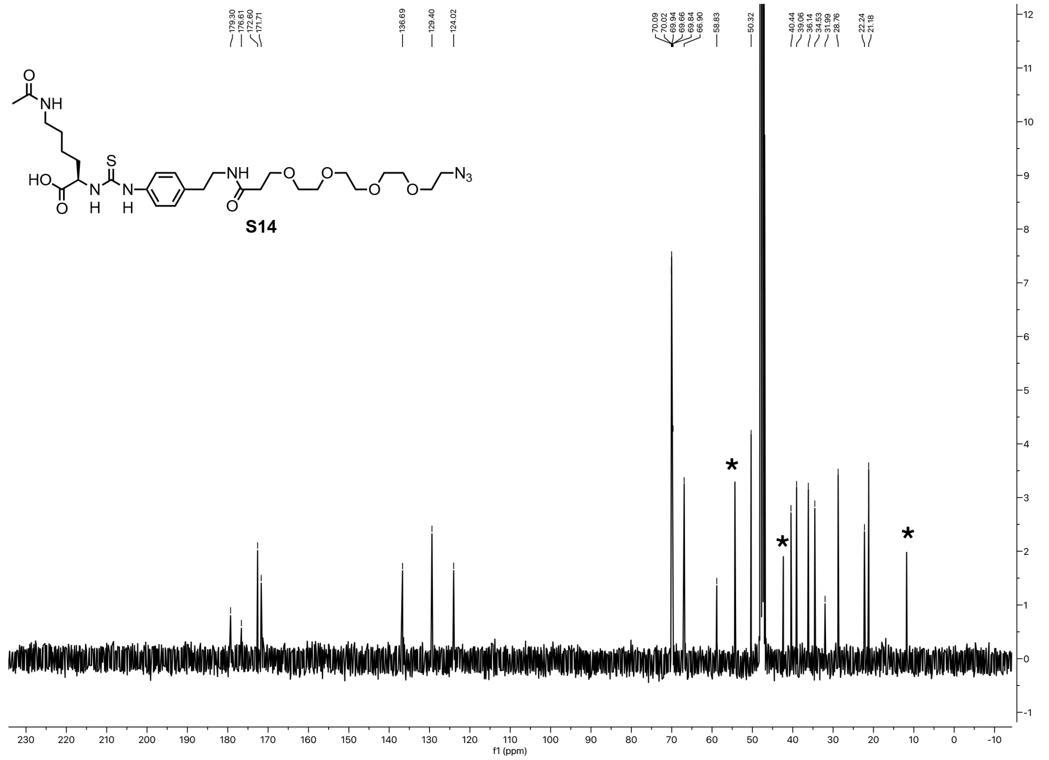

$^{13}C$ NMR (101 MHz, MeOH-d4) spectrum of **S14**. (* Impurities)
